# The Updateable Human Virome Database and toolkit: A novel framework for human virome analysis

**DOI:** 10.64898/2026.05.01.722327

**Authors:** Carson J. Miller, Christopher E. Pope, Margot H. Lavitt, Lindsay J. Caverly, John J. LiPuma, Kelsi Penewit, Janessa D. Lewis, Stephen J. Salipante, Lucas R. Hoffman

## Abstract

Current computational methods for analyzing viruses in human metagenomes rely on static databases composed largely of fragmented virus genomes mostly derived from gastrointestinal samples. This limits the identification of viruses exclusively found outside the gastrointestinal tract and impairs analyses requiring high-quality genomes. To address these issues, we created the Updateable Human Virome Database (UHVDB), an expandable database of high-quality, clustered, and annotated virus genomes integrated from pre-existing human virome databases and diverse human sample metagenome assemblies. We developed an associated toolkit that enables users to 1) update by UHVDB mining, clustering, and annotating virus sequences, 2) taxonomically profile viruses in metagenomes and metatranscriptomes, 3) estimate phage replication activity using phage-to-host ratios, and 4) identify putatively uninducible prophages from bulk metagenomes. To illustrate the utility of UHVDB and its associated toolkit, we analyzed 1,983 oral/airway samples from people with Cystic Fibrosis, finding that over 25% of viruses present were likely uninducible prophages and that many others had low replication activity, suggesting far less viral activity than estimated from previous studies. UHVDB is a novel framework for virome analysis that expands the capacity to define virus contributions to health and disease.

**GRAPHICAL ABSTRACT:** 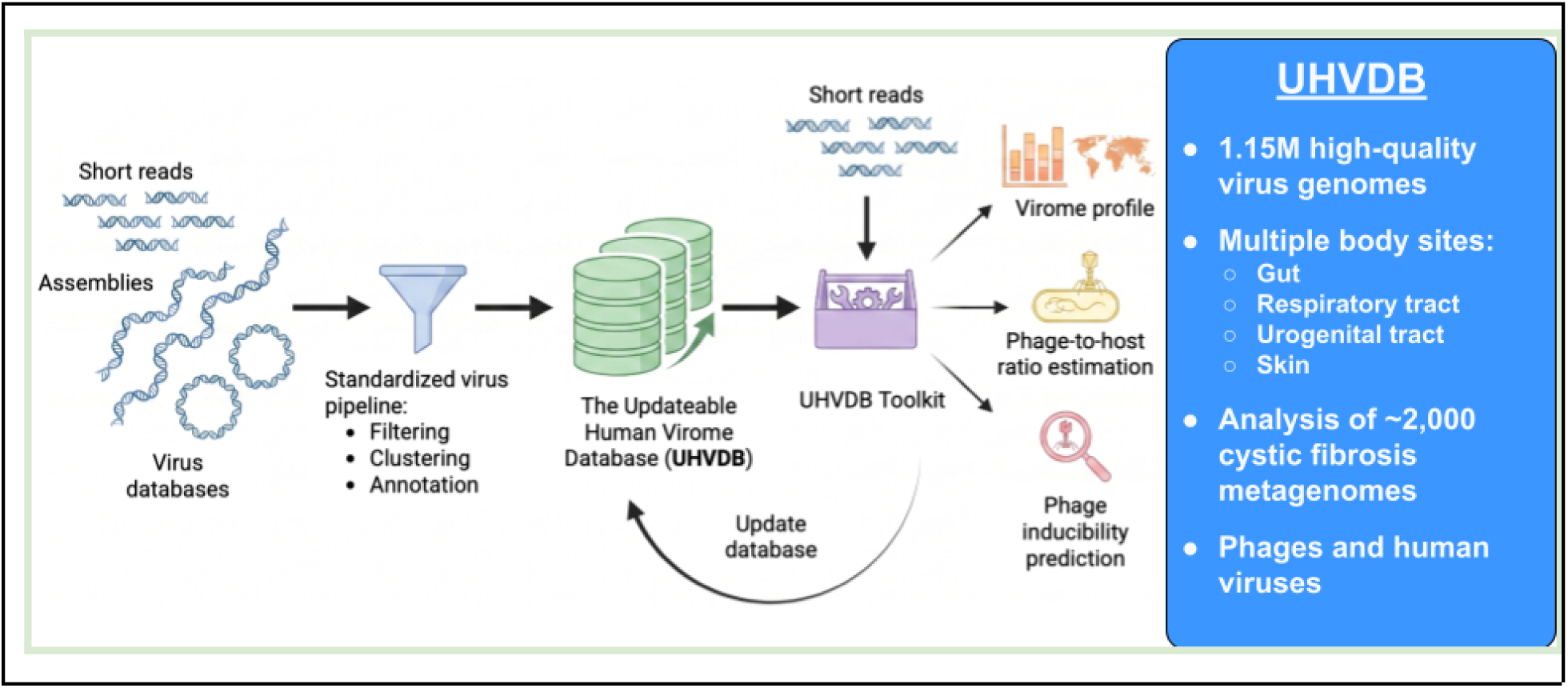

## INTRODUCTION

Viruses are the most abundant and diverse biological entities on Earth (1–3). The great majority of known viruses are bacteriophages (phages), which are viruses that infect bacteria. Common estimates suggest that there are 10^31^ phage particles on our planet, encompassing up to 1 billion unique species (1, 4). Importantly, phages are not passive entities; instead, they play important roles in shaping and regulating the biological environments they inhabit (5–7). Interest in phage has intensified as we increasingly understand both the impact of naturally-occurring phage and the potential of phage therapies in human health (8). Evidence suggests that phages can directly interact with immune cells and induce inflammation (9, 10). Phages can also indirectly affect health by altering the composition and function of bacterial microbiota through targeted lysis, horizontal gene transfer, and lysogeny (11–15). Additionally, the narrow bacterial host range of individual phages makes them promising candidates as targeted antibacterials, which is of particular interest given the global rise of antibiotic resistance (13).

Despite the significant abundance, prevalence, and health implications of phages and other viruses, their analysis using metagenomic data remains challenging. Studies utilizing large numbers of shotgun metagenomic datasets have illuminated bacterial diversity at an unprecedented scale (16–18) by leveraging conserved bacterial marker genes for sensitive genome detection and principled quality assessment (19, 20). Viruses lack universally conserved marker genes, precluding the use of similar strategies for identifying and characterizing them from metagenomic datasets. Current approaches for detecting viruses in metagenomes instead rely on assembly and the identification of virus-specific genes or nucleotide signatures, and these approaches have dramatically expanded our knowledge of virus diversity. The largest virus databases now contain millions of (predominantly phage) sequences (21–26).

Despite these important advances, extant analytical techniques for viruses carry important limitations. First, current virus databases primarily contain fragmented assemblies, each comprising a small fraction of a complete viral genome (21, 27, 28). These incomplete sequences can complicate accurate virus classification, taxonomic assignment, host prediction, and lifestyle assessment (29–32). Second, methodological variation among virus mining efforts has yielded multiple independent databases, each developed using a different approach (33–37). Camargo *et al.* recently developed the Unified Human Gut Virome Catalog (UHGV), which consolidated the largest databases of viruses identified in human fecal sample metagenomes, representing an important first step in standardizing virus identification, annotation, and clustering (38). However, this resource is restricted to one human anatomical site, necessarily omitting viruses exclusive to other human surfaces such as from the skin, oral/respiratory tract, and urogenital tract (16, 18).

Moreover, no infrastructure is available for updating UHGV with new viral sequences. Third, current analytical approaches generally do not distinguish active or potentially active phages (lytic or inducible prophages) from those that are cryptic (uninducible prophages), despite the different impacts that these viruses likely have on their environments (39–41). Many sequences identified as “phage” in metagenomic analyses may actually be uninducible prophages, given the high prevalence of these entities in bacterial genomes (41).

Here, we introduce a novel framework for consolidating and expanding human virome databases. We utilized this framework to filter virus sequences from pre-existing virus databases, mine diverse human sample metagenomes, and combine the results to create the Updateable Human Virome Database (UHVDB), substantially advancing the efforts established by UHGV. Additionally, we established phage-to-host ratios as a proxy for phage replication and developed a classifier that identifies viruses likely to be uninducible prophages. We demonstrated the practical utility of our database and its associated toolkit by analyzing respiratory metagenomes from people with cystic fibrosis (CF) to identify the diversity, characteristics, and longitudinal dynamics of viruses within that clinical context. Finally, we further expanded UHVDB by adding very recently published virus databases and RNA viruses, illustrating the utility of having an updateable database.

## MATERIAL AND METHODS

### Creating the Updateable Human Virome Database (UHVDB r1)

To create a foundation for UHVDB (release 1) and benchmark each step of database creation, high-quality viruses from UHGV (38) were run through the following steps.

Sequences were classified as viral using an updated version of UHGV’s pipeline (38). In brief, sequences were filtered to exclude those < 2,000 base pairs prior to being run through geNomad v1.11.1 (‘--relaxed’) (25). Then, sequences with a geNomad virus score ≤ 0.7 were removed unless they were classified as Inoviridae. Caudoviricetes sequences < 10,000 base pairs and Inoviridae sequences with lengths < 4,500 or > 12,500 base pairs were removed. Sequences not assigned to an International Committee on Taxonomy of Viruses (ICTV) class were also removed. Then, CheckV v1.0.3 (42) was run and sequences with high/medium confidence AAI-based completeness estimates of ≥ 50%, k-mer frequency ≤ 1.2, and length ≤ 1.5 times the expected length were retained. Remaining sequences were run through viralVerify v1.1 (43). Finally, sequences passing all prior filters were classified as “confident” (≥ 2 points), “uncertain” (0 or 1 points) or “non-viral” (< 0 points) using the following point scheme:

- +1: geNomad virus score ≥ 0.95
- +1: viralVerify viral score ≥ 15
- +1: ≥ 3 geNomad or CheckV viral hallmarks
- +1: assigned to an ICTV family by geNomad
- +1: contained a Zot gene (PF05707)
- +1: ≥ 5 Inoviridae marker genes
- −1: viralVerify “Chromosome” or “Plasmid” prediction
- −1: ≥ 2 CheckV host genes more host genes than viral genes
- −1: > 0 geNomad plasmid hallmarks
- −1: > 0 conjugation genes

After identifying “confident” and “uncertain” viruses using the protocol described above, we expanded CheckV’s database of direct terminal repeat (DTR) genomes as in UHGV. First, we identified UHGV’s DTR-containing species representatives and aligned these to CheckV’s existing database using vClust v1.3.1 (44). Sequences with < 95% average nucleotide identity (ANI) or < 85% alignment fraction (AF) to all CheckV sequences were then added to CheckV’s database using the ‘update_database’ command. Then, we identified other DTR-containing confident and uncertain viruses with ≥ 80% AAI-based completeness output by our virus classification workflow. These were dereplicated at the vOTU level (45) with vClust (95% ANI and 85% AF), then novel sequences were identified and added to CheckV’s database.0

Using our updated version of CheckV’s database, we re-calculated the AAI-based completeness for each of the confident and uncertain viruses, and retained only the high-quality (≥ 90% AAI-completeness) sequences.

High-quality viruses classified as “uncertain” were further investigated by running HMMER v3.4 (‘-E 1e-5’) (46) against geNomad’s (db v1.9) database of virus and plasmid hallmarks. Uncertain virus sequences with < 3 virus hallmarks or > 0 plasmid hallmarks were removed from the dataset (Supplementary Fig. S4).

All high-quality, confident sequences with DTRs had terminal repeats removed from the right side using tr-trimmer v0.4.0 (47). Then, the sequence hashes were calculated using seq-hasher v0.1.0 (48). Unique sequences were identified using unique sequence hashes.

### Clustering UHVDB

After creating UHVDB r1 as described above, virus sequences were run through the following clustering steps.

Unique sequences were grouped into genomovars (48, 49) by performing all-v-all alignment with vClust v1.3.1 (44) and clustering global ANI (ANI x AF) values from genome pairs with ≥ 99.5% ANI and 100% AF relative to the shorter sequence using MCL v22.282 (50) to yield genomovars (Supplementary Fig. S5).

Representatives for each genomovar cluster were selected using a modified version of UHGV’s species representative selection protocol. First, singletons were automatically set as the representative. Second, the longest sequences containing DTRs were assigned as representatives. Third, linear genomes having the most CheckV viral genes were chosen, with a tiebreaker being the genome whose length was closest to CheckV’s expected length.

To cluster UHVDB r1 at the species level, we performed an all-v-all alignment of genomovar representatives using vClust and retained genome alignments with ≥ 95% ANI and ≥ 85% AF relative to the shortest sequence (45). Then, the global ANI values were clustered using MCL.

UHVDB r1 species representatives were selected with the same approach used for genomovar representatives and then clustered at the family, subfamily, genus, and subgenus levels using an adapted version of UHGV’s hierarchical clustering protocol. First, protein sequences were predicted by running pyrodigal-gv v0.3.2 (25, 51, 52). Then, proteins were all-v-all aligned using Diamond v2.1.12 (‘--very-sensitive -k 0 -e 1e-3’) (53).

The proteomic similarity for each genome comparison was calculated by summing the bitscore of all alignments between a genome pair, and dividing by the summed bitscores of a self alignment.

After calculating proteomic similarities, the alignment graph was filtered to remove genome comparisons with < 5.5% protein similarity and run through MCL v22.282 (50) to create families. Then, the graph was filtered to retain only intra-family genomes with ≥ 32% protein similarity and clustered with MCL to yield subfamilies. Intra-subfamily genomes with ≥ 65% protein similarity were then clustered into genera. Finally, intra-genus genomes with ≥ 80% protein similarity were clustered into subgenera.

### Annotating UHVDB

Although UHGV taxonomy, host, and lifestyle annotations were often consistent at the genus level and below, we opted to annotate all UHVDB genomovar representatives because protein content can vary substantially at the species level (Supplementary Fig. S7).

To assign taxonomy to UHVDB r1 genomovar representatives, we first used geNomad’s taxonomy. To assign taxonomy at lower ranks (family and genus), we calculated the proteomic similarity between each genomovar representative and all complete ICTV genomes (VMR MSL40). UHVDB genomes with family, subfamily, genus, subgenus, or species-level proteomic similarities to an ICTV reference (family: 5.5%, subfamily: 32%, genus: 65%, subgenus: 80%, species: 95%) were assigned the reference’s taxonomy at the associated taxonomic rank.

To predict the bacterial hosts for genomovar representatives, we updated the approaches set forward by UHGV. First, we identified bacterial MAGs from human-associated metagenomes in mOTUs-db (n=825,354) (18). Then we added mOTUs-db isolate assemblies belonging to the same genus as human-associated MAGs (n=1,001,750). These 1,827,104 genomes were then searched against all genomovar representatives using PHIST (54). All bacteria-virus pairs where ≥ 20% of virus k-mers were contained in the bacterial assembly were retained. Host taxonomy was assigned at the lowest rank having ≥ 70% taxonomic agreement among at least 1 host connections.

The second method for identifying bacterial hosts was CRISPR spacer matching. We aligned all 24,639,433 spacers created for VIRE (55) to UHVDB genomovar representatives using SpacerExtractor (56). All phage-bacteria pairs with ≤ 1 mismatch over the entire length of the spacer were retained. Host taxonomy was assigned at the lowest rank having ≥ 70% taxonomic agreement among at least 1 host connections.

To assign a final host, we used the prediction method having the most host connections for each virus, or the CRISPR prediction when both methods were equal. Hosts for genomovars in a eukaryotic virus class (Arfiviricetes, Cardeaviricetes, Papovaviricetes, Quintoviricetes, and Repensiviricetes) were labeled as “Eukaryote”.

To functionally annotate UHVDB, we predicted protein protein sequences among UHVDB r’s genomovar representatives using pyrodigal-gv (25) and calculated sequence hashes using seqkit v2.12.0 (57) to identify unique proteins. Then, we assigned searched each unique protein against UniProt/InterPro (58, 59) with Bakta v1.12 (60) supplemented with viral UniRef50 proteins from March 2026 (61), foldseek v10.941cd33 (62) (‘-e 0.001 -c 0.9’) against BFVD (63), VAD (64), and VFOLD (65), and InterProScan v5.74-105.0 (66).

Our second approach for functional annotation was searching all unique proteins against phage-specific databases. To do this, we used Pharokka v1.9.1 (‘pharokka_proteins.py’) (67), Phold v1.2.2 (‘proteins-predict’ and ‘proteins-compare’ subcommands) (68), and Empathi v1.0.6 (‘--confident 0.5’) (69).

Our third approach was to use specialized tools/databases to identify functions of interest. In brief, we used DIAMOND v2.1.18 (‘ --very-sensitive --iterate --max-target-seqs 1 --id 80 --query-cover 40 --subject-cover 40‘) (70) to align proteins to the Comprehensive Antibiotic Resistance Database (CARD) v4.0.1 (71) and the Virulence Factor Database (VFDB) (72) from March 2026.

To predict the lifestyle of UHVDB r1 genomovars, we investigated the following temperate signals: geNomad or CheckV integration, a BACPHLIP v0.9.6 (73) temperate score > 0.5, and the presence of an integration or excision-related gene detected by Pharokka, Phold, or Empathi. We opted to classify genomovars with ≥ 1 signal as temperate.

### Updating UHVDB

To expand UHVDB, additional virus database sequences and metagenome assemblies were run through the classification, clustering, and annotation workflows described above.

The first expansion (r2) involved adding uncertain viruses from UHGV. For the next expansion (r3), virus sequences were downloaded from 9 different sources (21, 28, 74–80).

These databases, with the exception of IMG/VR4 (21), were composed entirely of human-related viruses, so every sequence was mined. For IMG/VR4, sequences were filtered to retain only those containing “Human” in the “Ecosystem Classification” field. For the fourth release (r4), metagenome assemblies from ENA (81), Logan (82), and SPIRE (16), representing samples from the human airways, urogenital tract, and skin, were identified and mined to expand UHVDB.

### Evaluating UHVDB’s comprehensiveness

To assess UHVDB r4’s comprehensiveness, we used two independent approaches. First, we created accumulation curves at the species, subgenus, genus, subfamily, and family levels (Figure 2A-C; Supplementary Fig. S12). These were created by splitting unique viruses by body site of origin, randomly subsampling each body site at 100 increasing intervals, and counting the number of different clusters present at each interval. This process was repeated 50 times and the mean count of clusters detected at each interval across replicates was plotted.

Additionally, we assessed how well UHVDB represents newly assembled viruses relative to other human virome databases. To do this, we assembled 400 metagenomes not mined in creating UHVDB from the gut, oral/respiratory tract, skin, and urogenital tract. High-quality, confident viruses were identified using the workflows described above. Then, these were aligned to Inphared (83), UHGV (38), IMG/VR4 (21), and UHVDB r4 using BLAST (84) and the anicalc scripts from CheckV. The highest global ANI (ANI x AF) value for each virus was identified and plotted (Figure 2D, Supplementary Fig. S12).

### Creating the UHVDB toolkit

To create the UHVDB toolkit pipeline, we first implemented the scripts used to create and update UHVDB as a ‘--run_update’ workflow. This Nextflow (85) workflow was formatted using the structure established by the nf-core community (86). To enable broad usage, the UHVDB toolkit is compatible with an infrastructure that can run conda/mamba, Singularity/Apptainer, or Docker.

To create a virome profiling workflow as a part of the UHVDB toolkit, we developed a ‘--run_analyze’ workflow that runs sylph (87) against UHVDB r4 species representatives (‘-c 200’) and GTDB r226 species representatives (‘-c 1000’) simultaneously (88). UHVDB species detected with sylph are then input to CoverM (89) for read alignment and breadth of coverage calculations.

### Benchmarking virome profilers

To benchmark virus taxonomic profilers, we used the high-quality, confident viruses that were assembled for the comprehensivity analysis described above. These viruses were dereplicated at the species level (95% ANI and 85% AF) and split into 17 sets of 100 viruses, each set containing viruses exclusively from one body site. For each set, 100,000 reads were simulated using InSilicoSeq (90). Reads were profiled using BAQLaVa (91), Phanta (92), Marker-MAGu (93), and the UHVDB toolkit (87) (sylph using UHVDB species representatives). The assembled viruses were aligned to each tool’s database of virus sequences using BLAST, and reciprocal-best-hits were determined using global ANI. For each body site, the recall (number of detected viruses divided by the total number of input viruses) and Pearson R values were calculated to evaluate performance (Figure 4A, Supplementary Fig. S13). To evaluate whether the performance was simply an artifact of using a different profiling method (sylph) we ran sylph using each tool’s database of viral genomes (Supplementary Fig. S13).

### Evaluating phage-to-host (PTH) ratios

To quality check UHVDB’s host predictions, we analyzed a dataset of fecal shotgun metagenomes (94) and calculated co-occurrence and Spearman correlations for phage-bacteria paired using UHVDB’s predicted host species, random phage-bacteria pairs, and pairs using a non-host species within UHVDB’s predicted host genus (Supplementary Fig. S14). Then, we calculated phage-to-host (PTH) ratios by dividing phage depth of read coverage by the bacterial host species depth of coverage (Supplementary Fig. S14).

To evaluate our calculated PTH ratios, we compared them to PTH values derived from running PropagAtE (95) on medium-quality or better proviruses identified using megahit, geNomad, and CheckV on the same samples. The biological utility of PTH ratios was investigated by running the UHVDB toolkit and PropagAtE on shotgun metagenomes from 10 bacterial isolates sequenced with and without Mitomycin C (MMC), a common prophage inducing agent (96) and compared the detection of induction (PTH in MMC > 2 times PTH in no MMC). As another test, we ran a dataset of highly enriched fecal viromes (ViromeQC score ≥ 50) and their paired unenriched (bulk) metagenomes through the UHVDB toolkit (94) and determined the proportion of bulk viruses that were detected in both bulk and enriched samples based on increasing minimum PTH thresholds (Supplementary Fig. S15).

### Creating a Random Forest Classifier to identify uninducible prophages

Highly enriched fecal viromes (94) along with their paired unenriched (bulk) samples, were downloaded, assembled, and run through the UHVDB toolkit’s analysis workflow. For each pair of samples, a virus was considered an uninducible prophage if it was detected only in the bulk sample. The precision (proportion of viruses passing a threshold that were only detected in the bulk) and recall (proportion of all viruses only detected in bulk) were calculated across different features (Supplementary Table S1) using a variety of thresholds (Supplementary Fig. S16) to determine their utility in distinguishing active from uninducible prophages.

Next, we trained a class-balanced random forest classifier (100 trees; Scikit-learn v1.5.1 (97)) with isotonic calibration using 20 features which were selected in an effort to detect genome truncation, the presence of direct terminal repeats (DTRs), spurious virus detections, viral confidence, virus integration, and coverage of structural virus genes. The training dataset consisted of the previously mentioned paired bulk and enriched fecal samples, with true positives (TPs) being UHVDB species detected in only the bulk samples, and false positives (FPs) being virus species detected only in the bulk sample. The classifier utilized 100 decision trees and a balanced class-weighting strategy was employed due to the inequality of TPs (2,532) and FPs (928). Performance was evaluated using group 5-fold cross-validation, grouping by sample of origin, and the Area Under the Precision-Recall curve (AUPRC) values were calculated (Figure 4D). A classification score ≥ 0.87 was determined to correlate with 95% precision in our cross-validation and was used in all downstream classifier applications.

To evaluate this classifier, we identified another dataset of 6 paired bulk and highly enriched fecal viromes (98) and processed this dataset using the UHVDB toolkit. Bulk viruses having a classifier confidence score ≥ 0.87 were labeled as uninducible prophages. These labels were compared to whether or not a virus species was also detected in the paired enriched sample to calculate precision and recall (Supplementary Fig. S17). Our classifier was also benchmarked by running the UHVDB toolkit on a dataset of bacterial isolates having paired no-Mitomycin C (MMC) and MMC short read data (39, 94). No MMC viruses were classified as uninducible if they had a classifier confidence score ≥ 0.89. These predictions were compared to whether a no-MMC virus had a ≥ 2 fold increase (our threshold for induction) in PTH ratio in paired MMC samples (Supplementary Fig. S17). As a final test, we processed 340 bulk metagenomes and 37 highly enriched viromes using the UHVDB toolkit. The proportion of bulk viruses and enriched viruses classified as uninducible prophages was determined for each metagenome set (Supplementary Fig. S17).

### Updating UHVDB with cystic fibrosis related viruses

To identify shotgun metagenomes derived from CF respiratory tract samples, we searched NCBI BioProjects for non-gut CF shotgun metagenomes. This resulted in 144 BioProjects, with an additional 12 identified through a literature search. Then, we queried SRA to identify the 4,448 SRA accessions associated with these BioProjects. Accessions with obvious off target organism labels were removed leaving 2,444. BioProjects having ambiguous organism labels were manually investigated to remove off-target samples, leaving 2,222 samples. Several BioProjects contained non-CF samples or less than 10 total samples, and these were also removed, creating a final dataset of 1,949 accessions. The update workflow of the UHVDB toolkit was run on these accessions as well as 34 locally sequenced samples to download and preprocess reads, perform metagenome assembly, mine high-quality, confident viruses, annotate these sequences, and add them to UHVDB.

The 34 locally sequenced CF sputum samples had previously been collected monthly from 3 people with CF over the course of one year. These samples were selected from a biorepository of stored, frozen CF sputum collected through a home sputum collection study and maintained by the University of Michigan (the Michigan CF Sputum Archive) as previously described (99, 100). Collection and analysis of these samples complied with all ethical regulations and was approved by the Institutional Review Board of the University of Michigan Medical School (HUM00037056).

For processing these samples, 250 mg of frozen sputum was added to 250 µL of 10% DTT and vortexed for 1 minute at room temperature. Homogenized samples were then centrifuged for 3 minutes at 17,000 x g to pellet large debris. The supernatant was filtered using Millex-HV Syringe Filter Unit (Sigma; 0.45 µm, PVDF, 13 mm). 150 µL of filtrate was treated with 10 µL benzonase buffer, 38 µL PBS, and 2 µL Benzonase Nuclease (Sigma) to digest extracellular DNA. This mixture was incubated at 37°C for 15 minutes with agitation. To stop the reaction, 2 µL of 0.5 mM EDTA was added. To extract encapsulated DNA, we used the MagMax Viral/Pathogen Nucleic Acid Isolation Kit (Thermo). DNA concentration was determined using Qubit (Thermo) and DNA was sheared to an average fragment size of 350 bp using Covaris R230, and sequencing libraries prepared using xGen ssDNA & Low-Input DNA Library Prep Kit (IDT). Sequencing utilized300 cycle NextSeq™ 2000 P4 XLEAP-SBS™ Reagent Kits (Illumina).

To improve representation of viruses infecting canonical CF pathogens, we downloaded high-quality (completeness ≥ 90% and contamination < 5%) isolate assemblies and metagenome assembled genomes (MAGs) from these CF pathogen genera, including 37,734 Staphylococcus, 7,684 Pseudomonas, 4,559 Burkholderia, 1,311 Haemophilus, 985 Stenotrophomonas, and 319 Achromobacter genomes from mOTUs-db (18). Then, using the update functionality of the UHVDB toolkit, we identified high-quality, confident viruses in these assemblies and added them to UHVDB (Supplementary Fig. S18).

### Analysis of cystic fibrosis oral/respiratory metagenomes

After updating UHVDB to r5, the 1,983 CF metagenomes mentioned above were run through the analyze workflow of the UHVDB toolkit to determine virus/bacteria abundances and PTH ratios. Likely uninducible prophages were identified using our Random Forest classifier with confidence of ≥ 0.87 (95% precision threshold). After including these viruses for comparison in diversity and prevalence estimates, they were removed from the remainder of the analysis.

For longitudinal analyses of CF sputum samples, we identified individuals having a minimum of 2 samples containing ≥ 100 megabases of non-human reads and a total sampling period of at least 50 days. After calculating PTH ratios for each phage in each sample, we calculated the standard deviation of each phage species PTH ratio over time. An additional dataset of CF sputum samples collected before, during, and after tobramycin inhaled powder (TIP) therapy as part of a previously published study (101) were analyzed in the same manner to investigate phage-bacteria dynamics before, during and after antibiotic perturbations. Host relative abundances for this dataset were taken from the UHVDB toolkit’s sylph outputs.

### Creating UHVDB r6

After updating UHVDB’s CheckV database with new, complete viruses from NCBI RefSeq (102), we mined human-associated viruses from VIRE (55), metaVR (103), OAVGC (104), and MAGIC (105) using the update workflow of the UHVDB toolkit. Complete, human-infecting viruses were mined from NCBI Virus (106) as well, with only one SARS-COV2 genome per Pango lineage included. Species, genus, and family accumulation curves were extrapolated and used to estimate total diversity (S_Chao_) and likelihood of a novel virus representing novel taxa (SC: sample coverage) for each body site using iNext v3.0.2 (107).

### Profiling metatranscriptomes with the UHVDB toolkit

To evaluate UHVDB r6’s ability to profile human-infecting viruses from metatranscriptomes, we ran the analysis subworkflow of the UHVDB toolkit on a dataset of previously sequenced metatranscriptomes (108). Detected UHVDB species were assigned to a human virus label matching the authors categories using the nearest ICTV reference hit identified during the UHVDB update workflow. Human virus coverage data was downloaded from the supplementary information of Rajagopala et al. and compared to the coverage outputs by the UHVDB toolkit. Virus detections with a depth of coverage < 3 were removed to keep only confident hits.

## RESULTS

### Creating the Updateable Human Virome Database (UHVDB)

To build upon and generalize the foundation established by the Unified Human Gut Virus catalog (UHGV) (38), we developed an updated version of the UHGV virus classification pipeline (Figure 1A), which produced virus classifications consistent with those made by UHGV (Supplementary Fig. S1). To address the difficulties in classifying, annotating, and clustering fragmented virus sequences, we retained only high-quality viruses (≥ 90% complete) (Supplementary Fig. S2-3) (25, 26, 29, 30, 45, 109). Additionally, viruses classified as “uncertain” were further filtered using virus and plasmid hallmark hidden markov models (HMMs), retaining only sequences having ≥ 3 virus hallmarks and 0 plasmid hallmarks for downstream analyses (Supplementary Fig. S4). When applied to UHGV’s 208,604 high-quality viruses, our pipeline retained 201,324 (96.5%) sequences. These sequences were dereplicated to 179,392 unique viruses, comprising the first release of the Updateable Human Virome Database (UHVDB r1).

Next, we clustered UHVDB r1 into 156,387 genomovars (49) using thresholds of 99.5% average nucleotide identity (ANI) and 100% alignment fraction (AF), which corresponds with natural intra-species units due to an ANI gap (Supplementary Fig. S5). Genomovar representatives were then clustered into 53,595 species using 95% ANI and 85% AF (Supplementary Fig. S5) (45). Species representatives were clustered at the family through subgenus levels using proteomic similarity as described in UHGV (38, 45), resulting in 815 family and 18,547 genus-level clusters (Supplementary Fig. S6). To annotate UHVDB, genomovar representatives were run through an annotation workflow inspired by UHGV’s and that predicts taxonomy, host, function, and lifestyle for virus sequences (Supplementary Figs. S7-S11).

### Updating UHVDB

To expand UHVDB, additional sequences (viral databases and metagenome assemblies, described below) were run through the virus classification, filtering, clustering, and annotation workflow described above with the following notable changes to limit redundant computations and improve computational efficiency (Figure 1A): First, after classifying virus sequences, CheckV’s database was updated with new, complete viruses, which were identified by the presence of direct terminal repeats (DTRs) (42). This expanded database of complete viruses was used to re-estimate sequence completeness (Supplementary Fig. S2). Second, new viruses sharing a sequence hash (a unique, compressed digital fingerprint for biological sequences) with a virus already present in UHVDB were removed from downstream analyses. Third, only genomovar representatives not present in previous releases were run through the clustering and annotation workflows. Fourth, only proteins with novel sequence hashes were functionally annotated. Fifth, pairwise genome comparisons used for clustering were recycled from previous releases, so nucleotide and protein alignments from these pairs were not re-run.

For the first expansion of UHVDB (r2), we applied the updated workflow to the 3,812 high-quality, uncertain viruses in UHGV. Then, to consolidate existing human virome databases for the third release (r3), the update workflow was applied to 9 different human virome databases containing a total of 1,793,177 viral contigs (21, 28, 74–80). Last, to improve representation of non-gut viruses in UHVDB, we mined 79,131 ENA (88), Logan (89), and SPIRE (16) metagenome assemblies from human oral/respiratory tract, urogenital tract, and skin samples. These updates resulted in the fourth release (r4), which contained 575,497 unique viruses, nearly a 3-fold expansion over UHGV alone (Figure 1B).

Most (70.5%) of UHVDB r4’s unique sequences were from pre-existing databases, primarily UHGV (Figure 1C). However, the added metagenome assemblies did expand representation of viruses from non-gut body sites, with 155,129 oral/respiratory-derived viruses not found in previous virus databases (Figure 1C). Interestingly, comparatively few novel viruses were identified in skin (n=11,640) and urogenital tract (n=2,801) sample metagenomes despite mining 31,937 assemblies from these body sites (Figure 1C). In contrast to other human virome databases, UHVDB r4 contains exclusively high-quality viruses, with 24.3% of sequences predicted to be complete (Figure 1C). Therefore, UHVDB r4 represents an expanded compendium of high-quality viruses from multiple body sites compared with prior virus databases.

**Figure 1.**
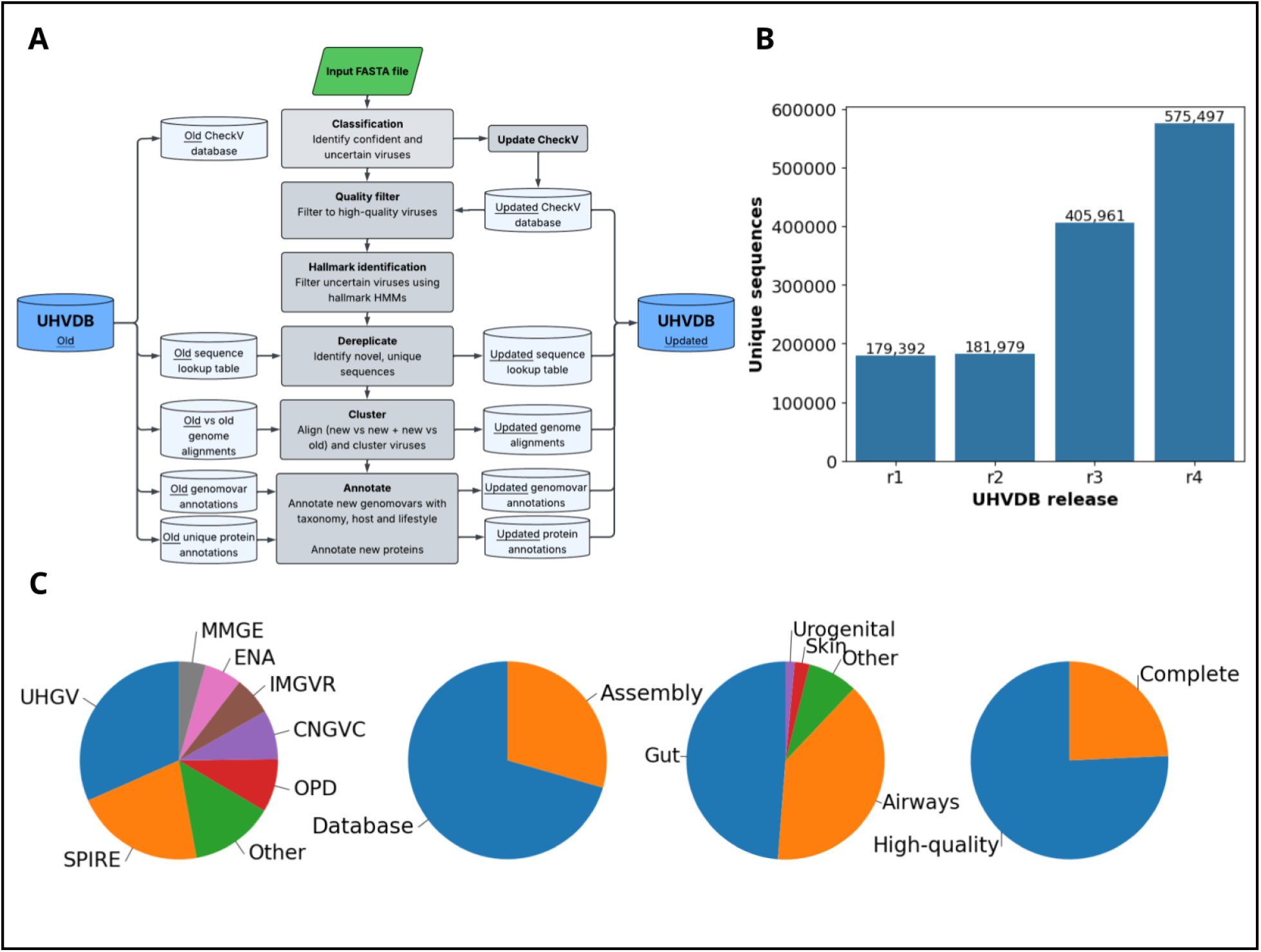
Overview of the creation and contents of the Updateable Human Virome Database (UHVDB). (a) Schematic of the virus classification, clustering, and annotation workflows used to create and update UHVDB. (b) Counts of unique viruses present in each release of UHVDB. In brief, the releases of UHVDB contain: r1) UHGV confident viruses r2) UHGV uncertain viruses r3) viruses from 9 different human virome databases r4) viruses from 79,131 human sample (non-gut) metagenome assemblies. (c) Proportion of UHVDB r4 unique sequences having different source annotations, from left to right: source dataset (UHGV: Unified Human Gut Virome Catalog (38), SPIRE: Searchable Planetary-scale Microbiome Resource (16), IMGVR: Integrated Microbial Genomes - Virus Resources v4 (21), OPD: Oral Phage Database (76), CNGVC: Chinese Gut Virus Catalog (77), MMGE: Metagenomic Mobile Genetic Elements (28), ENA: European Nucleotide Archive (81), “Other” includes the following: CNGVR: Chinese Gut Viral Reference (78), Logan (82), OVD: Oral Virome Database (75), SMGC: Skin Microbial Genome Collection (79), and VMGC: Vaginal Microbial Genome Collection(80)), source type (pre-existing virome database or metagenome assembly), body site of origin, and genome quality.

### Diversity of and comprehensivity of UHVDB

We next evaluated the diversity represented at different taxonomic levels in UHVDB r4. At the species level (95% ANI and 85% AF) (45), there were 199,442 clusters, nearly four times the number in UHGV (38). Despite this expansion, species clusters remained largely consistent across releases, with only 1.1% of clusters split and 3.7% merged between r1 and r4 (Supplementary Fig. S6). Nevertheless, accumulation curves indicated that r4 was not saturated at the species level for the gut, oral/respiratory tract, skin, or urogenital tract, meaning that new viral species were continuously being identified with each added dataset (Figure 2A).

To estimate diversity at higher ranks, UHVDB r4’s species representatives were clustered using 5.5% (family) and 65% (genus) protein similarity, yielding 41,354 genus and 1,444 family-level clusters (Figure 2B-C; Supplementary Fig. S6, S12), over twice as many genera and three times as many families as in UHGV (19,782 genera and 454 families). Among these clusters, 6,496 genera (15.7%) and 154 families (10.6%) exclusively contained viruses found in metagenome assemblies. Accumulation curves began to plateau at higher taxonomic ranks, particularly the family level (Figure 2C).

To evaluate how comprehensively UHVDB r4 represents viruses from the human virome, we assembled 400 public metagenomes comprising 100 fecal, 100 oral/respiratory tract, 100 skin, and 100 urogenital samples not included in UHVDB. Using our workflow, we identified high-quality, confident viruses from these assemblies. These were aligned to 4 different virus databases (21, 38, 83). The highest global ANI (ANI x AF) for each assembled virus was then determined with respect to each database. For fecal-derived viruses, UHGV, IMG/VR, and UHVDB achieved similar mean global ANIs of 79.5%, 80.0%, and 83.6%, respectively (Figure 2D). However, UHVDB displayed higher Pearson correlations and recall for all other sample types, with mean global ANI values of 85.0% (oral/respiratory tract), 78.7% (skin), and 88.1% (urogenital) (Figure 2D).

### Body site specificity of UHVDB clusters

Next, we determined whether specific virus clusters were identifiable across different body sites. At the genomovar level, 98.0% of non-singleton clusters contained virus sequences exclusively from one body site. At the species level, this value remained high at 94.0% (Figure 2E), indicating that most virus genomovar and species clusters are body-site specific. However, this percentage decreased at the genus (87.8%) and family (48.9%) levels.

**Figure 2.**
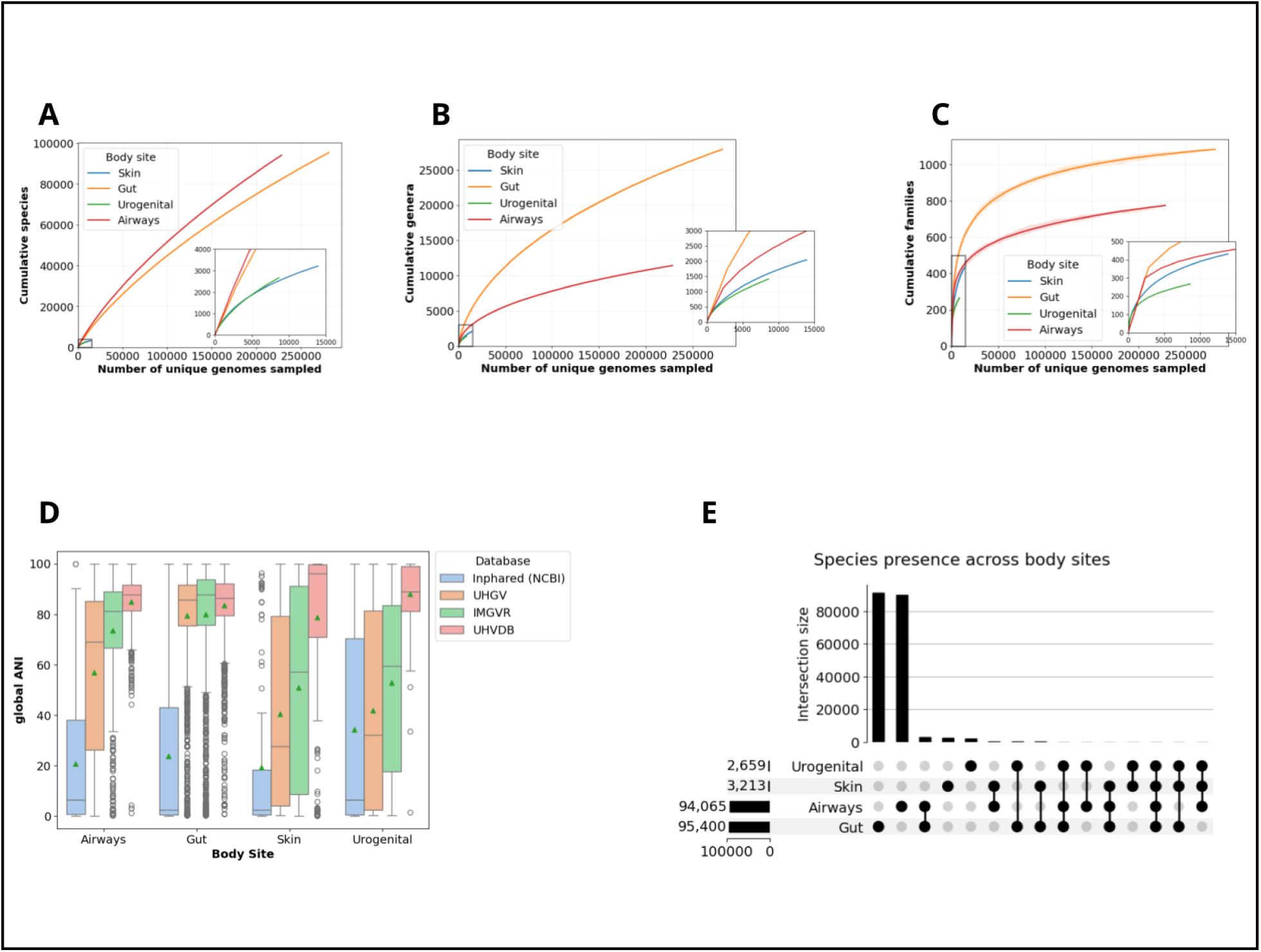
Diversity, comprehensivity, and body site specificity of UHVDB viruses. Accumulation curves at the (a) species (95% ANI and 85% AF) (b) genus (65% proteomic similarity) and (c) family (5.5% proteomic similarity) levels. UHVDB release 4 (r4) unique sequences were organized by body site and split into 100 increasing subsets, with the number of clusters present at each interval determined. This process was repeated 50 times. Lines depict the mean counts at each interval and shaded regions represent the area between the minimum and maximum values. (d) Maximum global ANI values obtained when aligning novel viruses assembled from 400 metagenomes from different body sites to 4 different human virome databases. Mean global ANIs are indicated by a green triangle. (e) Body site composition of UHVDB r4 species clusters. This plot displays the total number of species clusters containing viruses from each body site (bottom left bar chart). Additionally, the figure depicts the body site composition of different clusters (bottom right), with the number of species clusters composed of these combinations (top bar chart).

### Characteristics of UHVDB

We next analyzed the characteristics (taxonomy, host, lifestyle, function) of all genomovar representatives in UHVDB r4. First, taxonomy was assigned at the class level using geNomad (25). Across all body sites, Caudoviricetes (tailed dsDNA phages) was the most prevalent taxonomic class, accounting for 92.7% of UHVDB genomovars. The next most prevalent classes were Malgrandaviricetes (4.2%) and Faserviricetes (1.9%), both of which are ssDNA phages. In addition to phages, we detected eukaryotic viruses belonging to Papovaviricetes (0.5%) and Cardeaviricetes (0.4%). Interestingly, certain classes appeared to be associated with specific body sites (Figure 3A). For example, Faserviricetes was particularly prevalent in oral/respiratory tract and skin samples, while Malgrandaviricetes was more prevalent in gut and urogenital samples. Additionally, Papovaviricetes accounted for 26.9% of genomovars identified in skin metagenomes, while accounting for fewer than 5% of genomovars in all other body sites.

To assign taxonomy below the class level, we aligned genomovar representatives to viruses annotated by the International Committee for the Taxonomy of Viruses (ICTV) and calculated proteomic similarity values. UHVDB viruses with family (≥ 5%) or genus (≥ 65%) -level hits were assigned the reference virus’ taxonomy at the respective level. Using this approach, 58.9% and 5.6% of UHVDB genomovars had hits at the family and genus levels, respectively. However, many reference Caudoviricetes genomes lack family-level annotations, and therefore a majority (67.4%) of UHVDB viruses aligned to reference genomes were not assigned to an ICTV family.

Bacterial/archaeal hosts were assigned to genomovars based on a combination of CRISPR spacer matching and identification containment in bacterial genome assemblies (18, 54, 55). Host bacterial species, genera, and families could be predicted for 46.3%, 81.7%, and 88.9% of UHVDB genomovars, respectively. Predicted host families differed by body site; Streptococcaceae was the most prevalent predicted host among oral/respiratory tract viruses and Bacteroidaceae and Lachnospiraceae were the most prevalent among gut viruses. For the skin, Staphylococcaceae phages and eukaryotic viruses were the most prevalent, while Lactobacillaceae and Bifidobacteriaceae phages were the most prevalent in urogenital samples (Figure 3B). For genomovars taxonomically assigned to classes of eukaryotic viruses (Arfiviricetes, Cardeaviricetes, Papovaviricetes, Quintoviricetes, and Repensiviricetes), “Eukaryote” was assigned as the host.

Next, we predicted virus lifestyles by classifying viruses with ≥ 1 of the following signals as temperate: classification as an integrated provirus, BACPHLIP score > 0.5 (temperate), the presence of an integration-related gene (25, 42, 73). All other genomovars were classified as virulent. Most UHVDB genomovars contained one of these signals (87.4%) and were therefore classified as temperate (Figure 3C; Supplementary Fig. S10). The comparatively lower percentage of skin viruses classified as temperate was largely due to the higher proportion in skin metagenomes of eukaryotic viruses (29.6%; Figure 3A), which cannot be classified as temperate using our phage-centric approach.

After predicting and dereplicating proteins from UHVDB r4 genomovar representatives, we functionally characterized these sequences using several different approaches. First, we assigned 82.2% of the 15,114,978 unique proteins predicted from genomovar representatives to a UniProt or InterPro accession using Bakta, foldseek, and InterProScan (Supplementary Fig. S11) (53, 58, 62, 66). Then, each unique protein was also analyzed using Pharokka, Phold, and Empathi to identify putative phage-related functions. In total, 77.6% of unique proteins were annotated as phage-related by at least one of these tools (Supplementary Fig. S11). To evaluate the sensitivity of our phage annotations, we searched for three highly-conserved Caudoviricetes genes (major capsid protein, portal protein, terminase large subunit) and found that ≥ 2 of these genes were detected in 94.4% of Caudoviricetes genomovars (Figure 3D). By comparison, a similar search for antibiotic resistance genes (ARGs) and virulence factors (VFs) detected only 167 ARGs and 2,684 VFs among UHVDB r4 proteins (Supplementary Fig. S11).

**Figure 3.**
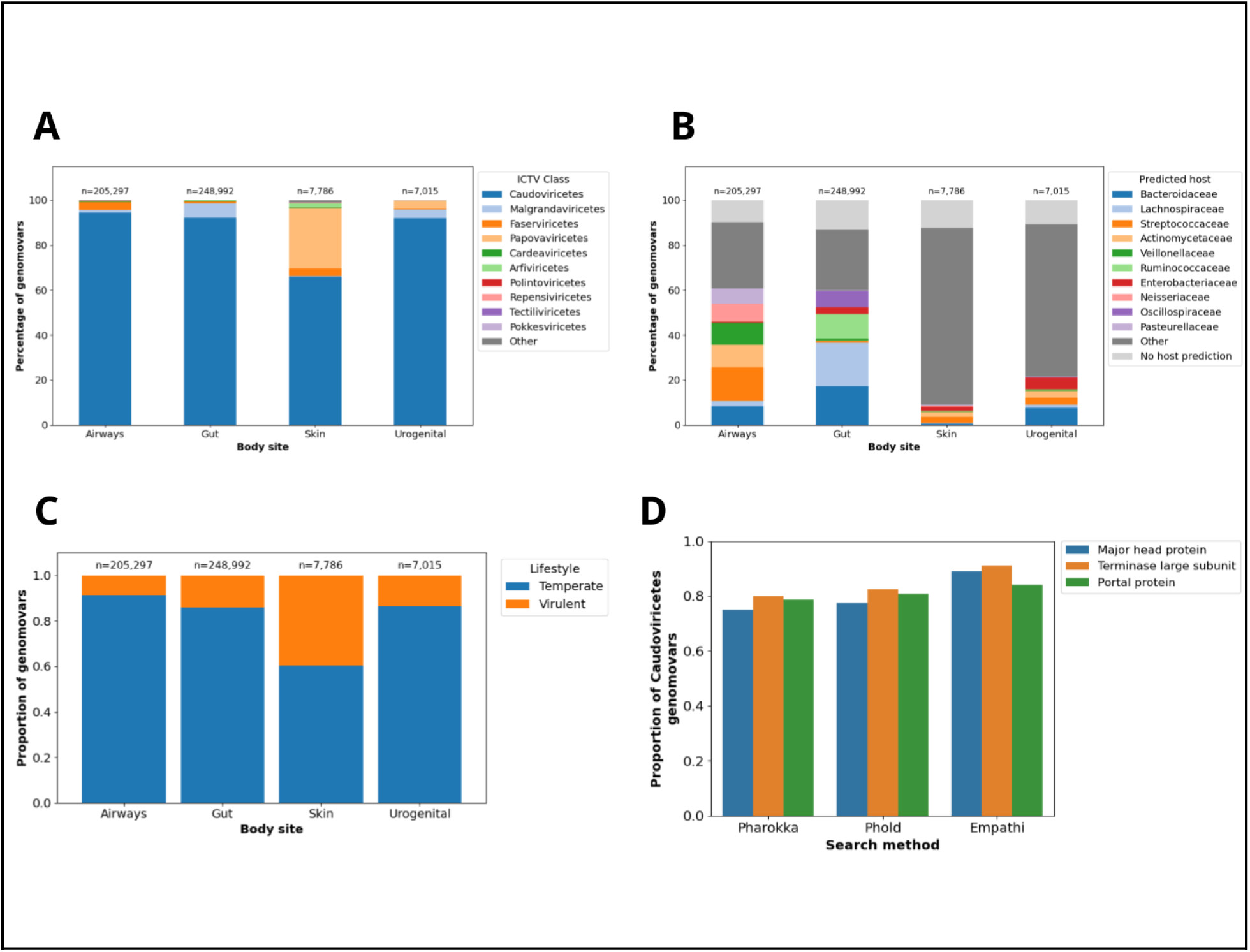
Characteristics of the Updateable Human Virome Database. Shown are proportions of UHVDB release 4 (r4) genomovar representatives assigned to an (a) ICTV class, (b) predicted host families, and (c) lifestyle split by body site of origin. The 10 most prevalent ICTV classes and host families are displayed, with all other predictions classified as “Other”. The proportion of viruses without host predictions is indicated by “No host prediction”. (d) Cumulative proportion of Caudoviricetes genomovars for which single-copy portal, major head protein, and terminase large subunit proteins were detected after using increasingly sensitive search methods.

### A toolkit for standardized, updateable human virome analyses

To aid in future UHVDB updates, we implemented the workflows we had used to create and expand UHVDB as a Nextflow pipeline called the UHVDB toolkit (Figure 4A). This pipeline enables users to recapitulate our methods and mine viruses from new human metagenomes, addressing a major problem in the field of viral metagenomics: the inability to easily update existing virus databases. In addition to the update functionalities described above (Figure 1A), we also designed UHVDB’s annotations with updates in mind. For taxonomic assignments, the ICTV accession most similar to each genomovar representative is stored, so that changes in reference taxonomy can easily be propagated to related UHVDB sequences. Similarly, bacterial and archaeal accessions used for host predictions are stored. For protein annotations, the association of UHVDB proteins with UniProt and InterPro accessions will enable updates to these resources to be utilized by future users of UHVDB.

To make UHVDB as usable as it is updateable, we added an analysis workflow that taxonomically profiles virus and bacterial species using sylph (87) (Figure 4A). To benchmark this approach, we split the viruses assembled for our comprehensivity analysis (Figure 2D) into multiple batches, each containing 100 viruses from a specific body site, and simulated reads *in silico* (90). Then, we ran the UHVDB toolkit, BAQLaVa (110), Marker-MAGu (93), and Phanta (92) on these simulated reads to compare performance with existing taxonomic analytical tools. Input viruses were aligned to each tool’s database of virus genomes and the most similar viruses were identified using global ANI. Input values were then compared to the outputs from each tool’s most similar virus so that accuracy (Pearson R) and recovery (recall) could be calculated. Across all body sites, the UHVDB toolkit yielded improved accuracy (Pearson R: UHVDB = 0.96, BAQLaVa = 0.58, Marker-MAGu = 0.57, Phanta = 0.0.49) and improved sensitivity (Recall: UHVDB = 82.4%, BAQLaVa = 53.5%, Marker-MAGu = 35.1%, Phanta = 28.7%) relative to all other tools (Figure 4B; Supplementary Fig. S13).

### Evaluating phage-to-host ratios

To evaluate how well UHVDB predicts phage-bacteria pairs, we analyzed properties of these predictions across a dataset of fecal metagenomes. We found that predicted pairs exhibited high co-occurrence and strong abundance correlations (Supplementary Fig. S14). We then calculated phage-to-host (PTH) ratios by dividing phage depth of coverage by predicted host species depth of coverage; Our PTH ratios were consistent with those calculated using PropagAte (95) (Pearson R = 0.825) but with relatively higher sensitivity (Supplementary Fig. S15). To evaluate the biological utility of PTH ratios, we ran the UHVDB toolkit and PropagAte on shotgun metagenomes from 10 bacterial isolates that had been treated with and without mitomycin C (MMC) to induce prophages (96). Both tools were able to detect induced proviruses for 3 different bacterial species, each of which exhibited a > 2-fold increase in PTH ratios with MMC compared with no MMC treatment (Figure 4C). Additionally, viruses with the highest PTH ratios in unenriched (bulk) metagenomes were the most likely to be detected in paired enriched (cellular depletion) metagenomes (94) (Supplementary Fig. S15). These findings suggest that the UHVDB toolkit’s PTH ratios could serve as a proxy for prophage induction or replication activity in metagenomes.

### Identifying likely active phages in metagenomes

Not all viruses detected in a metagenome represent active or potentially active viruses; some represent cryptic (uninducible) proviruses, which are prevalent among bacterial and eukaryotic genomes (111, 112). These different viral states can affect their environments in different ways; for example, active viruses (lytic viruses or inducible proviruses) can lyse their hosts and create infectious particles, while inactive (uninducible) viruses cannot.

Consequently, distinguishing between active and uninducible viruses has important biological implications. We therefore incorporated tools to predict virus activity into our toolkit.

One strong signal of provirus domestication, or inactivation, is genome truncation (41) (Supplementary Fig. S3). The detection of such inactive viruses is limited in part by the fact that UHVDB contains only high-quality (non-fragmented) genomes. To identify other signals of provirus (specifically prophage) inactivation, we used a dataset of 39 paired bulk and virus-enriched metagenomes from infant fecal samples (94). Viruses detected both in bulk and paired enriched metagenomes were considered active or inducible, while those detected only in bulk metagenomes were considered uninducible prophages. We then evaluated the ability of read alignment metrics (i.e. breadth/variance of coverage, gene breadth) and UHVDB annotations (i.e. presence of a DTR, virus score) to identify the prophages considered uninducible. Although false negatives (inducible prophages not detected in enriched metagenomes) likely existed in our training dataset, we were encouraged to find that many distinguishing features correspond with expected signals of uninducibility. For example, low breadth of coverage, which may be due to genome truncation, was associated with lack of detection in enriched metagenomes (Supplementary Fig. S16). Furthermore, the absence of essential genes (i.e. capsid), which is expected to affect a virus’ ability to form infectious particles, was also associated with detection exclusively in bulk metagenomes (Supplementary Fig. S16).

We next sought to integrate different signals for identifying uninducible prophages using a random forest classifier. Given that the tools we used to annotate UHVDB proteins were primarily developed for Caudoviricetes genomes, we limited our training and testing datasets to sequences assigned to that class. Classifier performance was evaluated using group 5-fold cross validation, grouping sequences by their sample of origin, resulting in a pooled area under the precision recall curve (AUPRC) of 0.920 (Figure 4D). When evaluating this classifier on another dataset of paired bulk and enriched fecal metagenomes (98), we found that the 95% precision (41.0% recall) threshold established using our training set achieved similar performance (94.5% precision, 47.5% recall; Supplementary Fig. S17). Furthermore, the classifier achieved nearly 90% precision in identifying prophages not induced by mitomycin C in bacterial isolates, and this tool rarely classified viruses from enriched metagenomes as uninducible (Supplementary Fig. S17).

**Figure 4.**
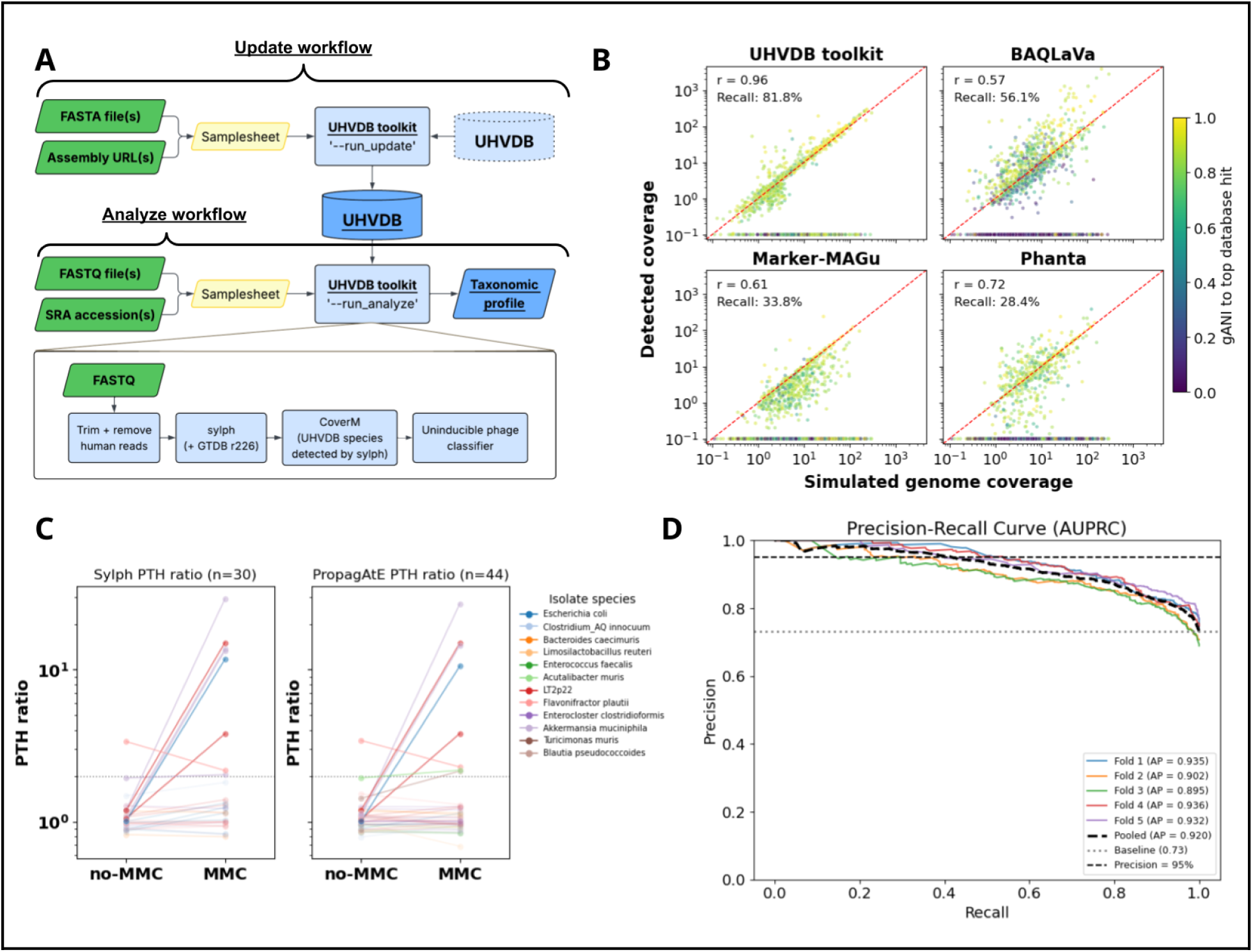
The UHVDB toolkit. (a) Schematic overview of the UHVDB toolkit. (*Top*) The “update” workflow, which mines, clusters, and annotates viruses to update UHVDB. (*Bottom*) The “analyze” workflow, which identifies contained UHVDB species with sylph, aligns reads to contained species with CoverM, and classifies phage species likely to be uninducible (inactive). (b) Performance of different virome profilers (UHVDB toolkit, BAQlava, Marker-MAGu, and Phanta) evaluated using short reads simulated from 400 metagenome assemblies not included in any tool’s virus database. Coverage values for each input virus were compared to output coverage values from the most similar virus contained in each tool’s database using global ANI (ANI x AF). Pearson R values and recall (percent of input viruses detected) are presented in the top left of each plot. Each point represents one input virus, which is colored based on the global ANI between the input virus and the most similar virus in each tool’s reference database. The red dashed line represents y=x. (c) Calculated phage-to-host (PTH) ratios from prophages detected in 10 bacterial isolates sequenced both with and without mitomycin C (MMC) treatment. PTH ratios were calculated using UHVDB toolkit (sylph, left) and PropagAte (right) and ratios derived from the same virus species are connected by a line across conditions. The number of virus species detected by each tool is depicted at the top, and the dashed horizontal line depicts at PTH value of 2. (d) Precision-recall curve for our uninducible prophage classifier. Each fold represents a subset of samples from our training set with the “Pooled” value indicating the mean performance across all folds. The baseline value of 0.73 is depicted by a light grey dotted line and our selected threshold of 95% precision is depicted by a black dashed line.

### Updating UHVDB with cystic fibrosis related viruses

To explore the utility of the UHVDB toolkit in a clinical context, we applied it to a collection of respiratory specimens (sputum, oropharyngeal or nasopharyngeal swabs, or bronchoalveolar lavage fluid) from people with Cystic Fibrosis (CF). We chose this application because CF lung disease is characterized by chronic, recalcitrant bacterial respiratory tract infections, and while there is growing interest in phage therapy for these infections, little is known about how phages and bacteria interact in the CF respiratory tract. Additionally, evidence suggests that phages can contribute to clinically relevant changes in the virulence and antibiotic resistance of bacterial pathogens in CF (113). Therefore, accurately characterizing the active phage in CF respiratory samples can provide clinically useful insights.

Using the UHVDB toolkit, we updated UHVDB’s database to release 5 (r5) by assembling and mining high-quality, confident viruses from 1,949 publicly-available shotgun sequencing datasets and 34 newly sequenced metagenomes from CF respiratory samples (101, 114–131). Furthermore, we mined viruses from isolate genomes and metagenome assembled genomes (MAGs) of conventional CF pathogens (*Staphylococcus, Pseudomonas, Burkholderia, Haemophilus, Stenotrophomonas, Achromobacter*) to improve representation of viruses infecting these pathogens. Together, these mining efforts added 42,318 unique viruses, 27,417 genomovars and 6,847 species to UHVDB (Supplementary Fig. S18).

### Characteristics of the cystic fibrosis respiratory tract virome

Using UHVDB r5, we applied the UHVDB toolkit to all 1,983 CF respiratory sample metagenomes and calculated virus species richness. Without removing likely uninducible Caudoviricetes prophages, the most deeply sequenced samples (≥ 1 Gbp) contained an average of 112.5 virus species for oropharyngeal and nasopharyngeal swabs (swabs) and 58.9 species per sputum/bronchoalveolar lavage (sputum/BAL) sample (Figure 5A; Supplementary Fig. S19). However, after removing viruses predicted to be uninducible by our classifier (precision threshold ≥ 95%), per-sample richness dropped to 79.5 for swabs and 39.6 for sputum/BAL samples (Figure 5A), suggesting that approximately 30% of detected viruses may represent active or inducible viruses. Importantly, the classification of a virus species as uninducible is sample-specific, so a species may be labeled as uninducible in one sample but not another. Of the 17,398 UHVDB species detected in this analysis, 71.2% were never classified as uninducible (“never uninducible”), 13.8% were classified as uninducible for some samples but not others (“sometimes uninducible”), and 15.0% were always classified as uninducible (“always uninducible”) (Supplementary Fig. S19). In an effort to focus our analysis on putatively inducible or active viruses, we removed viruses classified as uninducible prophages from the remainder of the analysis.

Next, we analyzed the prevalence of UHVDB taxa across unique specimens in this cohort and found that many virus species were only detected in a small percentage (swab mean=0.51%; sputum/BAL mean=0.40%) of the most deeply-sequenced (≥ 1 Gbp) samples. However, prevalence values increased at higher taxonomic ranks, with mean prevalence values at the family level being 6.64% for swabs and 4.44% for sputum/BAL (Figure 5B).

Despite the overall low prevalence of UHVDB taxa, a subset of species, genera, and families were detected in ≥ 10% of swab and sputum/BAL samples (Supplementary Fig. S19). Interestingly, one such species belonged to the ICTV class Arfiviricetes, which contains the family Redondoviridae, a virus family associated with periodontal disease and critical illness (132, 133).

Next, we analyzed the taxonomy, hosts, and lifestyles of viruses detected in CF swabs and sputum/BAL samples. Taxonomically, most detected viruses were phages, with a higher prevalence of filamentous phages (Faserviricetes) in swabs compared to sputum/BAL samples (Figure 5C). Predicted host families were similar across sample types, although phages infecting Streptococcaceae, Pseudomonadaceae, and Staphylococcaceae were more prevalent in sputum/BAL samples than swabs (Figure 5D). In both swabs and sputum/BAL samples, most viruses were classified as temperate (Figure 5E).

**Figure 5.**
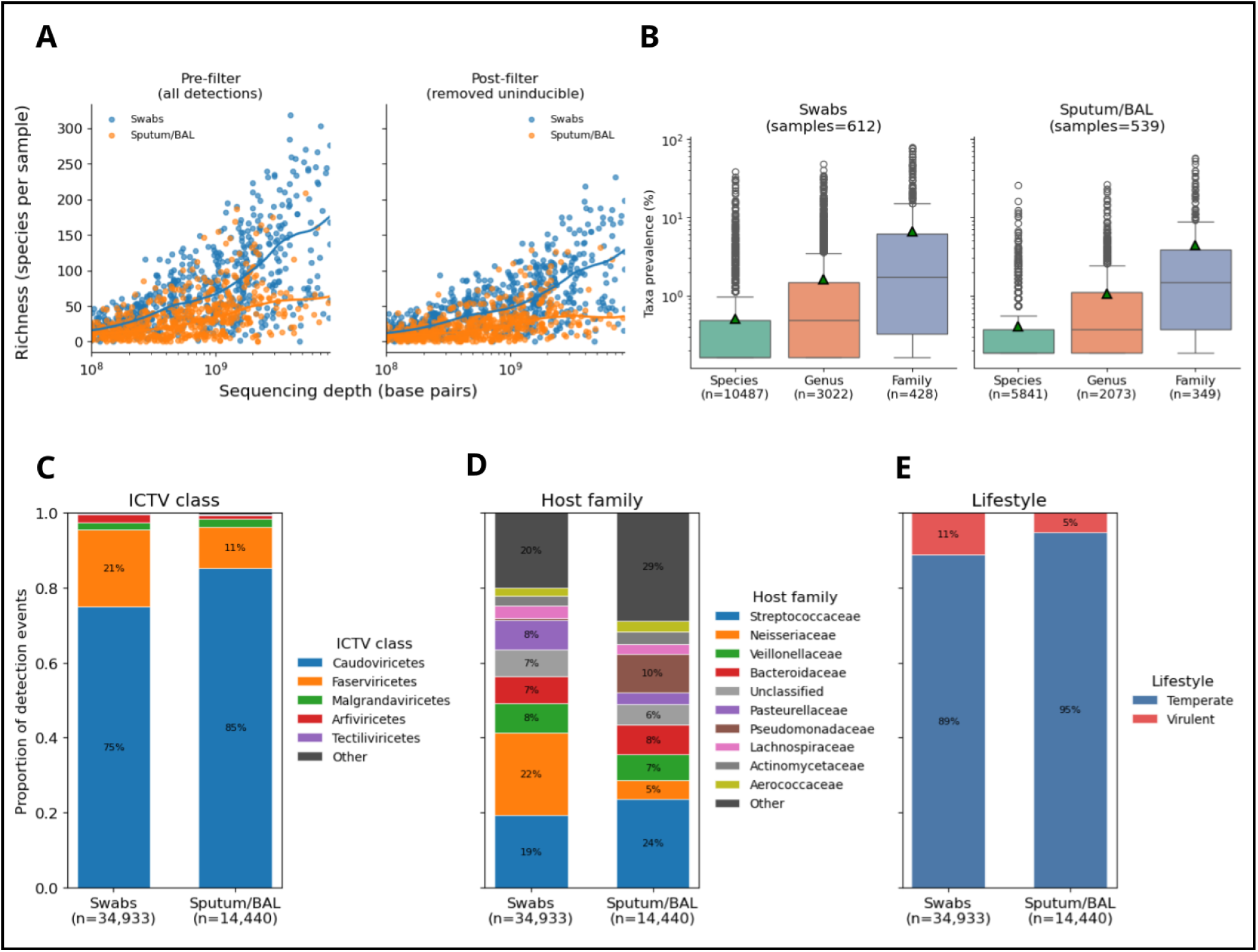
Characteristics of the CF respiratory sample virome. (a) Number of UHVDB species detected in CF oropharyngeal and nasopharyngeal swabs (Swabs) and sputum/bronchoalveolar lavage samples (Sputum/BAL) at increasing sequencing depths. The number of species is shown before and after removing sequences labeled as uninducible prophages by our random forest classifier. Each point represents one metagenome, and the lines were drawn using locally weighted scatterplot smoothing (LOWESS). (b) Prevalence, or the percentage of high sequencing depth samples (≥ 1 Gbp) that a UHVDB species, genus, or family was detected in. Prevalence was calculated separately for taxa exclusively classified as uninducible and taxa either never classified as uninducible, or only sometimes classified as uninducible. The proportion of UHVDB species detections not classified as uninducible having different (c) ICTV class, (d) bacterial host family, and (e) lifestyle annotations.

### Virus dynamics in CF respiratory tract samples

To investigate phage-bacteria dynamics in CF respiratory sample metagenomes, we calculated PTH ratios for all phage species not classified as uninducible with a predicted host species detected in the same sample (n=17,450; 34.3% of detections). PTH ratios were generally close to one (median=1.04), although some phages infecting several host genera consistently displayed PTH values > 2 (Supplementary Fig. S19).

Next, we focused our analysis on bacteria particularly relevant to CF, the conventional CF pathogens (*Achromobacter spp., Burkholderia spp., Haemophilus influenzae, Pseudomonas aeruginosa, Staphylococcus aureus,* and *Stenotrophomonas spp.).* Most phages infecting CF pathogens had PTH ratios of approximately one, with an overall median of 1.08 (Figure 6A). However, a subset of phages exhibited PTH ratios ≥ 2 (12.4%), and these were more commonly observed in samples where the CF pathogen comprised <50% relative abundance in the bacterial microbiota compared to samples where the CF pathogen comprised ≥50% relative abundance (Supplementary Fig. S20). Certain phage species consistently yielded PTH ratios ≥ 2, mostly common among *Pseudomonas*, *Stenotrophomonas,* and *Burkholderia* phages (Supplementary Fig. S20). Interestingly, 42.5% of phage species exhibiting a PTH ratio ≥ 2 in at least one sample belonged to the ICTV class Faserviricetes (filamentous ssDNA phages), which contains the clinically-relevant but non-lytic *Pseudomonas* infecting Pf phage (134).

We explored the stability of PTH ratios over time within patients for phages that infect CF pathogens. First, we identified individuals (n=32) having CF pathogen phages detected in at least 2 timepoints ≥ 50 days apart. Then, we calculated the standard deviation of PTH ratios across timepoints and found them generally to be low (median=0.18) (Figure 6B).

We additionally sought to evaluate phage-bacteria dynamics during antibiotic treatment, which could conceivably decrease phage abundances by killing host cells or, alternatively, increase phage production by inducing bacterial stress responses. We therefore analyzed a dataset of 157 sputum samples collected weekly from 30 individuals before, during, and after a month-long treatment cycle of a commonly-used CF maintenance antibiotic, tobramycin inhaled powder (TIP) (101). Again, PTH ratios for phages infecting CF pathogens were often close to one (median=1.04), largely consistent on and off TIP, and stable over time (Supplementary Fig. S20). Additionally, we did not observe a clear relationship between PTH ratios and changes in CF pathogen relative abundance over the course of treatment (Figure 6C).

**Figure 6.**
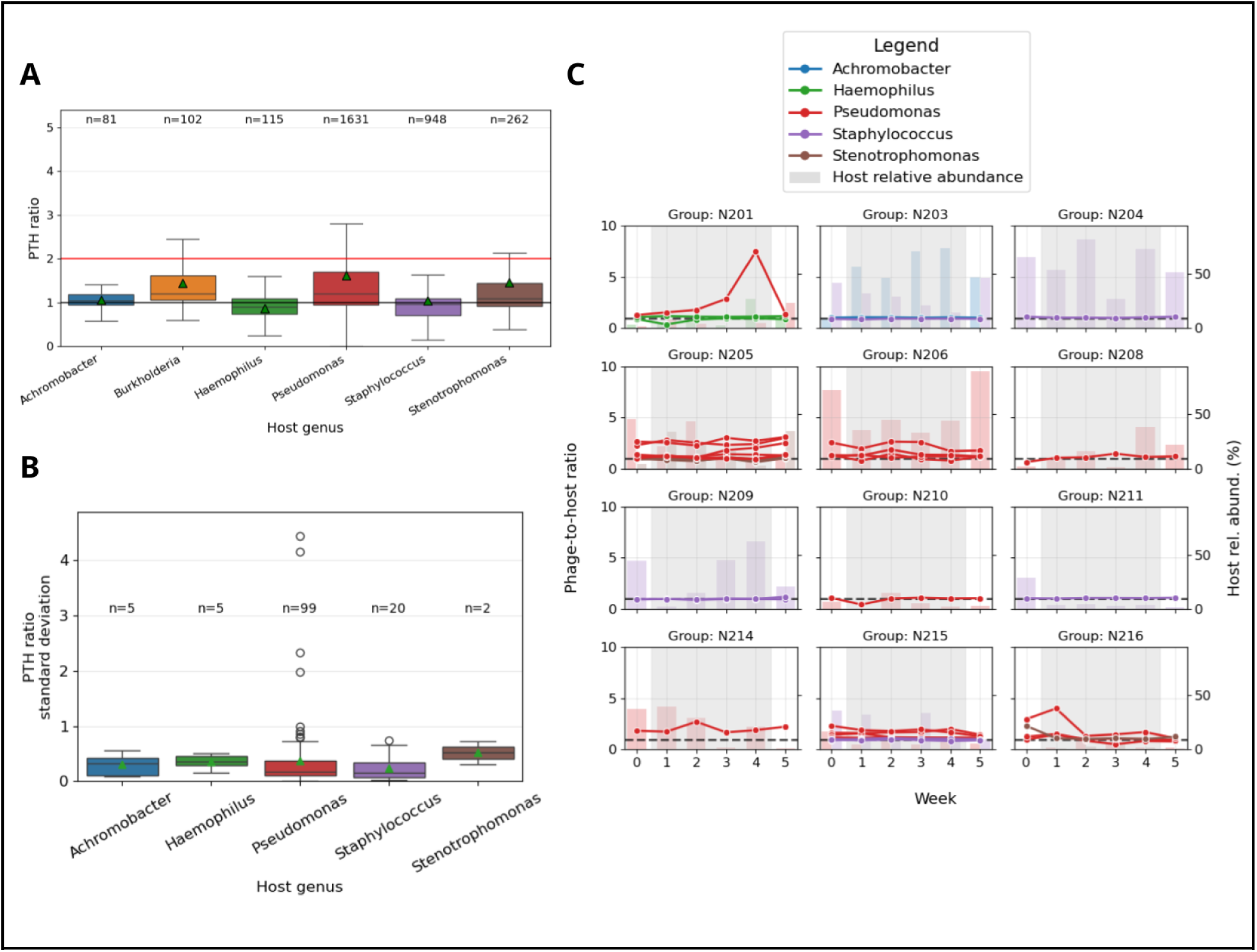
Phage-bacteria dynamics in CF respiratory samples. Phage-to-host (PTH) ratios were calculated for phages co-detected with their predicted host species. (a) PTH ratios for phages infecting conventional CF pathogens. Counts indicate the number of phage detections, the black horizontal line depicts a PTH of 1, the red line depicts a PTH of 2, and green triangles indicate the mean. (b) Standard deviation of PTH ratios for phages detected in ≥ 2 samples from the same person over at least 50 days. Numbers above each box indicate the number of phage species tracked longitudinally and green triangles indicate mean values. (c) Longitudinal analysis of PTH ratios before, during, and 1 month after tobramycin inhaled powder (TIP) treatment. Each panel represents one person with CF and each line represents one phage species, colored by host genus. The background barcharts and related right y-axis depict CF pathogen host species relative abundance after human read removal. The grey background indicates the TIP treatment period.

### Continued expansion and improvement of UHVDB

During preparation of this manuscript, several major virome databases were released (55, 103), and additional, pre-existing databases were identified (104, 105). To illustrate the updateablity of our database, we used the UHVDB toolkit to mine 1,261,909 virus sequences in these databases. The resulting sixth database release (UHVDB r6) nearly doubled UHVDB, bringing the number of unique viruses to 1,155,490 (Figure 7A), with many of the new sequences coming from the VIRE and OAVGC databases (Figure 7D). This update also substantially increased the number of species (from 206,289 to 291,605) and genera (from 43,125 to 53,506) represented in UHVDB (Figure 7B-C), while the number of families increased only modestly (from 1,446 to 1,590). We extrapolated our accumulation curves, finding that, while there is likely substantial species and genus-level diversity to be discovered, the rate of additional virus discovery is expected to be slow, with most viruses likely already represented in the species and genus clusters already present in UHVDB (Supplementary Fig. S21).

To allow UHVDB to more comprehensively reflect the human virome beyond human-associated phages, we also added 69,971 complete human-infecting viruses (DNA and RNA) to UHVDB r6 from NCBI VIRUS (Figure 7D) (106). With these additions, bacteria, phages, and human DNA/RNA viruses can be profiled in parallel using metatranscriptomes. We tested this functionality using previously sequenced metatranscriptomes from nasopharyngeal swabs of children both with and without respiratory illness (108), finding that the results of the UHVDB toolkit human RNA/DNA virus profiling agreed strongly with the original study findings (Figure 7E; Supplementary Fig. S21). Upon further analysis of this dataset, we observed substantial intra-species diversity among human-infecting RNA viruses; for example, 6 different UHVDB species clusters represented the 9 Human Rhinovirus A (HRV-A) detections in this dataset (Supplementary Fig. S21). This finding highlights the utility of UHVDB’s genome sequence-based outputs, as opposed to outputs that define viral composition by taxonomic labels alone, in reflecting the sequence diversity within viral taxa with comparatively high taxonomic resolution.

**Figure 7.**
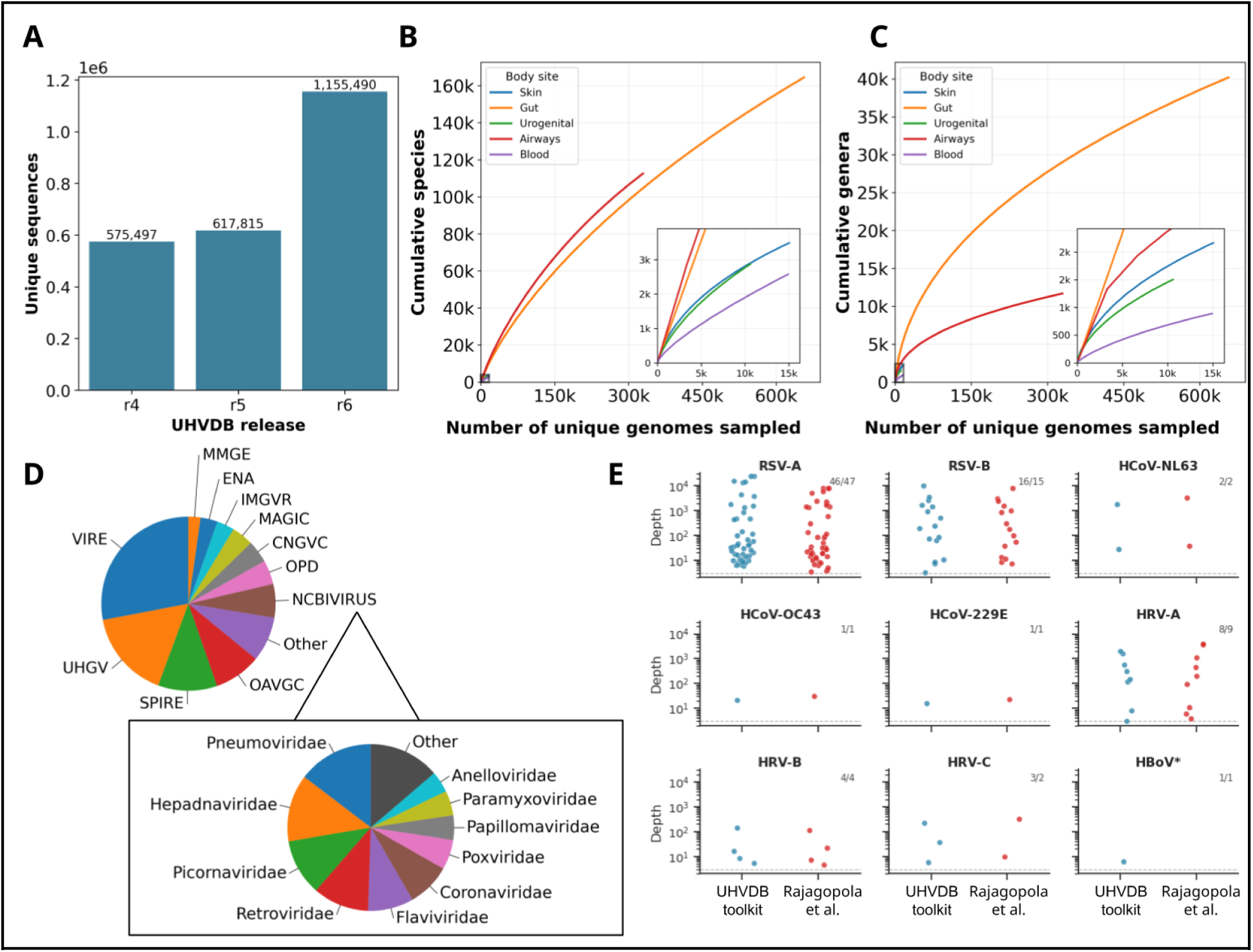
Expansion of UHVDB for release 6 (r6). (a) Number of unique sequences in UHVDB r4 (Figures 1-4), r5 (includes CF-related viruses), and r6 (includes recent virome databases). UHVDB r6 accumulation curves at the (b) species and (c) genus levels. (d) Pie chart depicting source databases (top pie chart) for unique viruses in UHVDB r6 (Added databases: VIRE (55), OAVGC (Oral and Airway Viral Genome Catalogue, (55, 104)), NCBIVIRUS (NCBI_VIRUS (106)). The lower pie chart depicts the ICTV family composition of human viruses added from NCBI. (e) Depth of coverage for human-infecting viruses detected in clinical metatranscriptomes using the UHVDB toolkit (UHVDB) and data from Rajagopola et al. (108). Counts in the top right of each panel indicate the number of samples (58 total samples) determined to contain the specified virus using the UHVDB toolkit or Rajagopola et al. data. Virus abbreviations: RSV: Respiratory syncytial virus, HCoV: Human coronavirus, HRV: Human rhinovirus, HBoV: Human bocavirus. * HBoV was reported as detected by Rajagopala et al. but no coverage data was included in the original supplement.

## DISCUSSION

Current approaches for studying the human virome are limited by a lack of standardization and the largely fragmented nature of viral sequences populating the many existing databases (21, 28, 38, 135). Moreover, many different tools are used to cluster and annotate virus sequences, further complicating cross-study comparisons. The Updateable Human Virome Database (UHVDB) is an attempt to address these issues by using a standardized virus mining workflow to identify and annotate high-quality viruses. Initially, we mined 1.8 million virus sequences and 80,000 metagenome assemblies to create the first (r1) through fourth (r4) releases of UHVDB (Figure 1). This effort uncovered substantial virus diversity in samples from different human body sites (Figure 2A-C) and increased the representation of non-gut viruses relative to previous human virome databases (Figure 2D). We observed that virus species tended to be body-site specific (Figure 2E) and that virus taxonomy and predicted bacterial hosts differed by body site (Figure 3A-B), although temperate phages consistently predominated regardless of body surface sampled (Figure 3C). The predominance of temperate phages contrasts with previous virome databases, in which only 13-55% of viruses were classified as temperate (32, 105, 136), likely owing to the inclusion of fragmented genomes and the use of less sensitive methods for detecting integration-related genes than used here.

We developed a Nextflow pipeline, the UHVDB toolkit, that enables users to update UHVDB with novel virus sequences, taxonomically profile viruses, calculate phage-to-host (PTH) ratios, and identify likely uninducible prophages in a metagenome (Figure 4A-D). By applying this toolkit to mine viruses from nearly 2,000 cystic fibrosis (CF) respiratory tract sample metagenomes and over 50,000 CF bacterial isolate assemblies, we created the fifth release of UHVDB (r5). The UHVDB toolkit was then used to profile these CF metagenomes, and we found that roughly one fourth of detected virus species were likely uninducible prophages (Figure 5A). After removing these viruses, the most deeply sequenced sputum and bronchoalveolar lavage (BAL) samples contained an average of 43.8 virus species, which is much lower than the hundreds of viral species hypothesized to be present by previous studies (137–139). This finding highlights the importance of considering evidence of virus activity when characterizing human viromes, where many detected “viruses” may actually be cryptic (or uninducible) prophages. We observed generally low virus prevalence across CF respiratory tract samples (Figure 5B), indicating that in general, each person with CF has a unique respiratory tract virome. When comparing viruses detected in upper respiratory tract samples (oropharyngeal and nasopharyngeal swabs) to those identified in lower respiratory tract samples (sputum/BAL), we observed that filamentous phages were more prevalent in the upper respiratory tract samples (Figure 5C), phages infecting Streptococcaceae, Pseudomonadaceae, and Staphylococcaceae were more prevalent in the lower respiratory tract samples (Figure 5D), and temperate viruses were dominant in both sample types (Figure 5E).

When analyzing phage-bacterial dynamics using phage-to-host (PTH) ratios, we observed that most phages existed at a 1:1 ratio with their bacterial host. This observation aligns with our finding that most viruses detected in the CF respiratory tract have a temperate lifestyle (Figure 5E) and supports the hypothesis that phage activity is generally low in human microbiomes (140). Low PTH ratios were also commonly observed among phages infecting conventional CF pathogens, although *Pseudomonas*, *Burkholderia*, and *Stenotrophomonas* phages occasionally exhibited PTH ratios ≥ 2 (Figure 6A). Notably, many (40.6%) phage species with PTH ratios ≥ 2 were filamentous phages, which may not lyse their host cells upon replication. When observed longitudinally, PTH ratios were very consistent (Figure 6B), even during periods of antibiotic treatment (Figure 6C). Together, our PTH analyses indicate that phage “blooms”, or high levels of induction/lytic activity, are uncommon in CF respiratory tract samples, even among phages infecting CF pathogens.

While these results do not directly show that highly-active, virulent phages can’t persist in CF respiratory tract samples, these results do raise questions about the potential efficacy, activity, and persistence of such phages when used as therapeutics in this context.

In an ongoing effort to keep UHVDB current, we updated this resource with the most recent human virome databases, some of which were released during the writing of this manuscript, illustrating the practical utility of an updatable framework (Figure 7A-C).

Additionally, we expanded the breadth of UHVDB as a human virome database by including human-infecting RNA viruses, which enables users to perform complete human virome profiling (human DNA/RNA viruses and phages) with a consistent, standardized framework (Figure 7D-E).

While UHVDB presents numerous new advantages and capabilities, this resource has several limitations. First, the increased stringency as a result of the quality and confidence filters the UHVDB toolkit imposes compared with other tools likely comes with decreased sensitivity for identifying highly novel/divergent viruses. Second, although UHVDB r6 does include RNA viruses, there has not been a significant effort to mine these sequences other than what has been submitted to NCBI, highlighting metatranscriptomes as an important area of emphasis for future virus identification efforts. Third, UHVDB’s mining protocol classifies sequences exclusively as virus or non-virus, a categorization that can be complicated for non-canonical entities, such as phage-plasmids, that have both viral and non-viral features and were treated as plasmid in our analysis (141). Finally, our classifier for identifying uninducible prophages was trained on a relatively small dataset from infant fecal samples. The identification or creation of another training dataset from different sample types would very likely improve our ability to distinguish between active/inducible viruses and uninducible prophages and should be considered a high priority.

In summary, UHVDB and the associated toolkit enable standardization and updatability in human virome analyses. The impact of this resource will largely depend on community adoption. However, the current study provides findings and illustrates concepts showing the utility of our strategy and potential benefits to users. Specifically: 1) We demonstrate the value of a unified, updateable database and profiling technique for human virome analyses. 2) We identify minimal overlap in human viromes across body sites. 3) We demonstrate the utility of phage-to-host ratios as a measure of virus activity. 4) We find that distinguishing active viruses from uninducible prophages decreases the number active viruses detected in human metagenomes. 5) We find that many of the viruses we detected in human microbiota are likely low-activity, temperate phages. 6) We show that sequence-based virus profiling can provide higher-resolution insights into the contents of a metagenome, beyond enumerating viruses by taxonomic label alone.

## ACKNOWLEDGEMENTS

We would like to acknowledge Stephen Nayfach for valuable discussions regarding virus mining and the methods involved in the creation of UHGV. We would also like to acknowledge Jeffrey McLean and Kristoffer Kerns for valuable discussions surrounding the analysis of viruses and bacteria in parallel, and Linda Kalikin for her help in identifying and shipping clinical samples. We would also like to acknowledge the University of Washington Microbial Interactions and Microbiome Center (mim_c) for technical and practical support.

## Artificial Intelligence Statement

Artificial intelligence was used as an aid for generating some of the figures in this manuscript.

## AUTHOR CONTRIBUTIONS

- Carson J. Miller (Conceptualization [lead], Data curation [lead], Formal analysis [lead], Investigation [lead], Methodology [lead], Software [lead], Validation [lead], Visualization [lead], Writing - original draft [lead], Writing - review & editing [equal])
- Christopher E. Pope (Conceptualization [supporting], Funding acquisition [supporting])
- Margot L. Lavitt (Methodology [supporting])
- Lindsay J. Caverly (Funding acquisition [supporting], Resources [supporting])
- John J. LiPuma (Funding acquisition [supporting], Resources [supporting], Writing - review & editing [supporting])
- Kelsi Penewit (Methodology [supporting])
- Janessa D. Lewis (Methodology [supporting])
- Stephen J. Salipante (Funding acquisition [supporting], Methodology [supporting], Resources [supporting], Writing - review & editing [supporting])
- Lucas R. Hoffman (Conceptualization [supporting], Funding acquisition [lead], Project Administration [lead], Resources [lead], Supervision [lead], Writing - review & editing [equal])

## SUPPLEMENTARY DATA

Supplementary Data are available at NAR online.

## DATA AVAILABILITY

Sequence reads generated in support of this study are accessible from the SRA (Bioproject PRJNA1457048) and will be released upon publication. UHVDB releases, including the latest r6 release, can be found at: https://zenodo.org/communities/uhvdb/

## CODE AVAILABILITY

The UHVDB toolkit, which was used to create UHVDB and analyze the 1,983 CF metagenomes, is publicly available at https://github.com/UHVDB/toolkit. All other code relating to this manuscript is available in the form of Jupyter notebooks at https://github.com/UHVDB/uhvdb-manuscript.

## REFERENCES

1. Suttle, C.A. (2005) Viruses in the sea. Nature, 437, 356–361.

2. Brüssow, H. and Hendrix, R.W. (2002) Phage genomics: small is beautiful. Cell, 108, 13–16.

3. Koonin, E.V., Dolja, V.V., Krupovic, M., Varsani, A., Wolf, Y.I., Yutin, N., Zerbini, F.M. and Kuhn, J.H. (2020) Global Organization and Proposed Megataxonomy of the Virus World. Microbiol Mol Biol Rev, 84.

4. Koonin, E.V., Krupovic, M. and Dolja, V.V. (2023) The global virome: How much diversity and how many independent origins? Environ Microbiol, 25, 40–44.

5. Piel, D., Bruto, M., Labreuche, Y., Blanquart, F., Goudenège, D., Barcia-Cruz, R., Chenivesse, S., Le Panse, S., James, A., Dubert, J., et al. (2022) Phage-host coevolution in natural populations. Nat Microbiol, 7, 1075–1086.

6. Breitbart, M., Bonnain, C., Malki, K. and Sawaya, N.A. (2018) Phage puppet masters of the marine microbial realm. Nat Microbiol, 3, 754–766.

7. Williamson, K.E., Fuhrmann, J.J., Wommack, K.E. and Radosevich, M. (2017) Viruses in Soil Ecosystems: An Unknown Quantity Within an Unexplored Territory. Annu Rev Virol, 4, 201–219.

8. Liang, G. and Bushman, F.D. (2021) The human virome: assembly, composition and host interactions. Nature Reviews Microbiology, 19, 514–527.

9. Champagne-Jorgensen, K., Luong, T., Darby, T. and Roach, D.R. (2023) Immunogenicity of bacteriophages. Trends Microbiol, 31, 1058–1071.

10. Zamora, P.F., Reidy, T.G., Armbruster, C.R., Sun, M., Van Tyne, D., Turner, P.E., Koff, J.L. and Bomberger, J.M. (2024) Lytic bacteriophages induce the secretion of antiviral and proinflammatory cytokines from human respiratory epithelial cells. PLoS Biol, 22, e3002566.

11. Borodovich, T., Shkoporov, A.N., Ross, R.P. and Hill, C. (2022) Phage-mediated horizontal gene transfer and its implications for the human gut microbiome. Gastroenterol Rep (Oxf), 10, goac012.

12. Bae, T., Baba, T., Hiramatsu, K. and Schneewind, O. (2006) Prophages of Staphylococcus aureus Newman and their contribution to virulence. Molecular Microbiology, 62, 1035–1047.

13. Kortright, K.E., Chan, B.K., Koff, J.L. and Turner, P.E. (2019) Phage Therapy: A Renewed Approach to Combat Antibiotic-Resistant Bacteria. Cell Host Microbe, 25, 219–232.

14. Secor, P.R., Sweere, J.M., Michaels, L.A., Malkovskiy, A.V., Lazzareschi, D., Katznelson, E., Rajadas, J., Birnbaum, M.E., Arrigoni, A., Braun, K.R., et al. (2015) Filamentous Bacteriophage Promote Biofilm Assembly and Function. Cell Host Microbe, 18, 549–559.

15. The Diverse Impacts of Phage Morons on Bacterial Fitness and Virulence (2019) In Advances in Virus Research. Academic Press, Vol. 103, pp. 1–31.

16. Schmidt, T.S.B., Fullam, A., Ferretti, P., Orakov, A., Maistrenko, O.M., Ruscheweyh, H.-J., Letunic, I., Duan, Y., Van Rossum, T., Sunagawa, S., et al. (2024) SPIRE: a Searchable, Planetary-scale mIcrobiome REsource. Nucleic Acids Res, 52, D777–D783.

17. Website 10.1101/2024.03.08.584059.

18. Dmitrijeva, M., Ruscheweyh, H.-J., Feer, L., Li, K., Miravet-Verde, S., Sintsova, A., Mende, D.R., Zeller, G. and Sunagawa, S. (2025) The mOTUs online database provides web-accessible genomic context to taxonomic profiling of microbial communities. Nucleic Acids Res, 53, D797–D805.

19. Parks, D.H., Rinke, C., Chuvochina, M., Chaumeil, P.-A., Woodcroft, B.J., Evans, P.N., Hugenholtz, P. and Tyson, G.W. (2017) Recovery of nearly 8, 000 metagenome-assembled genomes substantially expands the tree of life. Nature Microbiology, 2, 1533–1542.

20. Blanco-Míguez, A., Beghini, F., Cumbo, F., McIver, L.J., Thompson, K.N., Zolfo, M., Manghi, P., Dubois, L., Huang, K.D., Thomas, A.M., et al. (2023) Extending and improving metagenomic taxonomic profiling with uncharacterized species using MetaPhlAn 4. Nat Biotechnol, 41, 1633–1644.

21. Camargo, A.P., Nayfach, S., Chen, I.-M.A., Palaniappan, K., Ratner, A., Chu, K., Ritter, S.J., Reddy, T.B.K., Mukherjee, S., Schulz, F., et al. (2023) IMG/VR v4: an expanded database of uncultivated virus genomes within a framework of extensive functional, taxonomic, and ecological metadata. Nucleic Acids Res, 51, D733–D743.

22. Ren, J., Ahlgren, N.A., Lu, Y.Y., Fuhrman, J.A. and Sun, F. (2017) VirFinder: a novel k-mer based tool for identifying viral sequences from assembled metagenomic data. Microbiome, 5, 1–20.

23. Roux, S., Enault, F., Hurwitz, B.L. and Sullivan, M.B. (2015) VirSorter: mining viral signal from microbial genomic data. PeerJ, 3, e985.

24. Paez-Espino, D., Chen, I.-M.A., Palaniappan, K., Ratner, A., Chu, K., Szeto, E., Pillay, M., Huang, J., Markowitz, V.M., Nielsen, T., et al. (2016) IMG/VR: a database of cultured and uncultured DNA Viruses and retroviruses. Nucleic Acids Res, 45, gkw1030.

25. Camargo, A.P., Roux, S., Schulz, F., Babinski, M., Xu, Y., Hu, B., Chain, P.S.G., Nayfach, S. and Kyrpides, N.C. (2024) Identification of mobile genetic elements with geNomad. Nat Biotechnol, 42, 1303–1312.

26. Guo, J., Bolduc, B., Zayed, A.A., Varsani, A., Dominguez-Huerta, G., Delmont, T.O., Pratama, A.A., Gazitúa, M.C., Vik, D., Sullivan, M.B., et al. (2021) VirSorter2: a multi-classifier, expert-guided approach to detect diverse DNA and RNA viruses. Microbiome, 9, 37.

27. Camarillo-Guerrero, L.F., Almeida, A., Rangel-Pineros, G., Finn, R.D. and Lawley, T.D. (2021) Massive expansion of human gut bacteriophage diversity. Cell, 184, 1098–1109.e9.

28. Lai, S., Jia, L., Subramanian, B., Pan, S., Zhang, J., Dong, Y., Chen, W.-H. and Zhao, X.-M. (2020) mMGE: a database for human metagenomic extrachromosomal mobile genetic elements. Nucleic Acids Res, 49, D783–D791.

29. Pratama, A.A., Bolduc, B., Zayed, A.A., Zhong, Z.-P., Guo, J., Vik, D.R., Gazitúa, M.C., Wainaina, J.M., Roux, S. and Sullivan, M.B. (2021) Expanding standards in viromics: in silico evaluation of dsDNA viral genome identification, classification, and auxiliary metabolic gene curation. PeerJ, 9, e11447.

30. Roux, S., Emerson, J.B., Eloe-Fadrosh, E.A. and Sullivan, M.B. (2017) Benchmarking viromics: an evaluation of metagenome-enabled estimates of viral community composition and diversity. PeerJ, 5, e3817.

31. Roux, S., Camargo, A.P., Coutinho, F.H., Dabdoub, S.M., Dutilh, B.E., Nayfach, S. and Tritt, A. (2023) iPHoP: An integrated machine learning framework to maximize host prediction for metagenome-derived viruses of archaea and bacteria. PLOS Biology, 21, e3002083.

32. Nayfach, S., Páez-Espino, D., Call, L., Low, S.J., Sberro, H., Ivanova, N.N., Proal, A.D., Fischbach, M.A., Bhatt, A.S., Hugenholtz, P., et al. (2021) Metagenomic compendium of 189, 680 DNA viruses from the human gut microbiome. Nat Microbiol, 6, 960–970.

33. Galperina, A., Lugli, G.A., Milani, C., De Vos, W.M., Ventura, M., Salonen, A., Hurwitz, B. and Ponsero, A.J. (2025) The Aggregated Gut Viral Catalogue (AVrC): A unified resource for exploring the viral diversity of the human gut. PLOS Computational Biology, 21, e1012268.

34. GitHub - snayfach/UHGV: Unified Human Gut Virome Catalog GitHub.

35. Rangel-Pineros, G., Millard, A., Michniewski, S., Scanlan, D., Sirén, K., Reyes, A., Petersen, B., Clokie, M.R.J. and Sicheritz-Pontén, T. (2021) From Trees to Clouds: PhageClouds for Fast Comparison of ∼640, 000 Phage Genomic Sequences and Host-Centric Visualization Using Genomic Network Graphs. PHAGE, 10.1089/phage.2021.0008.

36. Roux, S., Páez-Espino, D., Chen, I.-M.A., Palaniappan, K., Ratner, A., Chu, K., Reddy, T.B.K., Nayfach, S., Schulz, F., Call, L., et al. (2020) IMG/VR v3: an integrated ecological and evolutionary framework for interrogating genomes of uncultivated viruses. Nucleic Acids Res, 49, D764–D775.

37. Wang, R.H., Yang, S., Liu, Z., Zhang, Y., Wang, X., Xu, Z., Wang, J. and Li, S.C. (2023) PhageScope: a well-annotated bacteriophage database with automatic analyses and visualizations. Nucleic Acids Res, 52, D756–D761.

38. Camargo, A.P., Baltoumas, F.A., Ndela, E.O., Fiamenghi, M.B., Merrill, B.D., Carter, M.M., Pinto, Y., Chakraborty, M., Andreeva, A., Ghiotto, G., et al. (2025) A genomic atlas of the human gut virome elucidates genetic factors shaping host interactions. bioRxivorg, 10.1101/2025.11.01.686033.

39. Dahlman, S., Avellaneda-Franco, L., Rutten, E.L., Gulliver, E.L., Solari, S., Chonwerawong, M., Kett, C., Subedi, D., Young, R.B., Campbell, N., et al. (2025) Isolation, engineering and ecology of temperate phages from the human gut. Nature, 647, 698–705.

40. Wirbel, J., Hickey, A.S., Chang, D., Enright, N.J., Dvorak, M., Chanin, R.B., Schmidtke, D.T. and Bhatt, A.S. (2025) Long-read metagenomics reveals phage dynamics in the human gut microbiome. Nature, 649, 982–990.

41. Bobay, L.-M., Touchon, M. and Rocha, E.P.C. (2014) Pervasive domestication of defective prophages by bacteria. Proc Natl Acad Sci U S A, 111, 12127–12132.

42. Nayfach, S., Camargo, A.P., Schulz, F., Eloe-Fadrosh, E., Roux, S. and Kyrpides, N.C. (2021) CheckV assesses the quality and completeness of metagenome-assembled viral genomes. Nat Biotechnol, 39, 578–585.

43. Antipov, D., Raiko, M., Lapidus, A. and Pevzner, P.A. (2020) Metaviral SPAdes: assembly of viruses from metagenomic data. Bioinformatics, 36, 4126–4129.

44. Zielezinski, A., Gudyś, A., Barylski, J., Siminski, K., Rozwalak, P., Dutilh, B.E. and Deorowicz, S. (2025) Ultrafast and accurate sequence alignment and clustering of viral genomes. Nat Methods, 22, 1191–1194.

45. Roux, S., Adriaenssens, E.M., Dutilh, B.E., Koonin, E.V., Kropinski, A.M., Krupovic, M., Kuhn, J.H., Lavigne, R., Brister, J.R., Varsani, A., et al. (2019) Minimum Information about an Uncultivated Virus Genome (MIUViG). Nat Biotechnol, 37, 29–37.

46. Eddy, S.R. (2011) Accelerated Profile HMM Searches. PLOS Computational Biology, 7, e1002195.

47. GitHub - apcamargo/tr-trimmer: Identify and trim terminal repeats from sequences in FASTA files GitHub.

48. GitHub - apcamargo/seq-hasher: Compute hash digests for DNA sequences in a FASTA file GitHub.

49. Aldeguer-Riquelme, B., Conrad, R.E., Antón, J., Rossello-Mora, R. and Konstantinidis, K.T. (2024) A natural ANI gap that can define intra-species units of bacteriophages and other viruses. mBio, 15, e0153624.

50. Van Dongen, S. (2008) Graph Clustering Via a Discrete Uncoupling Process. SIAM Journal on Matrix Analysis and Applications, 10.1137/040608635.

51. Hyatt, D., Chen, G.-L., Locascio, P.F., Land, M.L., Larimer, F.W. and Hauser, L.J. (2010) Prodigal: prokaryotic gene recognition and translation initiation site identification. BMC Bioinformatics, 11, 119.

52. Larralde, M. (2022) Pyrodigal: Python bindings and interface to Prodigal, an efficient method for gene prediction in prokaryotes. Journal of Open Source Software, 7, 4296.

53. Buchfink, B., Reuter, K. and Drost, H.-G. (2021) Sensitive protein alignments at tree-of-life scale using DIAMOND. Nat Methods, 18, 366–368.

54. Zielezinski, A., Deorowicz, S. and Gudyś, A. (2022) PHIST: fast and accurate prediction of prokaryotic hosts from metagenomic viral sequences. Bioinformatics, 38, 1447–1449.

55. Nishijima, S., Fullam, A., Schmidt, T.S.B., Kuhn, M. and Bork, P. (2026) VIRE: a metagenome-derived, planetary-scale virome resource with environmental context. Nucleic Acids Res., 54, D902–D911.

56. Roux, S., Neri, U., Bushnell, B., Fremin, B., George, N.A., Gophna, U., Hug, L.A., Camargo, A.P., Wu, D., Ivanova, N., et al. (2025) Planetary-scale metagenomic search reveals new patterns of CRISPR targeting. bioRxiv, 10.1101/2025.06.12.659409.

57. Shen, W., Le, S., Li, Y. and Hu, F. (2016) SeqKit: A Cross-Platform and Ultrafast Toolkit for FASTA/Q File Manipulation. PLoS One, 11, e0163962.

58. The UniProt Consortium, Bateman, A., Martin, M.-J., Orchard, S., Magrane, M., Adesina, A., Ahmad, S., Bowler-Barnett, E.H., Bye-A-Jee, H., Carpentier, D., et al. (2024) UniProt: the Universal Protein Knowledgebase in 2025. Nucleic Acids Res, 53, D609–D617.

59. Blum, M., Andreeva, A., Florentino, L.C., Chuguransky, S.R., Grego, T., Hobbs, E., Pinto, B.L., Orr, A., Paysan-Lafosse, T., Ponamareva, I., et al. (2024) InterPro: the protein sequence classification resource in 2025. Nucleic Acids Res, 53, D444–D456.

60. Schwengers, O., Jelonek, L., Dieckmann, M.A., Beyvers, S., Blom, J. and Goesmann, A. (2021) Bakta: rapid and standardized annotation of bacterial genomes via alignment-free sequence identification. Microb Genom, 7.

61. Suzek, B.E., Huang, H., McGarvey, P., Mazumder, R. and Wu, C.H. (2007) UniRef: comprehensive and non-redundant UniProt reference clusters. Bioinformatics, 23, 1282–1288.

62. van Kempen, M., Kim, S.S., Tumescheit, C., Mirdita, M., Lee, J., Gilchrist, C.L.M., Söding, J. and Steinegger, M. (2023) Fast and accurate protein structure search with Foldseek. Nature Biotechnology, 42, 243–246.

63. Kim, R.S., Levy Karin, E., Mirdita, M., Chikhi, R. and Steinegger, M. (2025) BFVD-a large repository of predicted viral protein structures. Nucleic Acids Res, 53, D340–D347.

64. Odai, R., Leemann, M., Al-Murad, T., Abdullah, M., Shyrokova, L., Tenson, T., Hauryliuk, V., Durairaj, J., Pereira, J. and Atkinson, G.C. (2025) The Viral AlphaFold Database of monomers and homodimers reveals conserved protein folds in viruses of bacteria, archaea, and eukaryotes. Sci Adv, 11, eadz8560.

65. Trgovec-Greif, L., Hellinger, H.-J., Mainguy, J., Pfundner, A., Frishman, D., Kiening, M., Webster, N.S., Laffy, P.W., Feichtinger, M. and Rattei, T. (2024) VOGDB-Database of Virus Orthologous Groups. Viruses, 16.

66. Jones, P., Binns, D., Chang, H.-Y., Fraser, M., Li, W., McAnulla, C., McWilliam, H., Maslen, J., Mitchell, A., Nuka, G., et al. (2014) InterProScan 5: genome-scale protein function classification. Bioinformatics, 30, 1236–1240.

67. Bouras, G., Nepal, R., Houtak, G., Psaltis, A.J., Wormald, P.-J. and Vreugde, S. (2023) Pharokka: a fast scalable bacteriophage annotation tool. Bioinformatics, 39.

68. Bouras, G., Grigson, S.R., Mirdita, M., Heinzinger, M., Papudeshi, B., Mallawaarachchi, V., Green, R., Kim, R.S., Mihalia, V., Psaltis, A.J., et al. (2026) Protein structure-informed bacteriophage genome annotation with Phold. Nucleic Acids Res, 54.

69. Boulay, A., Leprince, A., Enault, F., Rousseau, E. and Galiez, C. (2025) Empathi: embedding-based phage protein annotation tool by hierarchical assignment. Nat Commun, 16, 9114.

70. Enault, F., Briet, A., Bouteille, L., Roux, S., Sullivan, M.B. and Petit, M.-A. (2017) Phages rarely encode antibiotic resistance genes: a cautionary tale for virome analyses. ISME J., 11, 237–247.

71. Alcock, B.P., Huynh, W., Chalil, R., Smith, K.W., Raphenya, A.R., Wlodarski, M.A., Edalatmand, A., Petkau, A., Syed, S.A., Tsang, K.K., et al. (2023) CARD 2023: expanded curation, support for machine learning, and resistome prediction at the Comprehensive Antibiotic Resistance Database. Nucleic Acids Res, 51, D690–D699.

72. Zhou, S., Liu, B., Zheng, D., Chen, L. and Yang, J. (2025) VFDB 2025: an integrated resource for exploring anti-virulence compounds. Nucleic Acids Res, 53, D871–D877.

73. Hockenberry, A.J. and Wilke, C.O. (2021) BACPHLIP: predicting bacteriophage lifestyle from conserved protein domains. PeerJ, 9, e11396.

74. Tisza, M.J. and Buck, C.B. (2021) A catalog of tens of thousands of viruses from human metagenomes reveals hidden associations with chronic diseases. Proc Natl Acad Sci U S A, 118.

75. Li, S., Guo, R., Zhang, Y., Li, P., Chen, F., Wang, X., Li, J., Jie, Z., Lv, Q., Jin, H., et al. (2022) A catalog of 48, 425 nonredundant viruses from oral metagenomes expands the horizon of the human oral virome. iScience, 25, 104418.

76. Jie, Z., Liang, H., Meng, Y., Zhang, J., Zhang, T., Li, W., Lin, X., Hu, T., Han, M., Liang, W., et al. (2025) Integrating metagenomics and cultivation unveils oral phage diversity and potential impact on hosts. NPJ Biofilms Microbiomes, 11, 145.

77. Yan, Q., Huang, L., Li, S., Zhang, Y., Guo, R., Zhang, P., Lei, Z., Lv, Q., Chen, F., Li, Z., et al. (2025) The Chinese gut virus catalogue reveals gut virome diversity and disease-related viral signatures. Genome Med, 17, 30.

78. Wang, X., Dong, Q., Huang, P., Yang, S., Gao, M., Zhang, C., Zhang, C., Deng, Y., Huang, Z., Ma, B., et al. (2025) The genetic diversity and populational specificity of the human gut virome at single-nucleotide resolution. Microbiome, 13, 188.

79. Saheb Kashaf, S., Proctor, D.M., Deming, C., Saary, P., Hölzer, M., NISC Comparative Sequencing Program, Taylor, M.E., Kong, H.H., Segre, J.A., Almeida, A., et al. (2022) Integrating cultivation and metagenomics for a multi-kingdom view of skin microbiome diversity and functions. Nat Microbiol, 7, 169–179.

80. Huang, L., Guo, R., Li, S., Wu, X., Zhang, Y., Guo, S., Lv, Y., Xiao, Z., Kang, J., Meng, J., et al. (2024) A multi-kingdom collection of 33, 804 reference genomes for the human vaginal microbiome. Nat Microbiol, 9, 2185–2200.

81. Richardson, L., Allen, B., Baldi, G., Beracochea, M., Bileschi, M.L., Burdett, T., Burgin, J., Caballero-Pérez, J., Cochrane, G., Colwell, L.J., et al. (2023) MGnify: the microbiome sequence data analysis resource in 2023. Nucleic Acids Res, 51, D753–D759.

82. Chikhi, R., Lemane, T., Loll-Krippleber, R., Montoliu-Nerin, M., Raffestin, B., Camargo, A.P., Miller, C.J., Fiamenghi, M.B., Agustinho, D.P., Majidian, S., et al. (2025) Logan: Planetary-scale genome assembly surveys life’s diversity. bioRxivorg, 10.1101/2024.07.30.605881.

83. Cook, R., Brown, N., Redgwell, T., Rihtman, B., Barnes, M., Clokie, M., Stekel, D.J., Hobman, J., Jones, M.A. and Millard, A. (2021) INfrastructure for a PHAge REference Database: Identification of Large-Scale Biases in the Current Collection of Cultured Phage Genomes. Phage (New Rochelle), 2, 214–223.

84. Altschul, S.F., Gish, W., Miller, W., Myers, E.W. and Lipman, D.J. (1990) Basic local alignment search tool. J Mol Biol, 215, 403–410.

85. Di Tommaso, P., Chatzou, M., Floden, E.W., Barja, P.P., Palumbo, E. and Notredame, C. (2017) Nextflow enables reproducible computational workflows. Nat. Biotechnol., 35, 316–319.

86. Ewels, P.A., Peltzer, A., Fillinger, S., Patel, H., Alneberg, J., Wilm, A., Garcia, M.U., Di Tommaso, P. and Nahnsen, S. (2020) The nf-core framework for community-curated bioinformatics pipelines. Nat. Biotechnol., 38, 276–278.

87. Shaw, J. and Yu, Y.W. (2024) Rapid species-level metagenome profiling and containment estimation with sylph. Nat Biotechnol, 10.1038/s41587-024-02412-y.

88. Parks, D.H., Chaumeil, P.-A., Mussig, A.J., Rinke, C., Chuvochina, M. and Hugenholtz, P. (2026) GTDB release 10: a complete and systematic taxonomy for 715 230 bacterial and 17 245 archaeal genomes. Nucleic Acids Res., 54, D743–D754.

89. Aroney, S.T.N., Newell, R.J.P., Nissen, J.N., Camargo, A.P., Tyson, G.W. and Woodcroft, B.J. (2025) CoverM: read alignment statistics for metagenomics. Bioinformatics, 41.

90. Gourlé, H., Karlsson-Lindsjö, O., Hayer, J. and Bongcam-Rudloff, E. (2018) Simulating Illumina metagenomic data with InSilicoSeq. Bioinformatics, 35, 521–522.

91. GitHub - biobakery/baqlava: Bioinformatic Application for Quantification and LAbeling of VirAl taxonomy GitHub.

92. Pinto, Y., Chakraborty, M., Jain, N. and Bhatt, A.S. (2024) Phage-inclusive profiling of human gut microbiomes with Phanta. Nat Biotechnol, 42, 651–662.

93. Tisza, M.J., Lloyd, R.E., Hoffman, K., Smith, D.P., Rewers, M., Javornik Cregeen, S.J. and Petrosino, J.F. (2025) Longitudinal phage-bacteria dynamics in the early life gut microbiome. Nat Microbiol, 10, 420–430.

94. Liang, G., Zhao, C., Zhang, H., Mattei, L., Sherrill-Mix, S., Bittinger, K., Kessler, L.R., Wu, G.D., Baldassano, R.N., DeRusso, P., et al. (2020) The stepwise assembly of the neonatal virome is modulated by breastfeeding. Nature, 581, 470–474.

95. Kieft, K. and Anantharaman, K. (2022) Deciphering Active Prophages from Metagenomes. mSystems, 7, e0008422.

96. Zünd, M., Ruscheweyh, H.-J., Field, C.M., Meyer, N., Cuenca, M., Hoces, D., Hardt, W.-D. and Sunagawa, S. (2021) High throughput sequencing provides exact genomic locations of inducible prophages and accurate phage-to-host ratios in gut microbial strains. Microbiome, 9, 77.

97. Pedregosa, F., Varoquaux, G., Gramfort, A., Michel, V., Thirion, B., Grisel, O., Blondel, M., Müller, A., Nothman, J., Louppe, G., et al. (2012) Scikit-learn: Machine Learning in Python. arXiv [cs.LG], 10.48550/arXiv.1201.0490.

98. Chen, J., Sun, C., Dong, Y., Jin, M., Lai, S., Jia, L., Zhao, X., Wang, H., Gao, N.L., Bork, P., et al. (2024) Efficient Recovery of Complete Gut Viral Genomes by Combined Short- and Long-Read Sequencing. Adv Sci (Weinh), 11, e2305818.

99. Zhao, J., Schloss, P.D., Kalikin, L.M., Carmody, L.A., Foster, B.K., Petrosino, J.F., Cavalcoli, J.D., VanDevanter, D.R., Murray, S., Li, J.Z., et al. (2012) Decade-long bacterial community dynamics in cystic fibrosis airways. Proc. Natl. Acad. Sci. U. S. A., 109, 5809–5814.

100. Widder, S., Carmody, L.A., Opron, K., Kalikin, L.M., Caverly, L.J. and LiPuma, J.J. (2024) Microbial community organization designates distinct pulmonary exacerbation types and predicts treatment outcome in cystic fibrosis. Nat. Commun., 15, 4889.

101. Nelson, M.T., Wolter, D.J., Eng, A., Weiss, E.J., Vo, A.T., Brittnacher, M.J., Hayden, H.S., Ravishankar, S., Bautista, G., Ratjen, A., et al. (2020) Maintenance tobramycin primarily affects untargeted bacteria in the CF sputum microbiome. Thorax, 75, 780–790.

102. Goldfarb, T., Kodali, V.K., Pujar, S., Brover, V., Robbertse, B., Farrell, C.M., Oh, D.-H., Astashyn, A., Ermolaeva, O., Haddad, D., et al. (2025) NCBI RefSeq: reference sequence standards through 25 years of curation and annotation. Nucleic Acids Res, 53, D243–D257.

103. Fiamenghi, M.B., Camargo, A.P., Chasapi, I.N., Baltoumas, F.A., Roux, S., Egorov, A.A., Aplakidou, E., Ndela, E.O., Vasquez, Y.M., Chen, I.-M.A., et al. (2026) Meta-virus resource (MetaVR): expanding the frontiers of viral diversity with 24 million uncultivated virus genomes. Nucleic Acids Res, 54, D801–D812.

104. Zenodo (2026) The human Oral and Airway Viral Genome Catalogue from metagenomes enables virome characterization informing respiratory health. 10.5281/ZENODO.15037348.

105. Peng, Y., Zhu, J., Wang, S., Liu, Y., Liu, X., DeLeon, O., Zhu, W., Xu, Z., Zhang, X., Zhao, S., et al. (2024) A metagenome-assembled genome inventory for children reveals early-life gut bacteriome and virome dynamics. Cell Host Microbe, 32, 2212–2230.e8.

106. Brister, J.R., Ako-Adjei, D., Bao, Y. and Blinkova, O. (2015) NCBI viral genomes resource. Nucleic Acids Res, 43, D571–7.

107. Hsieh, T.C., Ma, K.H. and Chao, A. (2016) iNEXT: an R package for rarefaction and extrapolation of species diversity (Hill numbers). Methods in Ecology and Evolution, 7, 1451–1456.

108. Rajagopala, S.V., Bakhoum, N.G., Pakala, S.B., Shilts, M.H., Rosas-Salazar, C., Mai, A., Boone, H.H., McHenry, R., Yooseph, S., Halasa, N., et al. (2021) Metatranscriptomics to characterize respiratory virome, microbiome, and host response directly from clinical samples. Cell Rep Methods, 1.

109. Glickman, C., Hendrix, J. and Strong, M. (2021) Simulation study and comparative evaluation of viral contiguous sequence identification tools. BMC Bioinformatics, 22, 329.

110. Jensen, J.S.L., Maharjan, S., Münch, P.C., Shen, J., Bowcutt, B., Sumner, J.T., Morgan, X.C., Thompson, K.N., Nguyen, L.H., Franzosa, E.A., et al. (2026) Enhanced multi-omic viral profiling from microbial community sequencing with BAQLaVa. bioRxivorg, 10.64898/2026.02.11.705346.

111. Dougherty, P.E., Bernard, C., Carstens, A.B., Bumunang, E., Gerovac, M., Müsken, M., Stanford, K., McAllister, T.A., Rocha, E.P.C. and Hansen, L.H. (2025) Persistent virulent phages exist across bacterial isolates. Nature Microbiology, 11, 31–41.

112. Perfilyev, A., Gæde, A., Hooton, S., Zahran, S.A., Kalatzis, P.G., Winther-Have, C.S., Ibarra Chavez, R., Wilkinson, R.C., Thanki, A.M., Liu, Z., et al. (2025) Large-scale analysis of bacterial genomes reveals thousands of lytic phages. Nature Microbiology, 11, 42–52.

113. Burgener, E.B., Sweere, J.M., Bach, M.S., Secor, P.R., Haddock, N., Jennings, L.K., Marvig, R.L., Johansen, H.K., Rossi, E., Cao, X., et al. (2019) Filamentous bacteriophages are associated with chronic Pseudomonas lung infections and antibiotic resistance in cystic fibrosis. Sci. Transl. Med., 11, eaau9748.

114. Cauwenberghs, E., De Boeck, I., Delanghe, L., Van Rillaer, T., Demuyser, T., Spacova, I., Verhulst, S., Van Hoorenbeeck, K. and Lebeer, S. (2025) Shallow metagenomic shotgun sequencing improves detection of pathogenic species in cystic fibrosis respiratory samples. J Cyst Fibros, 24, 909–915.

115. Shumyatsky, G., Burrell, A., Chaney, H., Sami, I., Koumbourlis, A.C., Freishtat, R.J., Crandall, K.A., Zemanick, E.T. and Hahn, A. (2022) Using metabolic potential within the airway microbiome as predictors of clinical state in persons with cystic fibrosis. Front Med (Lausanne), 9, 1082125.

116. Vieira, J., Jesudasen, S., Bringhurst, L., Sui, H.-Y., McIver, L., Whiteson, K., Hanselmann, K., O’Toole, G.A., Richards, C.J., Sicilian, L., et al. (2022) Supplemental Oxygen Alters the Airway Microbiome in Cystic Fibrosis. mSystems, 7, e0036422.

117. Bacci, G., Taccetti, G., Dolce, D., Armanini, F., Segata, N., Di Cesare, F., Lucidi, V., Fiscarelli, E., Morelli, P., Casciaro, R., et al. (2020) Untargeted metagenomic investigation of the airway microbiome of cystic fibrosis patients with moderate-severe lung disease. Microorganisms, 8, 1003.

118. Bacci, G., Mengoni, A., Fiscarelli, E., Segata, N., Taccetti, G., Dolce, D., Paganin, P., Morelli, P., Tuccio, V., De Alessandri, A., et al. (2017) A Different Microbiome Gene Repertoire in the Airways of Cystic Fibrosis Patients with Severe Lung Disease. Int J Mol Sci, 18.

119. Rossi, E., Lausen, M., Øbro, N.F., Colque, C.A., Nielsen, B.U., Møller, R., de Gier, C., Hald, A., Skov, M., Pressler, T., et al. (2024) Widespread alterations in systemic immune profile are linked to lung function heterogeneity and airway microbes in cystic fibrosis. J Cyst Fibros, 23, 885–895.

120. de Almeida, O.G.G., Capizzani, C.P. da C., Tonani, L., Grizante Barião, P.H., da Cunha, A.F., De Martinis, E.C.P., Torres, L.A.G.M.M. and von Zeska Kress, M.R. (2020) The Lung Microbiome of Three Young Brazilian Patients With Cystic Fibrosis Colonized by Fungi. Front Cell Infect Microbiol, 10, 598938.

121. Dmitrijeva, M., Kahlert, C.R., Feigelman, R., Kleiner, R.L., Nolte, O., Albrich, W.C., Baty, F. and von Mering, C. (2021) Strain-resolved dynamics of the lung microbiome in patients with cystic fibrosis. MBio, 12.

122. Pallenberg, S.T., Pust, M.-M., Rosenboom, I., Hansen, G., Wiehlmann, L., Dittrich, A.-M. and Tümmler, B. (2022) Impact of Elexacaftor/Tezacaftor/Ivacaftor Therapy on the Cystic Fibrosis Airway Microbial Metagenome. Microbiol Spectr, 10, e0145422.

123. Motta, H., Reuwsaat, J.C.V., Lopes, F.C., Viezzer, G., Volpato, F.C.Z., Barth, A.L., de Tarso Roth Dalcin, P., Staats, C.C., Vainstein, M.H. and Kmetzsch, L. (2024) Comparative microbiome analysis in cystic fibrosis and non-cystic fibrosis bronchiectasis. Respir Res, 25, 211.

124. Ghuneim, L.-A.J., Raghuvanshi, R., Neugebauer, K.A., Guzior, D.V., Christian, M.H., Schena, B., Feiner, J.M., Castillo-Bahena, A., Mielke, J., McClelland, M., et al. (2022) Complex and unexpected outcomes of antibiotic therapy against a polymicrobial infection. ISME J, 16, 2065–2075.

125. Inam, Z., Felton, E., Burrell, A., Chaney, H., Sami, I., Koumbourlis, A.C., Freishtat, R.J., Zemanick, E.T., Crandall, K.A. and Hahn, A. (2022) Impact of Antibiotics on the Lung Microbiome and Lung Function in Children With Cystic Fibrosis 1 Year After Hospitalization for an Initial Pulmonary Exacerbation. Open Forum Infect Dis, 9, ofac466.

126. Pust, M.-M., Wiehlmann, L., Davenport, C., Rudolf, I., Dittrich, A.-M. and Tümmler, B. (2020) The human respiratory tract microbial community structures in healthy and cystic fibrosis infants. NPJ Biofilms Microbiomes, 6, 61.

127. Mancabelli, L., Milani, C., Fontana, F., Lugli, G.A., Tarracchini, C., Turroni, F., van Sinderen, D. and Ventura, M. (2022) Mapping bacterial diversity and metabolic functionality of the human respiratory tract microbiome. J Oral Microbiol, 14, 2051336.

128. Pienkowska, K., Pust, M.-M., Gessner, M., Gaedcke, S., Thavarasa, A., Rosenboom, I., Morán Losada, P., Minso, R., Arnold, C., Hedtfeld, S., et al. (2023) The Cystic Fibrosis Upper and Lower Airway Metagenome. Microbiol Spectr, 11, e0363322.

129. Mirhakkak, M.H., Chen, X., Ni, Y., Heinekamp, T., Sae-Ong, T., Xu, L.-L., Kurzai, O., Barber, A.E., Brakhage, A.A., Boutin, S., et al. (2023) Genome-scale metabolic modeling of Aspergillus fumigatus strains reveals growth dependencies on the lung microbiome. Nat Commun, 14, 4369.

130. Mostacci, N., Wüthrich, T.M., Siegwald, L., Kieser, S., Steinberg, R., Sakwinska, O., Latzin, P., Korten, I. and Hilty, M. (2023) Informed interpretation of metagenomic data by StrainPhlAn enables strain retention analyses of the upper airway microbiome. mSystems, 8, e0072423.

131. Nelson, M.T., Pope, C.E., Marsh, R.L., Wolter, D.J., Weiss, E.J., Hager, K.R., Vo, A.T., Brittnacher, M.J., Radey, M.C., Hayden, H.S., et al. (2019) Human and Extracellular DNA Depletion for Metagenomic Analysis of Complex Clinical Infection Samples Yields Optimized Viable Microbiome Profiles. Cell Rep, 26, 2227–2240.e5.

132. Abbas, A.A., Taylor, L.J., Dothard, M.I., Leiby, J.S., Fitzgerald, A.S., Khatib, L.A., Collman, R.G. and Bushman, F.D. (2019) Redondoviridae, a Family of Small, Circular DNA Viruses of the Human Oro-Respiratory Tract Associated with Periodontitis and Critical Illness. Cell Host Microbe, 25, 719–729.e4.

133. Merenstein, C., Liang, G., Whiteside, S.A., Cobián-Güemes, A.G., Merlino, M.S., Taylor, L.J., Glascock, A., Bittinger, K., Tanes, C., Graham-Wooten, J., et al. (2021) Signatures of COVID-19 Severity and Immune Response in the Respiratory Tract Microbiome. mBio, 10.1128/mbio.01777-21.

134. Secor, P.R., Burgener, E.B., Kinnersley, M., Jennings, L.K., Roman-Cruz, V., Popescu, M., Van Belleghem, J.D., Haddock, N., Copeland, C., Michaels, L.A., et al. (2020) Pf bacteriophage and their impact on Pseudomonas virulence, mammalian immunity, and chronic infections. Front. Immunol., 11, 244.

135. Tisza, M.J., Pastrana, D.V., Welch, N.L., Stewart, B., Peretti, A., Starrett, G.J., Pang, Y.-Y.S., Krishnamurthy, S.R., Pesavento, P.A., McDermott, D.H., et al. (2020) Discovery of several thousand highly diverse circular DNA viruses. Elife, 9.

136. Gregory, A.C., Zablocki, O., Zayed, A.A., Howell, A., Bolduc, B. and Sullivan, M.B. (2020) The Gut Virome Database Reveals Age-Dependent Patterns of Virome Diversity in the Human Gut. Cell Host Microbe, 28, 724–740.e8.

137. Willner, D., Furlan, M., Haynes, M., Schmieder, R., Angly, F.E., Silva, J., Tammadoni, S., Nosrat, B., Conrad, D. and Rohwer, F. (2009) Metagenomic Analysis of Respiratory Tract DNA Viral Communities in Cystic Fibrosis and Non-Cystic Fibrosis Individuals. PLOS ONE, 4, e7370.

138. Willner, D., Haynes, M.R., Furlan, M., Hanson, N., Kirby, B., Lim, Y.W., Rainey, P.B., Schmieder, R., Youle, M., Conrad, D., et al. (2012) Case studies of the spatial heterogeneity of DNA viruses in the cystic fibrosis lung. Am. J. Respir. Cell Mol. Biol., 46, 127–131.

139. Lim, Y.W., Schmieder, R., Haynes, M., Willner, D., Furlan, M., Youle, M., Abbott, K., Edwards, R., Evangelista, J., Conrad, D., et al. (2013) Metagenomics and metatranscriptomics: windows on CF-associated viral and microbial communities. J. Cyst. Fibros., 12, 154–164.

140. Lopez, J.A., McKeithen-Mead, S., Shi, H., Nguyen, T.H., Huang, K.C. and Good, B.H. (2025) Abundance measurements reveal the balance between lysis and lysogeny in the human gut microbiome. Curr. Biol., 35, 2282–2294.e11.

141. Ilchenko, K., Bonnin, R.A., Rocha, E.P.C. and Pfeifer, E. (2026) Efficient detection and typing of phage-plasmids. MBio, 17, e0300025.

