## Supplementary Information for "The Updateable Human Virome Database and toolkit: A novel framework for human virome analysis"

**Supplementary Figures**

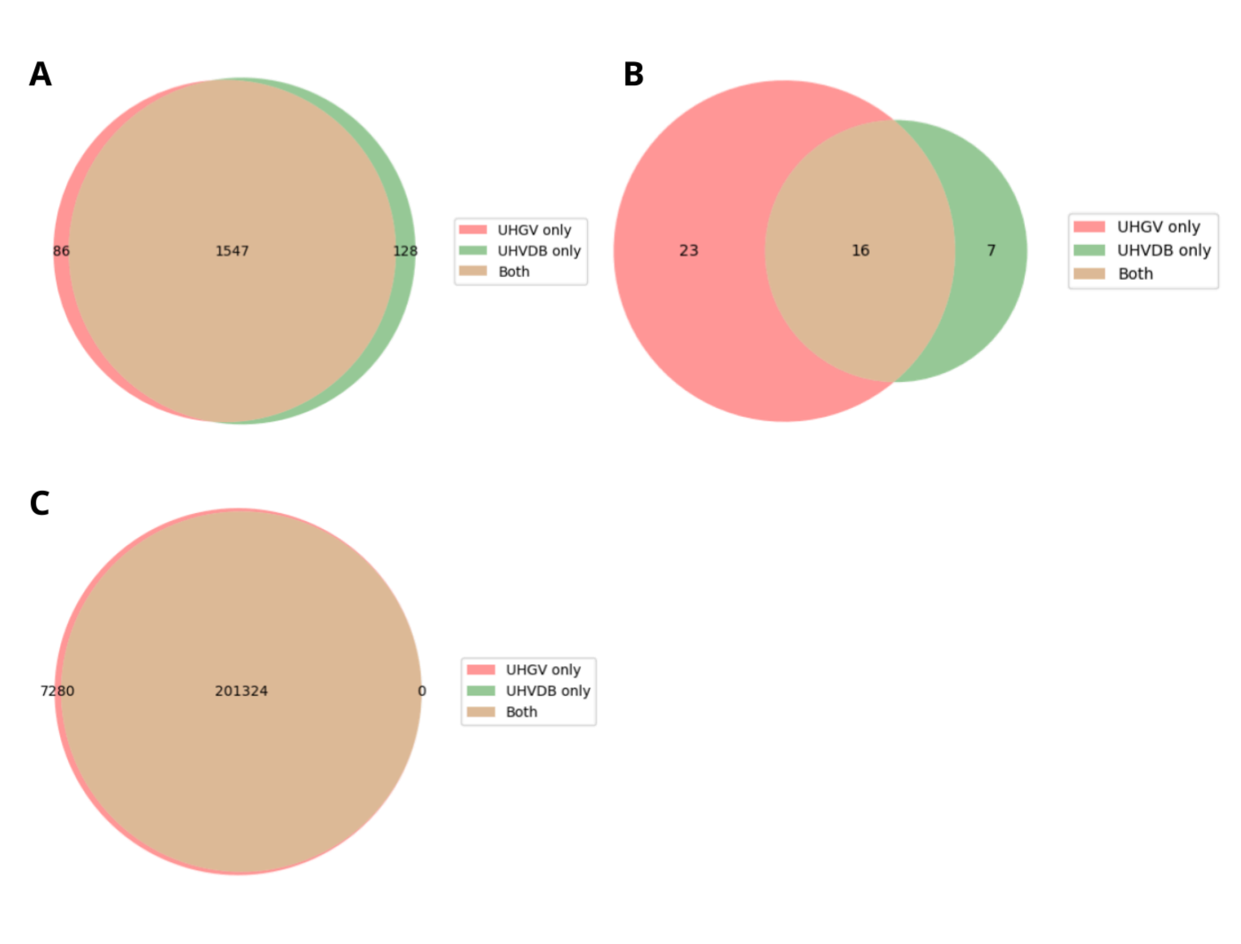


**Supplementary Figure S1.** Benchmarking UHVDB's virus mining workflow. Overlap between (**a**) confident and (**b**) uncertain viruses identified in the Danish Enteric Virome Catalog (DECoV) (142) subset of UHGV sequences using UHGV's workflow (*red*), UHVDB's workflow (*green*) or both workflows (*brown*). (**c**) Overlap of UHGV's high-quality viruses also classified as high-quality viruses with UHVDB's workflow. UHVDB yielded 98.2% agreement with UHGV's classification of the same sequences.

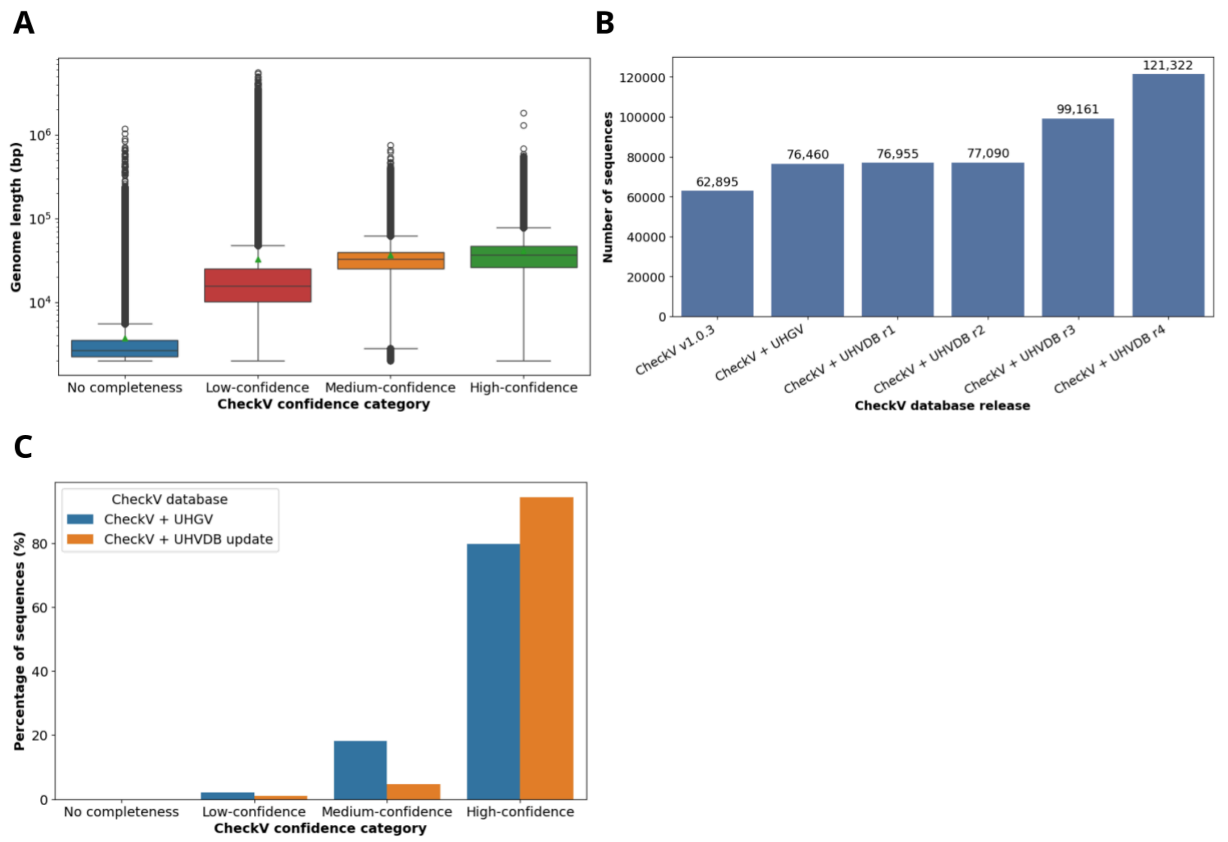


**Supplementary Figure S2.** Expanding CheckV's database of complete viruses. (**a**) Genome lengths of viruses having different levels of confidence in their CheckV predictions. Green triangles indicate mean values. (**b**) Count of sequences in CheckV's database after adding complete sequences from UHGV and subsequent UHVDB releases. (**c**) Percentage of UHVDB sequences receiving high, medium, and low confidence CheckV completeness estimates before (*blue*) and after (*orange*) updating CheckV with additional complete viruses.

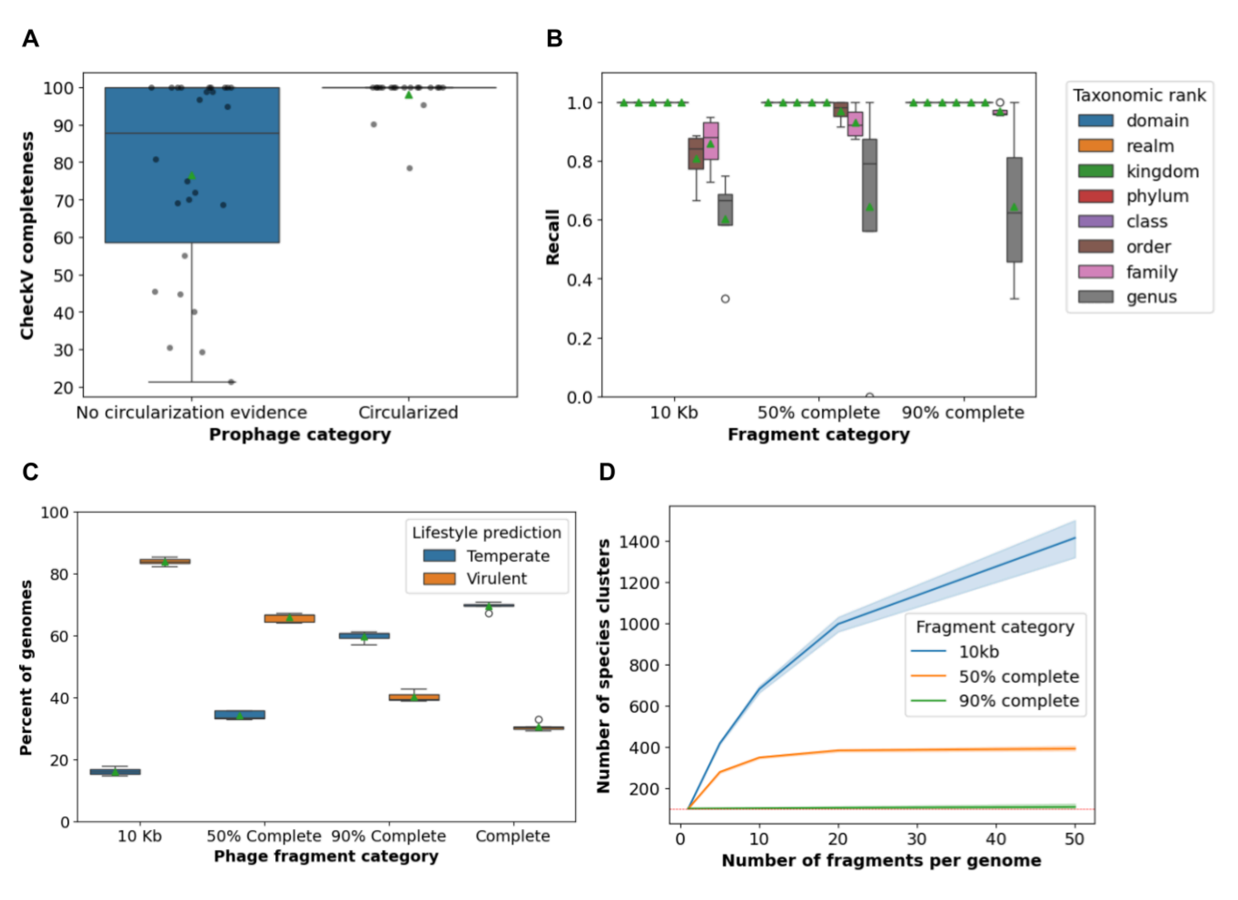


**Supplementary Figure S3.** The effect of genome fragmentation on virus induction, annotation, and clustering. (**a**) CheckV completeness of geNomad predicted prophages, based on whether they were circularized or not during Mitomycin C treatment. Only complete (circular) bacterial reference genomes were used for this analysis. (**b**) Recall (proportion of sequences receiving a taxonomic annotation) of taxonomic annotations at different ranks when complete virus genomes are fragmented at different levels *in silico.* The complete genomes from which the fragments were derived had annotations at all taxonomic ranks. (**c**) Percentage of genomes classified as temperate (*blue*) or virulent (*orange*) using BACPHLIP at different fragmentation levels. The fragments are derived from the complete genomes plotted on the right side of the graph. Green triangles indicate mean values for all box plots. (**d**) Count of species-level clusters yielded when 100 complete viruses (indicated by red dashed line) are clustered using an increasing number of fragments per genome at different fragmentation levels.

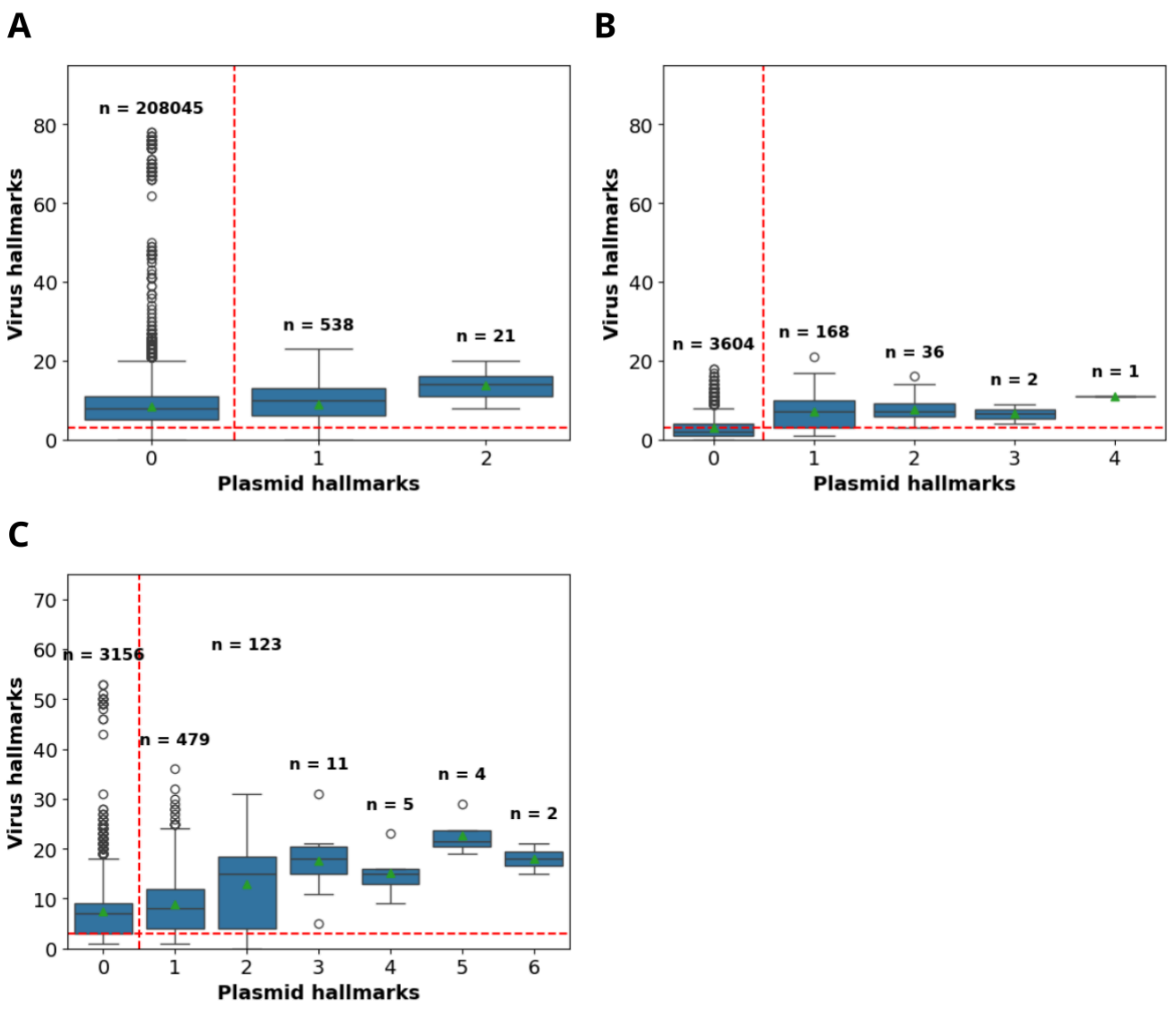


**Supplementary Figure S4.** Filtering uncertain genomes using geNomad hallmarks. Count of virus and plasmid hallmarks detected by MMSeqs2 among UHGV's high-quality (**a**) confident and (**b**) uncertain viruses. (**c**) Virus and plasmid hallmarks detected among UHGV's high-quality uncertain viruses using HMMER. Red dashed lines indicate the selected thresholds of ≥ 3 virus hallmarks and < 1 plasmid hallmark. Green triangles indicate mean values, with the counts displayed above each box.

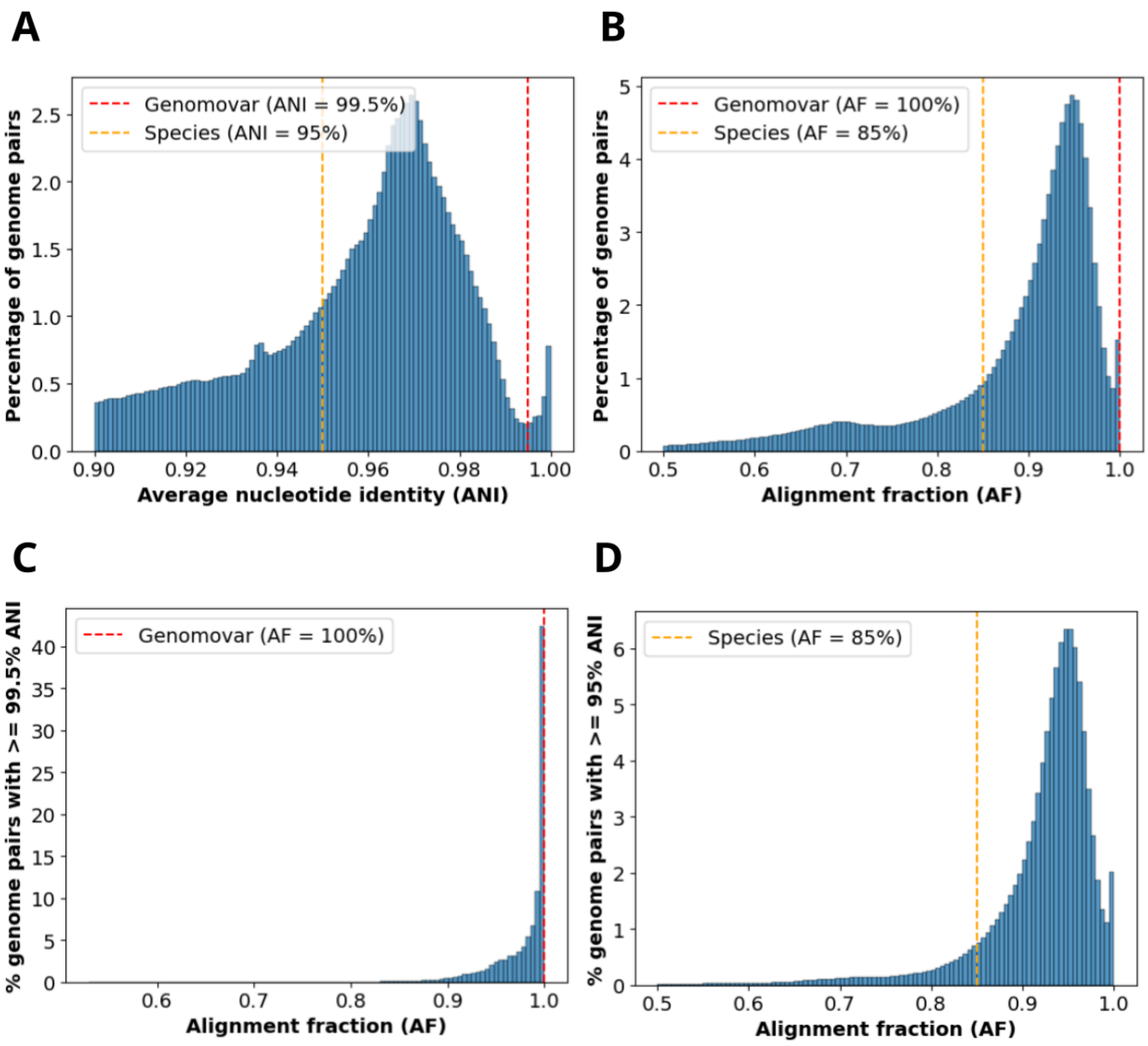


**Supplementary Figure S5.** Genomovar and species clustering thresholds. Percentage of genome pairs having (**a**) average nucleotide identities (ANI) and (**b**) alignment fractions (AF) falling into different bins. Distribution of AF for genome pairs having (**c**) ≥ 99.5% ANI and (**d**) ≥ 95% ANI.

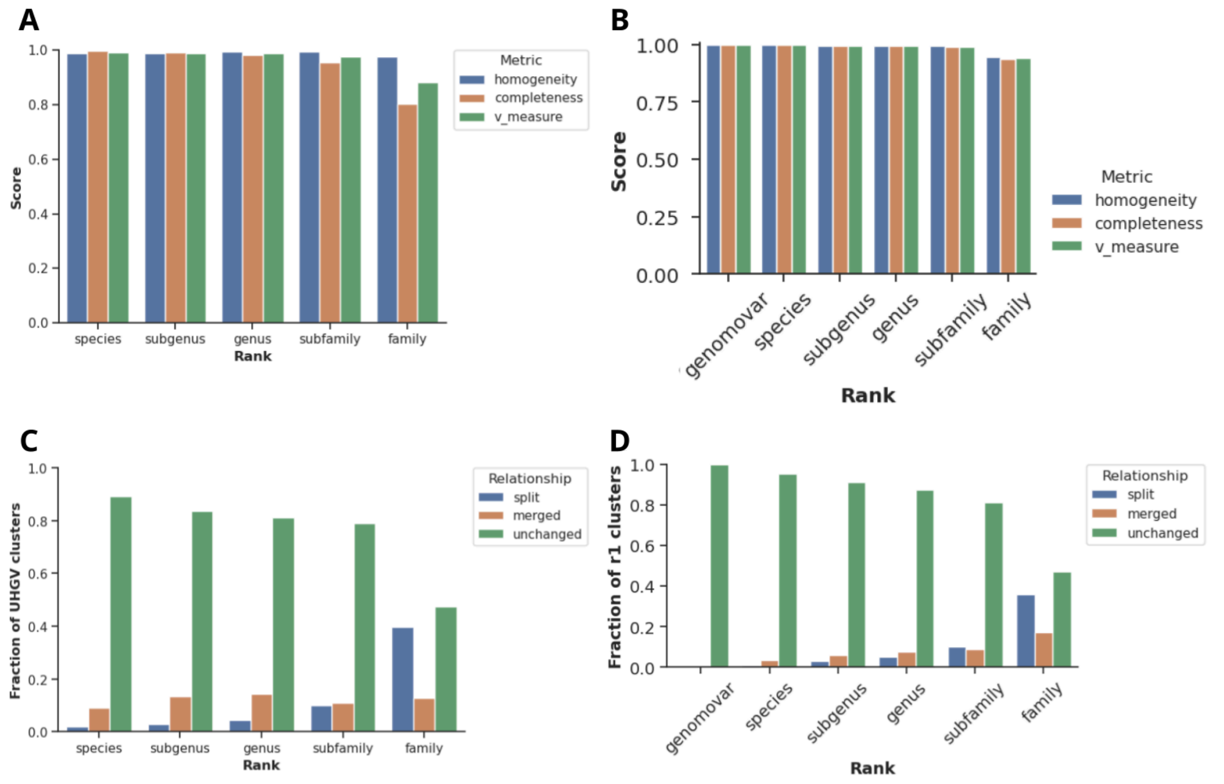


**Supplementary Figure S6.** UHVDB clustering benchmarks. Homogeneity, completeness, and v-measure when comparing (**a**) UHVDB r1 to UHGV clusters and (**b**) UHVDB release 1 (r1) to UHVDB r4 at different taxonomic ranks. The fraction of clusters that were split, merged, or unchanged at different taxonomic ranks when comparing (**c**) UHVDB r1 to UHGV and (**d**) UHVDB r1 to UHVDB release 4 (r4).

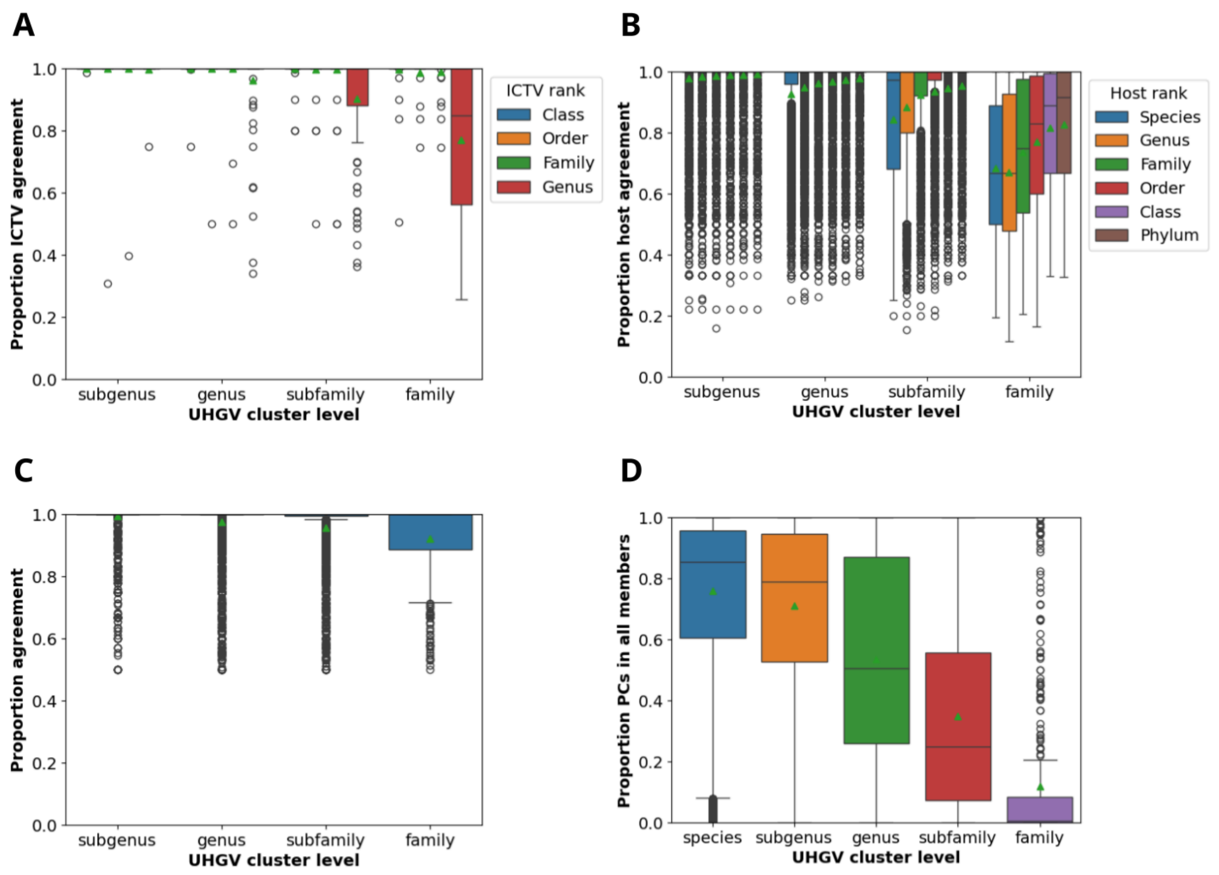


**Supplementary Figure S7.** Intra-cluster annotation consistency within UHGV high-quality clusters. Agreement in (**a**) ICTV taxonomy (**b**) predicted hosts and (**c**) lifestyle at different ranks within UHGV clusters of high-quality viruses. (**d**) Proportion of protein clusters shared by all members of a UHGV cluster. Green triangles indicate mean values.

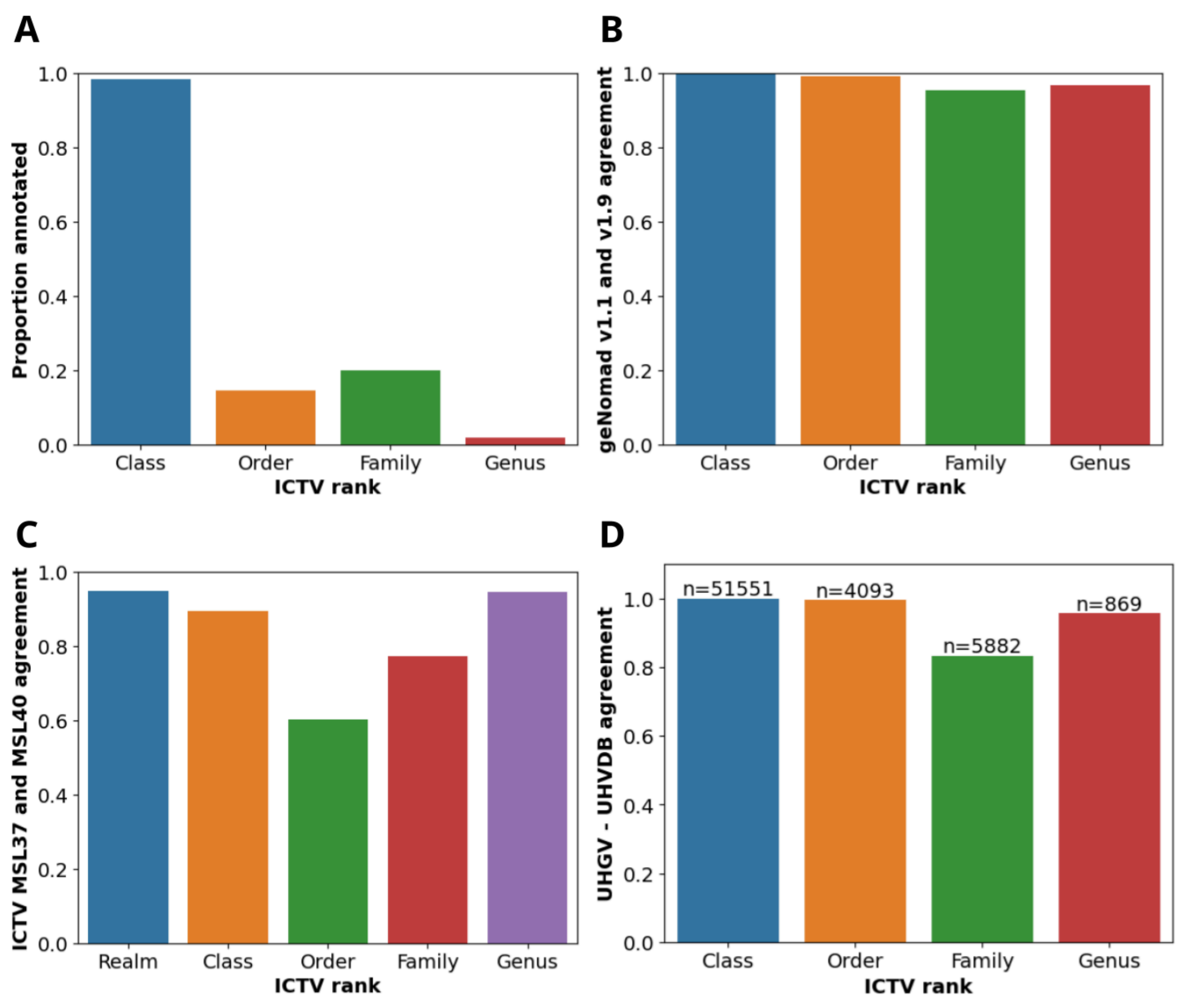


**Supplementary Figure S8.** Benchmarking UHVDB's taxonomic annotations. (**a**) Proportion of UHGV genomes receiving a taxonomic annotation at different ranks. (**b**) Agreement between the taxonomy assigned by geNomad v1.1 and geNomad v1.9 to UHGV sequences at different ranks. (**c**) Agreement between the taxonomy assigned to reference sequences present in both ICTV MSL 37 and ICTV MSL 40 at different taxonomic ranks. (**d**) Agreement between taxonomic assignments for UHGV sequences derived from UHGV's and UHVDB's taxonomy protocol at different ranks. The number of UHGV sequences annotated at each rank is depicted above each bar.

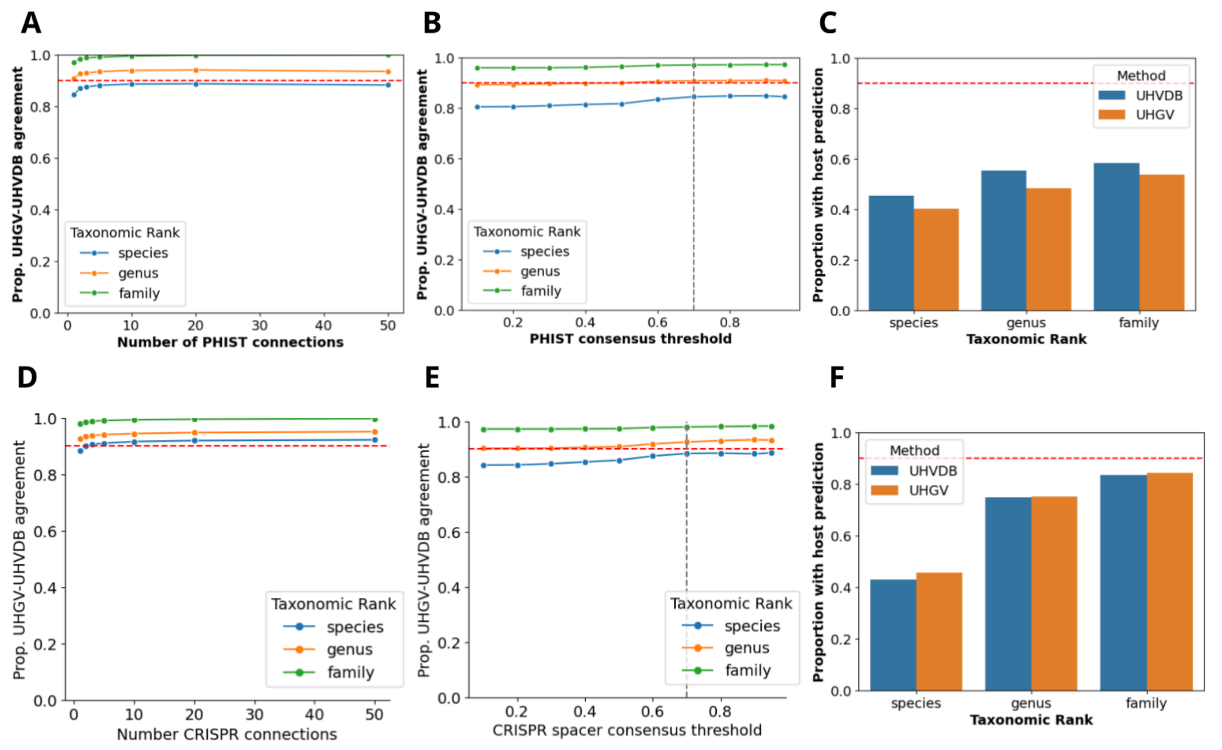


**Supplementary Figure S9.** UHVDB host prediction benchmarking. Agreement between UHGV and UHVDB release 1 (r1) PHIST host predictions on UHGV genomes as a function of (**a**) the minimum required percentage of phage k-mers contained in the host genome (**b**) the number of host genomes connected to each virus by PHIST. Agreement was calculated at the family, genus, and species levels. (**c**) Proportion of UHGV genomes having host predictions at the species, genus, and family levels with at least 1 PHIST connection and a consensus of ≥ 70% connections agreement. (**d**) Agreement between UHGV and UHVDB CRISPR host predictions on UHGV genomes as a function of the number of CRISPR connections. (**e**) Proportion of UHGV genomes having host predictions at the species, genus, and family levels with at least 1 CRISPR connection and a consensus of ≥ 70% connections agreement. (**f**) Proportion of UHGV genomes having a species, genus, or family-level prediction using either UHVDB's (blue) or UHGV's (orange) CRISPR spacer predictions.

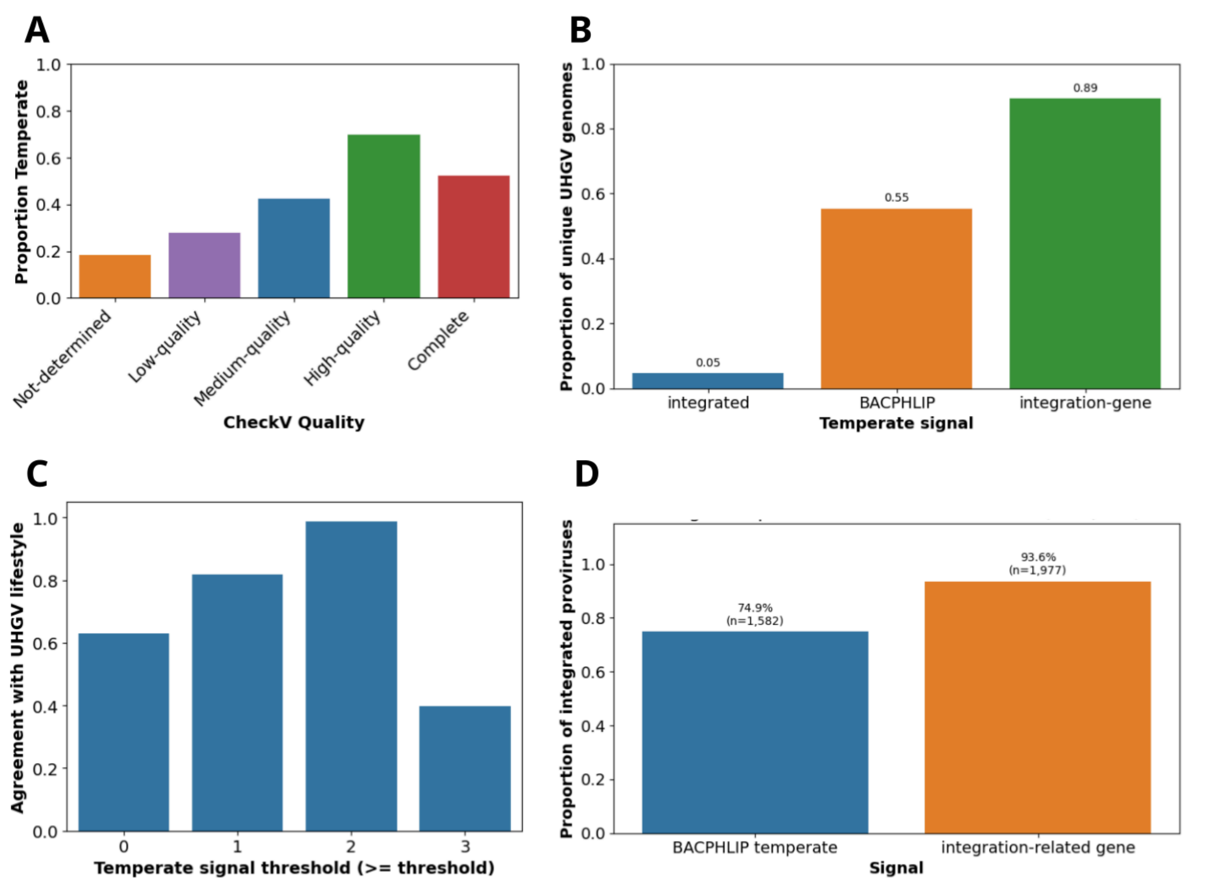


**Supplementary Figure S10.** Benchmarking lifestyle annotations. (**a**) Proportion of UHGV viruses classified as temperate by UHGV as a function of virus completeness. (**b**) Proportion of UHGV viruses having different temperate signals used by UHVDB. (**c**) Agreement between UHGV and UHVDB lifestyle classification based on the minimum number of temperate signals (see figure b) to classify a virus as temperate. (**d**) Proportion of UHGV integrated proviruses classified as temperate using a BACPHLIP score > 0.5 or by requiring the presence of a UHVDB-detected integration gene.

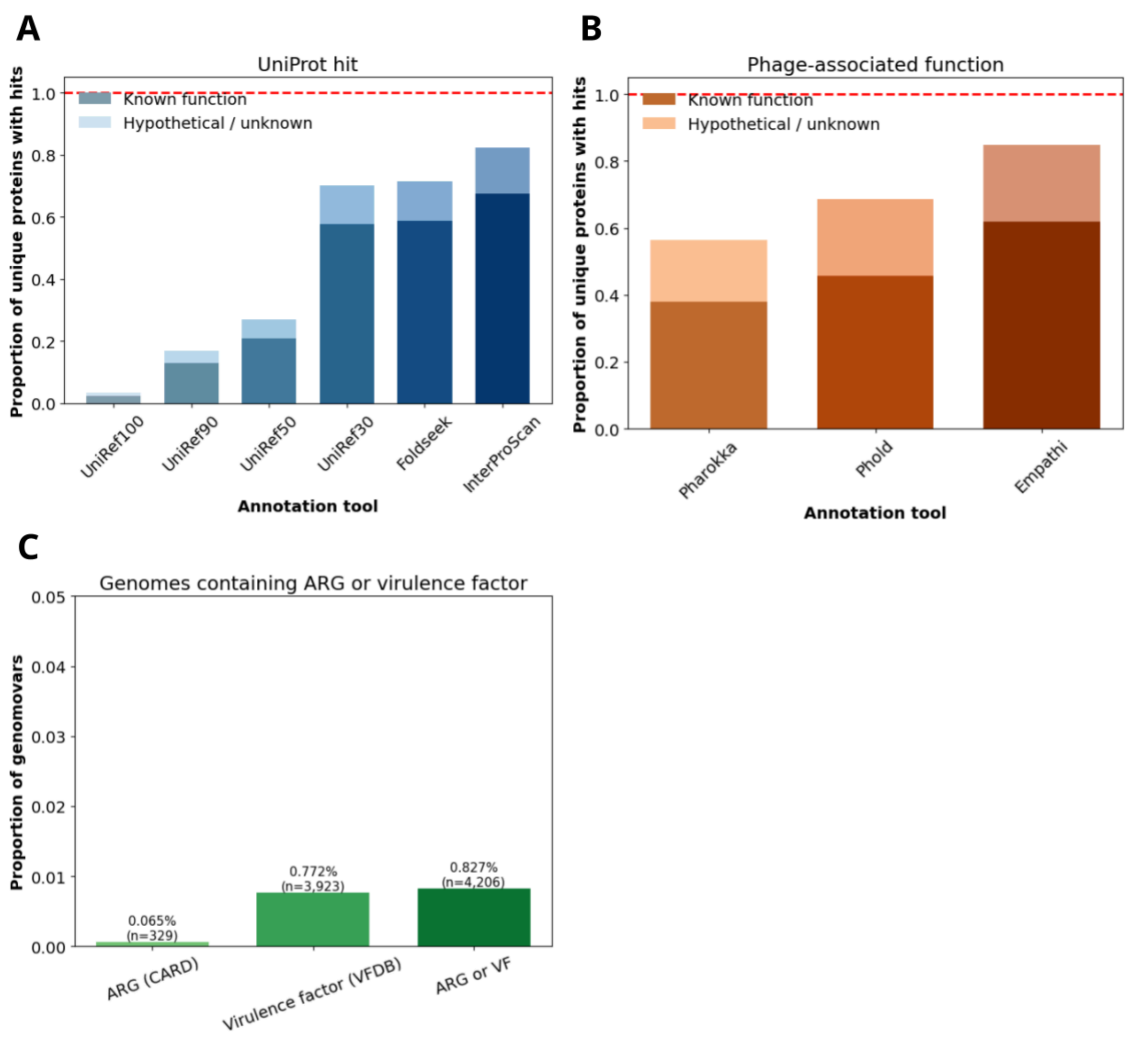


**Supplementary Figure S11.** Benchmarking functional annotations. Cumulative proportion of unique UHVDB release 4 (r4) proteins annotated with (**a**) UniProt/InterPro accessions and (**b**) phage-related functions after each search method was implemented. (**c**) Proportion of genomovar representatives with virulence factors (VFs) detected from the Virulence Factor Database and antibiotic resistance genes (ARGs) detected from the Comprehensive Antibiotic Resistance Database (CARD).

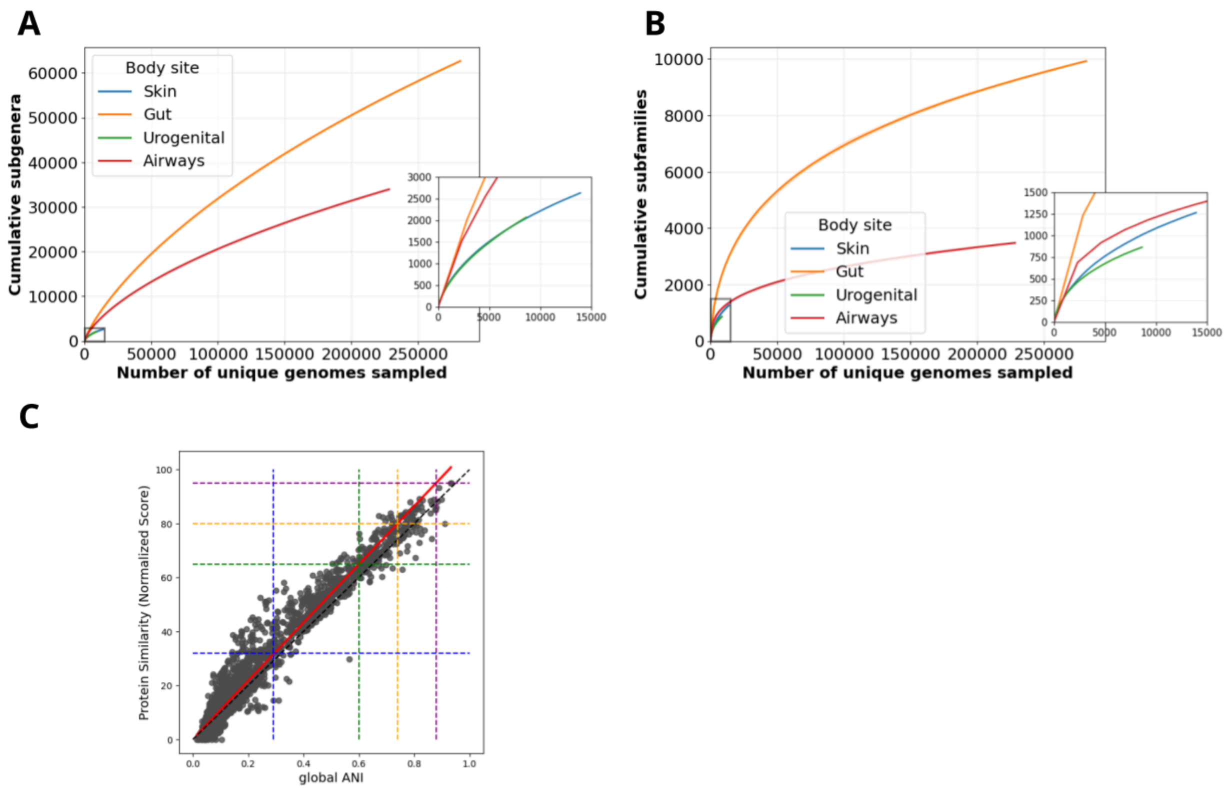


**Supplementary Figure S12.** Sub-taxa accumulation curves and relationship between protein similarity and global average nucleotide identity. (**a**) Subgenus (80% proteomic similarity) and (**b**) subfamily (32% proteomic similarity) accumulation curves. UHVDB release 4 (r4) unique sequences were split into 100 increasing subsets and analyzed to determine the number of clusters present at each interval. This process was repeated 50 times. Lines depict the mean species count at each interval and shaded regions represent the area between the minimum and maximum values. (**c**) Relationship between global average nucleotide identity (ANI) and proteomic similarity. 1,000 different UHGV species representatives were selected and run through all-v-all nucleotide alignments with vClust and proteomic similarity calculations using our pipeline. Then a regression (red line) was plotted to display the relationship between subfamily (blue), genus (green), subgenus (orange), and species (purple) thresholds when using global ANI versus proteomic similarity.

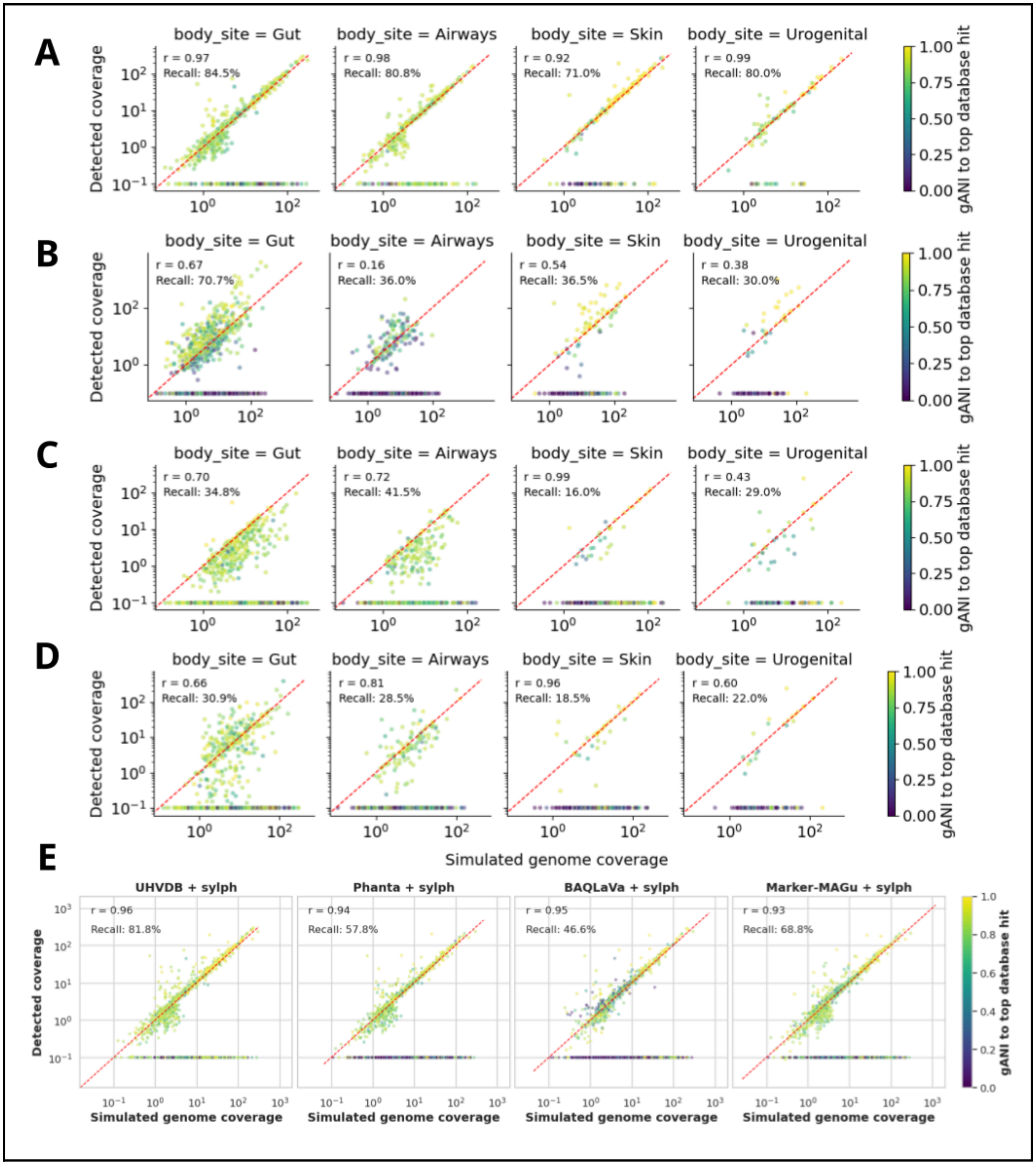
A
B
C
D
E
body_site = Gut
body_site = Airways
body_site = Skin
body_site = Urogenital
Detected coverage
Simulated genome coverage
gANI to top database hit
r = 0.97
Recall: 84.5%
r = 0.98
Recall: 80.8%
r = 0.92
Recall: 71.0%
r = 0.99
Recall: 80.0%
r = 0.67
Recall: 70.7%
r = 0.16
Recall: 36.0%
r = 0.54
Recall: 36.5%
r = 0.38
Recall: 30.0%
r = 0.70
Recall: 34.8%
r = 0.72
Recall: 41.5%
r = 0.99
Recall: 16.0%
r = 0.43
Recall: 29.0%
r = 0.66
Recall: 30.9%
r = 0.81
Recall: 28.5%
r = 0.96
Recall: 18.5%
r = 0.60
Recall: 22.0%
UHVDB + sylph
Phanta + sylph
BAQLaVa + sylph
Marker-MAGu + sylph
r = 0.96
Recall: 81.8%
r = 0.94
Recall: 57.8%
r = 0.95
Recall: 46.6%
r = 0.93
Recall: 68.8%

**Supplementary Figure S13.** Virome profiler benchmarks. Virome profiling results for (**a**) UHVDB toolkit (**b**) BAQLaVa (**c**) Marker-MAGu and (**d**) Phanta split by body site. To benchmark different virus profilers, we used the high-quality, confident viruses that were assembled as part of our comprehensivity analysis. These viruses were clustered at the species level (95% ANI and 85% AF) and split into 17 sets of 100 viruses, each containing viruses exclusively from one body site. For each set, 100,000 reads were simulated using InSilicoSeq. Then BAQLaVa, Phanta, Marker-MAGu, and the UHVDB toolkit were run. Input viruses were aligned to the virus sequences used by each profiler with BLAST, and reciprocal-best-hits were identified using global ANI (ANI x AF). For each body site, recall (number of detected viruses divided by the total number of input viruses) and Pearson R values were calculated. Here, each point represents an input genome, point color denotes the global ANI between the input and most similar database genome, and the red dashed line indicates a perfect 1:1 correlation. (**e**) Sylph was run using each tool's database of virus genomes, and performance was evaluated on these outputs across all body sites as described above.

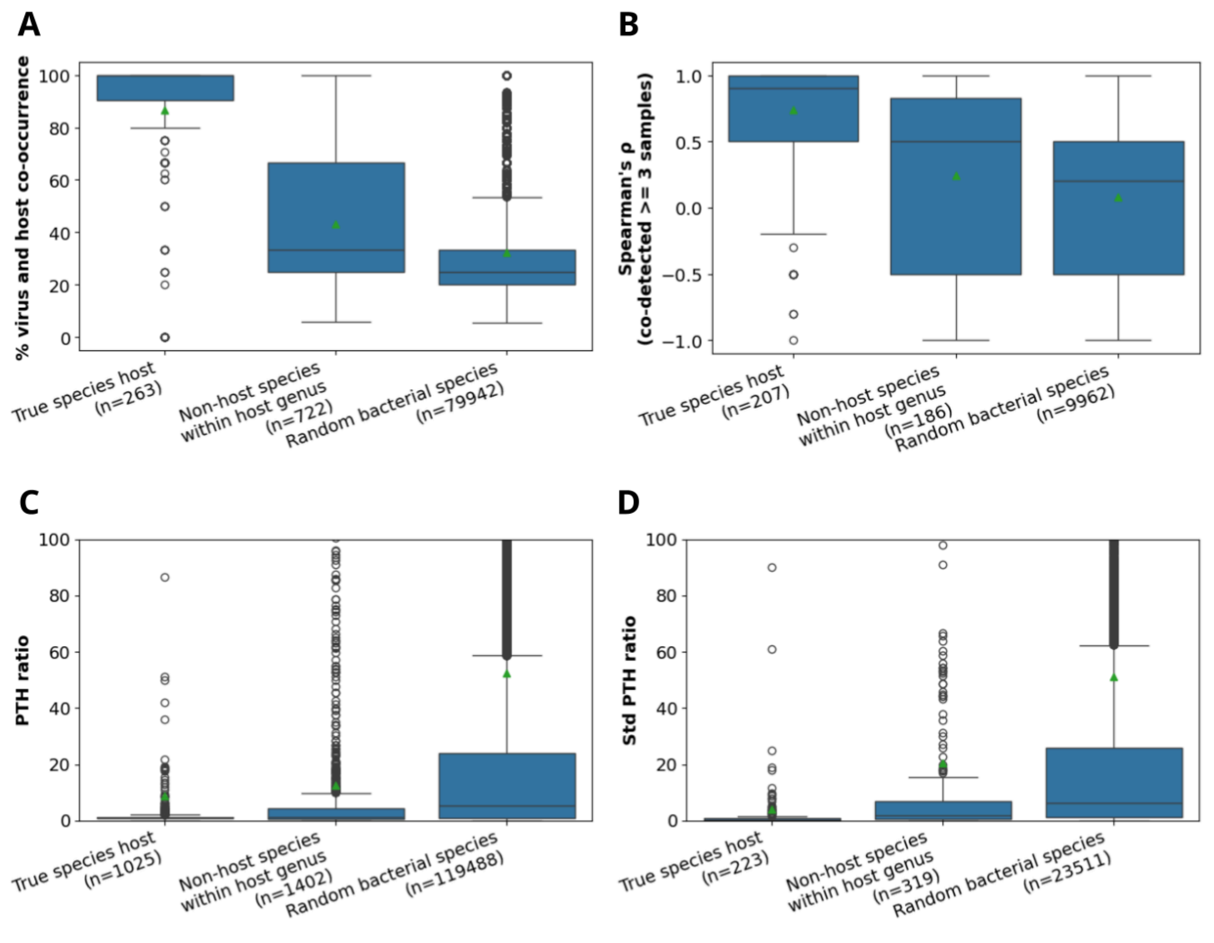

**Supplementary Figure S14.** Analyzing phage host species predictions. The analyze workflow from the UHVDB toolkit was run on 39 bulk metagenomes from infant fecal samples. For all viruses detected in ≥ 3 samples, (**a**) the percentage of samples where viruses co-occured with their predicted bacterial species host ("True Species Host"), another bacterial species within the same genus as their predicted host ("Non-host species within host genus"), and a random bacterial species ("Random Host Species") was evaluated. (**b**) Spearman correlation was calculated for virus-host pairs co-occurring in ≥ 3 samples. (**c**) Phage-to-host (PTH) ratios (phage depth of coverage divided by bacterial species coverage) was calculated within each sample for each phage-host combination described in S14A. (**d**) For phage-host pairs co-occurring in ≥ 2 samples, the standard deviation (STD) in PTH ratios was determined.

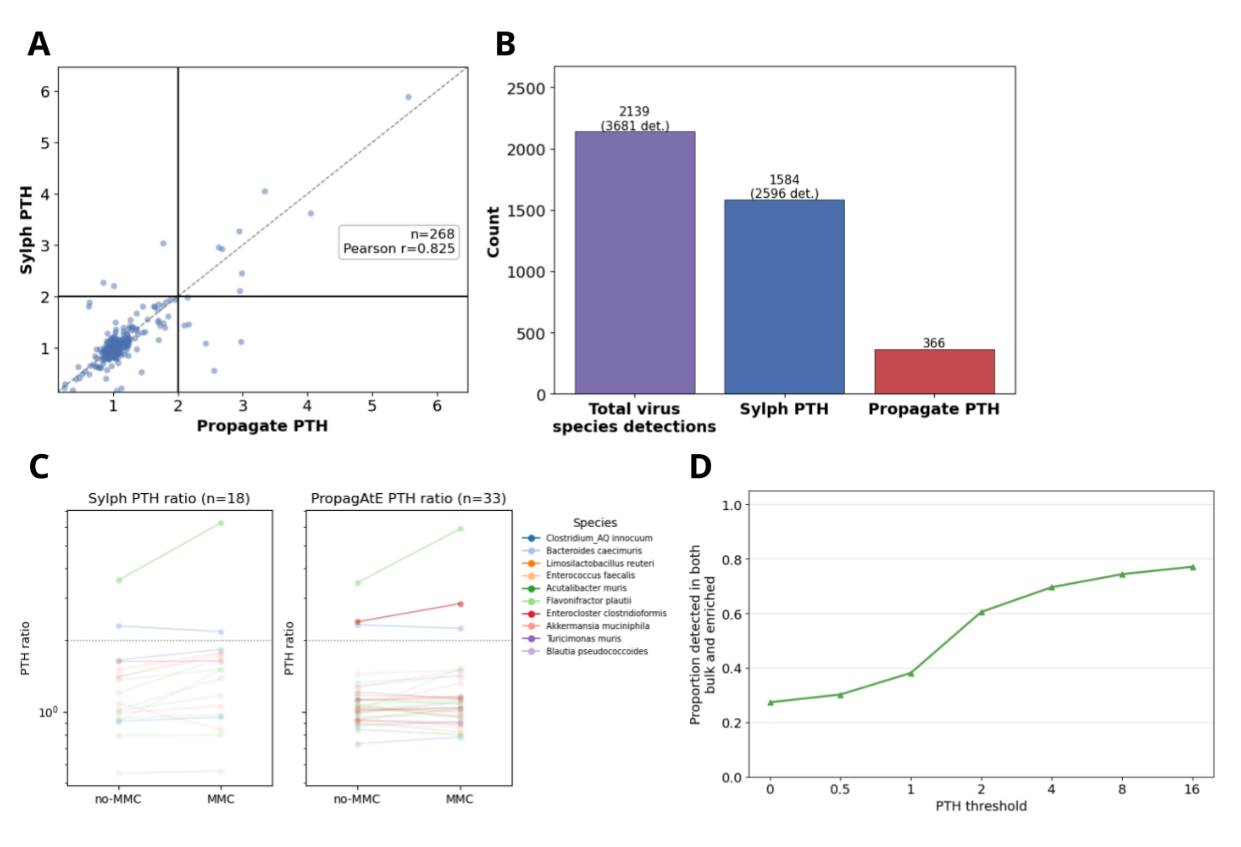


**Supplementary Figure S15.** Benchmarking phage-to-host (PTH) ratios. (**a**) PTH ratios were calculated using sylph and PropagAte on a set of fecal metagenomes (94), and PTH ratios output by each tool were compared by identifying reciprocal best hits between the *de novo* assemblies used by PropagAte and UHVDB species representatives. (**b**) The number of total virus species detected in these metagenomes is compared to the number of PTH ratios calculated using sylph and PropagAte. Top numbers above the bars indicate the number of unique species detected, while the bottom number indicates the total number of virus species detections. (**c**) A set of short reads from a shotgun metagenome of a mock bacterial community treated with and without mitomycin C (MMC).(Zünd et al. 2021) PTH ratios were calculated with and without MMC, with the same virus species connected by a line across conditions. The count at the top indicates the number of species detected by each method (complete circular reference assemblies were used as input for PropagAte) and the dashed horizontal line indicates a PTH of 2. (**d**) Sylph PTH ratios were calculated from paired unenriched (bulk) and enriched (for virus-like particles) fecal metagenomes. The proportion of UHVDB species detected in both the bulk and enriched metagenomes is depicted as a function of increasing minimum PTH thresholds, indicating likely phage particle production.

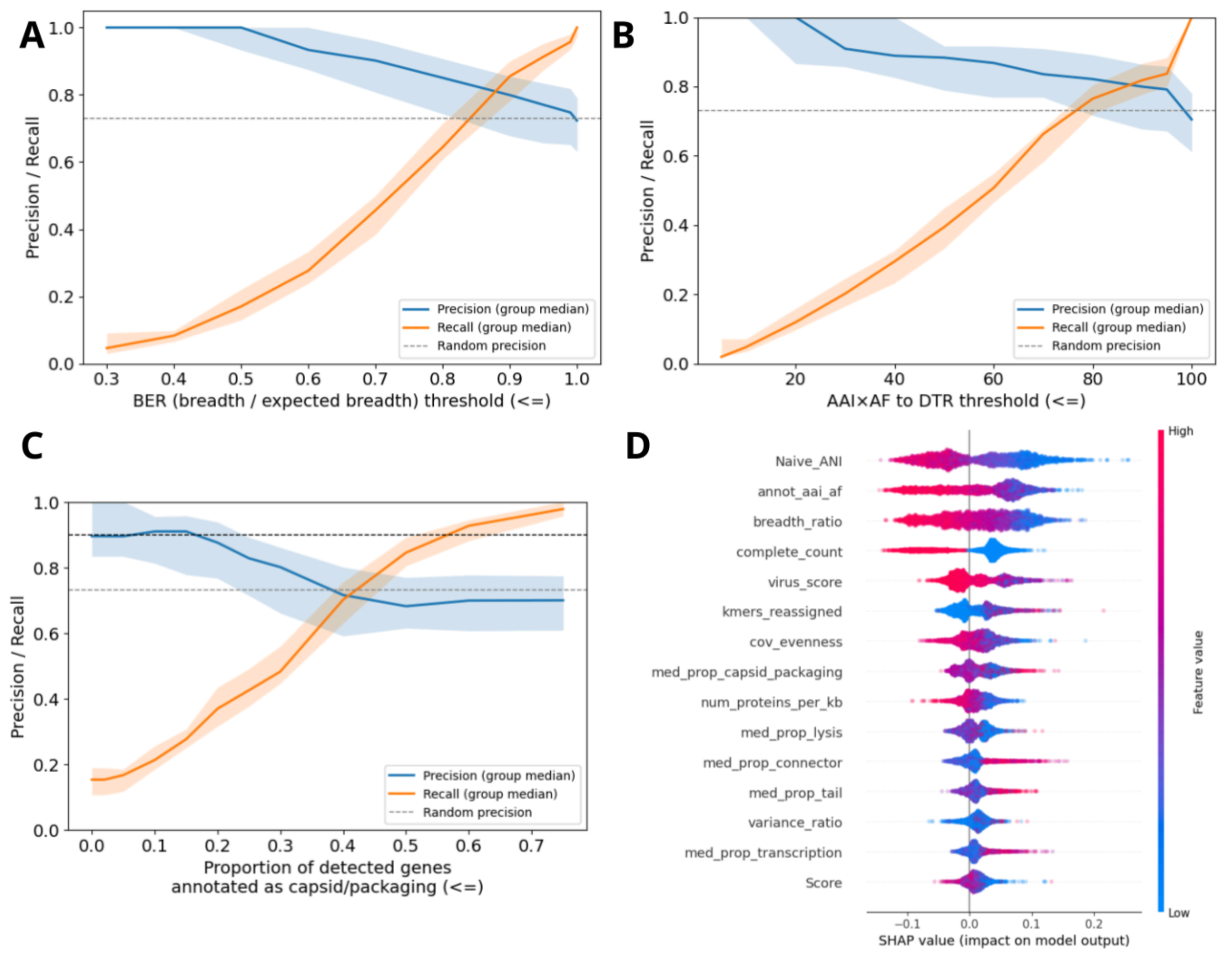


**Supplementary Figure S16.** Evaluating signals of uninducible prophages. Paired unenriched (bulk) and enriched fecal metagenomes were analyzed to determine if certain features were associated with detection exclusively in the bulk sample. Precision (proportion of viruses passing a threshold that were exclusively detected in the bulk sample) and recall (proportion of all viruses detected exclusively in bulk samples that also passed a threshold) were evaluated for (**a**) breadth-expected-breadth (BER) ratio (**b**)

AAI x AF to a direct terminal repeat (DTR) containing genome (**c**) proportion of virus genes having a breadth of coverage > 80% that were annotated as capsid/packaging. Performance was evaluated per sample. Shaded regions indicate the interquartile range with lines indicating the median. (**d**) Shapley additive explanations (SHAP) values for the uninducible prophage random forest classifier. Each point is one genome, point color indicates the feature value (blue=low value; red=high value), and position on the x-axis (SHAP value) indicates the impact on the model's uninducible probability (right=more likely to be uninducible; left=less likely to be uninducible).

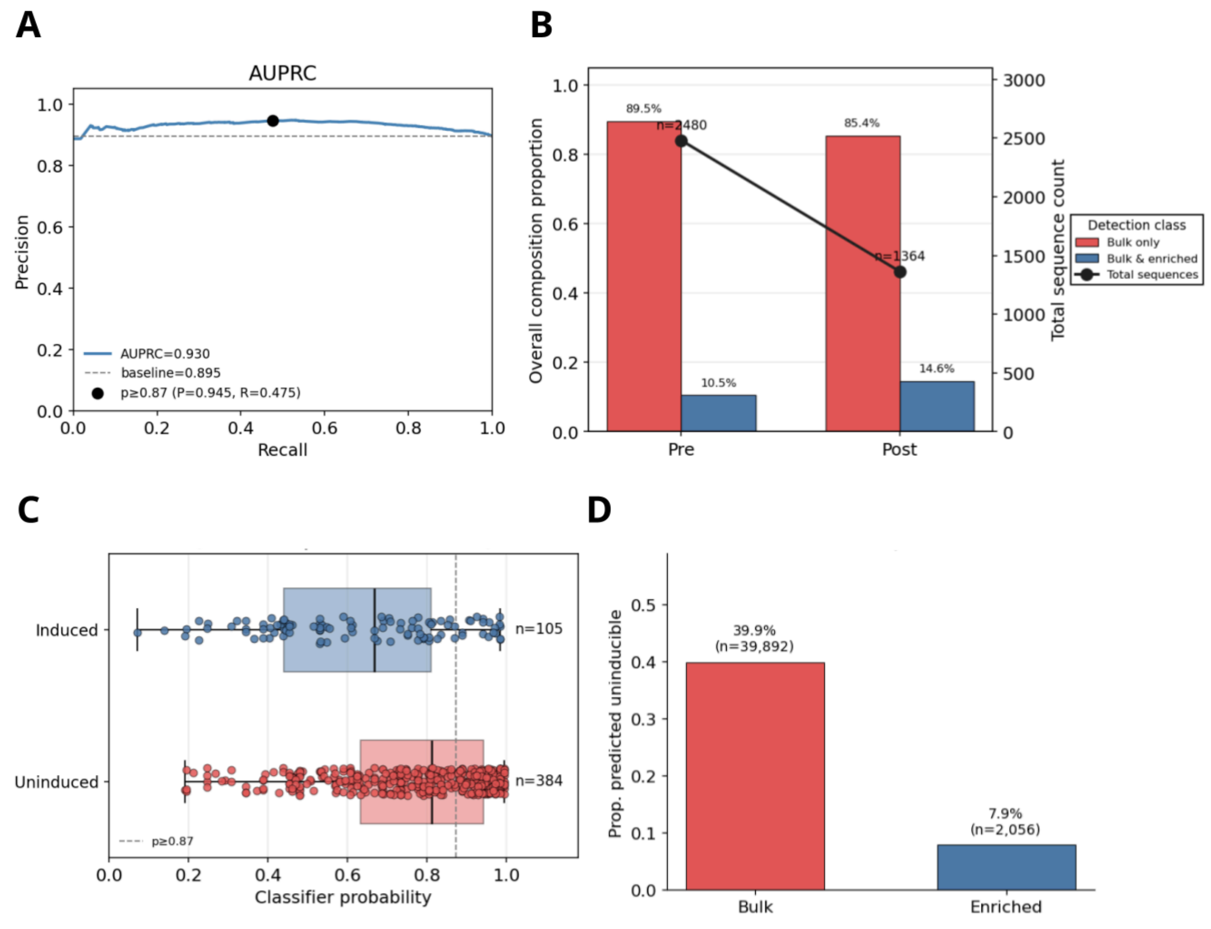


**Supplementary Figure S17.** Evaluating a random forest classifier for predicting uninducible prophages. (**a**) Precision and Recall when applying the classifier to a validation dataset of paired bulk and enriched fecal metagenomes. (**b**) The proportion of viruses that were detected in bulk only or both bulk and enriched samples before (Pre) and after (Post) removing viruses classified as uninducible prophages. The line indicates the total number of virus species pre and post filtering. (**c**) The uninducible prophage classifier was run on 187 isolates having paired mitomycin C (MMC) and no MMC samples. The classifier probability of being uninducible was split based on whether or not a virus had a ≥ 2x increase (induced) in PTH in MMC sequences compared to no MMC sequences. Counts are the number of virus species detected in this isolate dataset. (**d**) The proportion of bulk metagenome viruses and enriched metagenome viruses labeled as uninducible by our classifier.

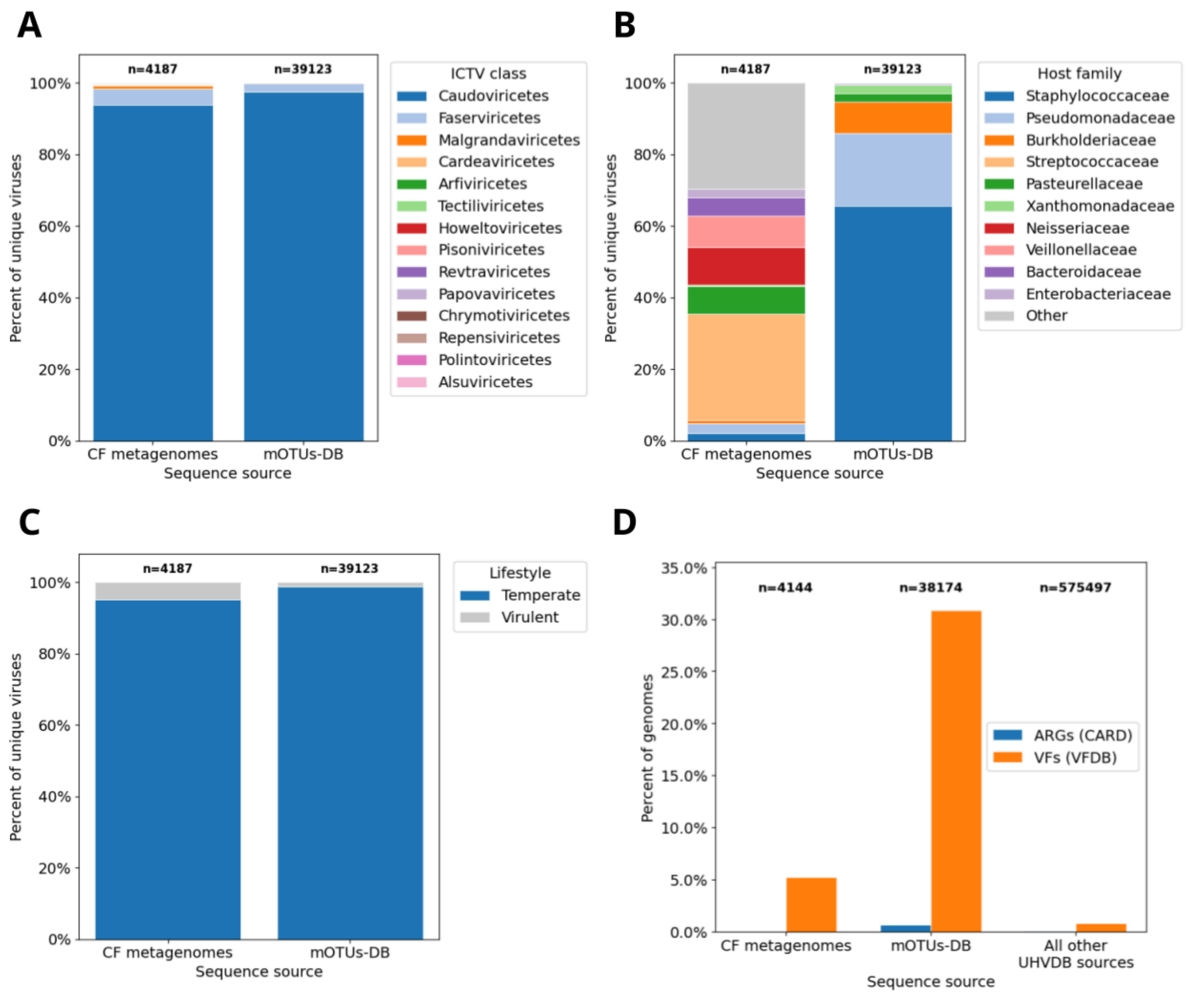


**Supplementary Figure S18.** Characteristics of viruses added to UHVDB for r5. Newly-added UHVDB r5 viruses were split based on whether they were derived from CF metagenomes or from prophages in a bacterial genome database (mOTUs-db) (18). Then, we compared (**a**) ICTV taxonomic annotations (**b**) Host family predictions (**c**) lifestyle predictions and (**d**) the percent of virus genomes with detectable antibiotic resistance genes (ARGs) and virulence factors (VFs) compared to all other UHVDB viruses.

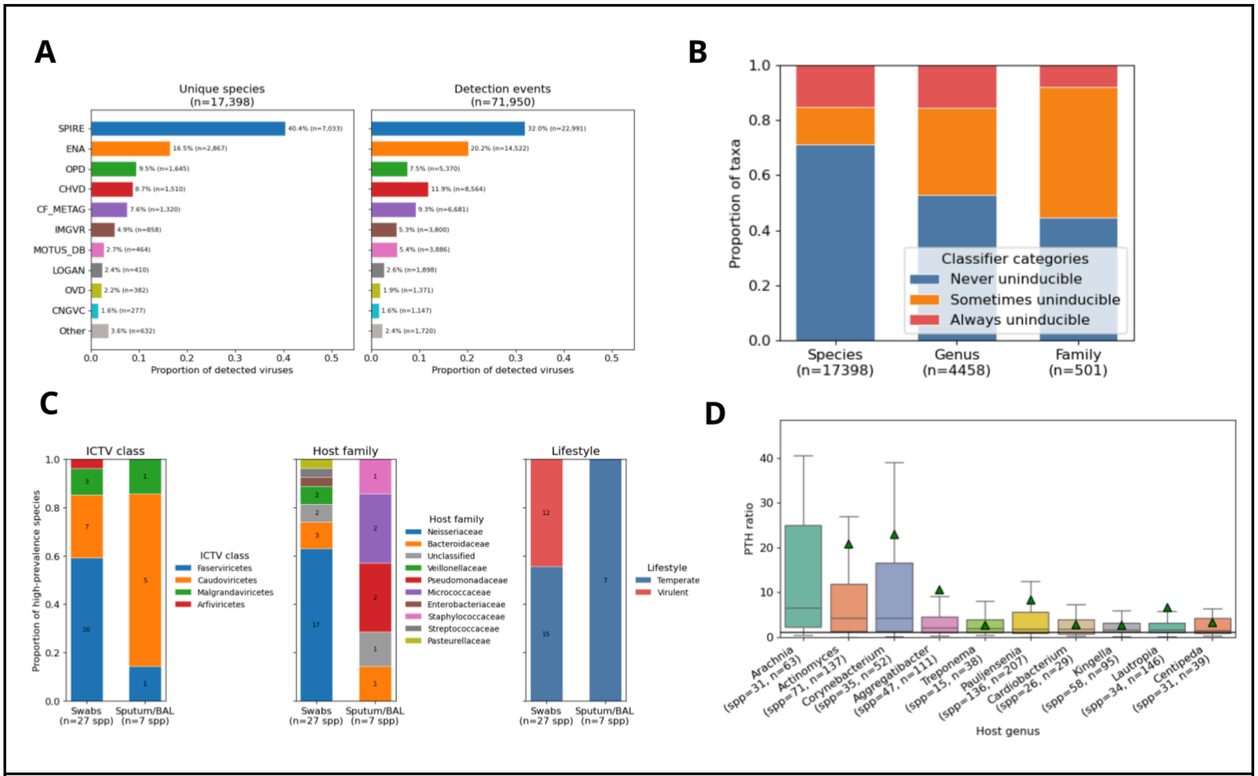


**Supplementary Figure S19.** CF respiratory tract tract sample virome analysis. (**a**) Proportion of unique species (left) and total detected viruses (right) in CF respiratory metagenomes that were mined from different source databases (CF_METAG=CF metagenome assemblies, MOTUS_DB=CF conventional pathogen assemblies). (**b**) The proportion of detected virus species, genera, and families that were never classified as inducible, sometimes classified as inducible, or always classified as inducible across all CF respiratory metagenomes. (**c**) The ICTV class (left), host family (center), and lifestyle (right) assignments of viruses detected in ≥ 10% of swab or sputum/BAL samples. (**d**) Phage-to-host (PTH) ratios for the 10 bacterial genera displaying the highest median PTH values across all phage-bacteria pairs detected in CF respiratory tract sample metagenomes.

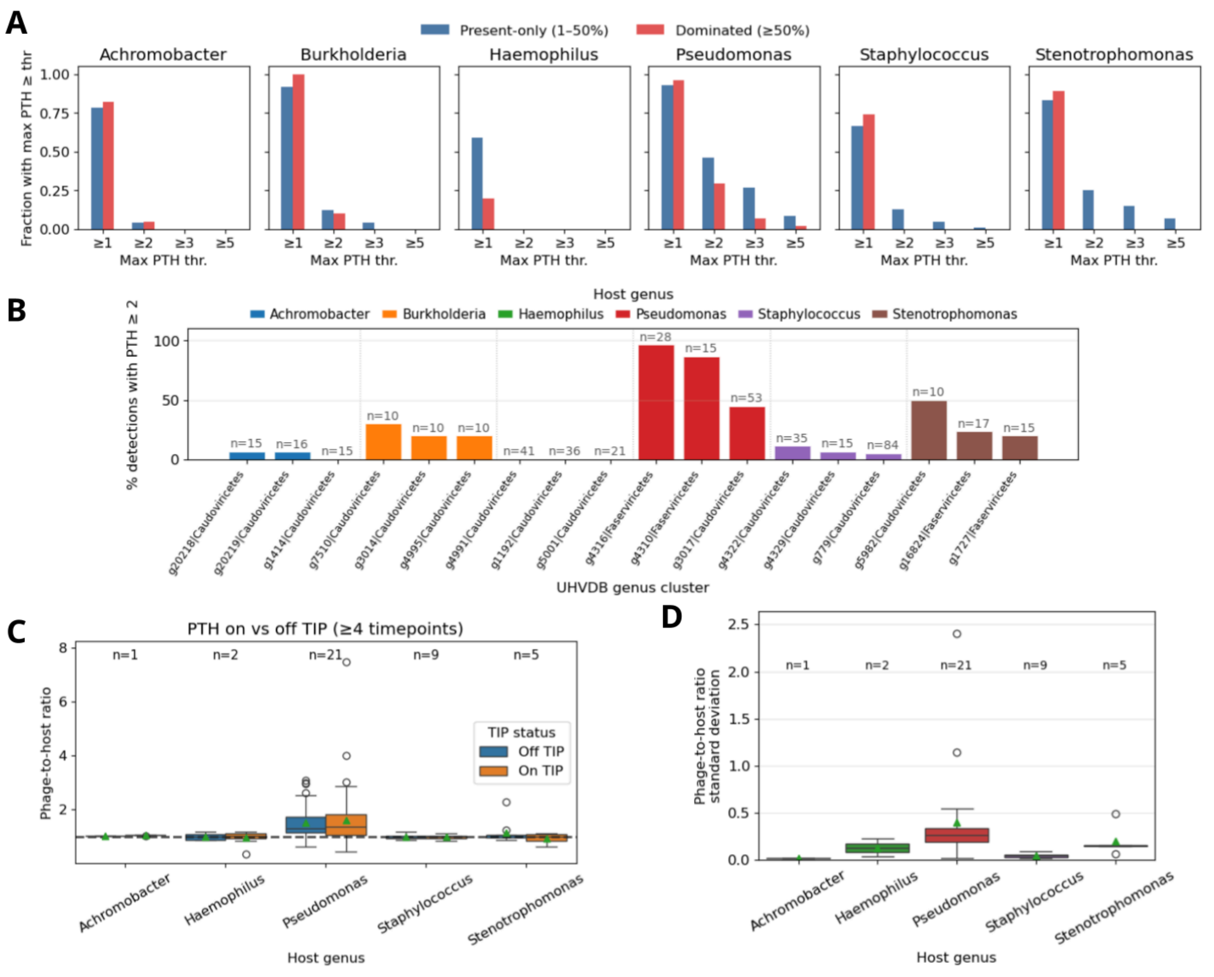


**Supplementary Figure S20.** Dynamics of phages infecting conventional CF pathogens. (**a**) Proportion of conventional CF pathogens having phages with phage-to-host (PTH) ratios ≥ the threshold indicated on the x-axis. Bars are split based on whether the pathogen was present only (1-50% relative abundance) or dominant (≥ 50% relative abundance) in a sample. (**b**) Percentage of UHVDB genera detections having PTH ratios ≥ 2, split by host genus. The UHVDB genus cluster identified and ICTV class are indicated on the x-axis. (**c**) PTH ratios for each detected virus infecting a conventional CF pathogen split by whether the sample was collected on or off the antibiotic tobramycin inhaled powder (TIP). (**d**) Standard deviation in PTH ratios across all timepoints for each virus virus species displayed in C.

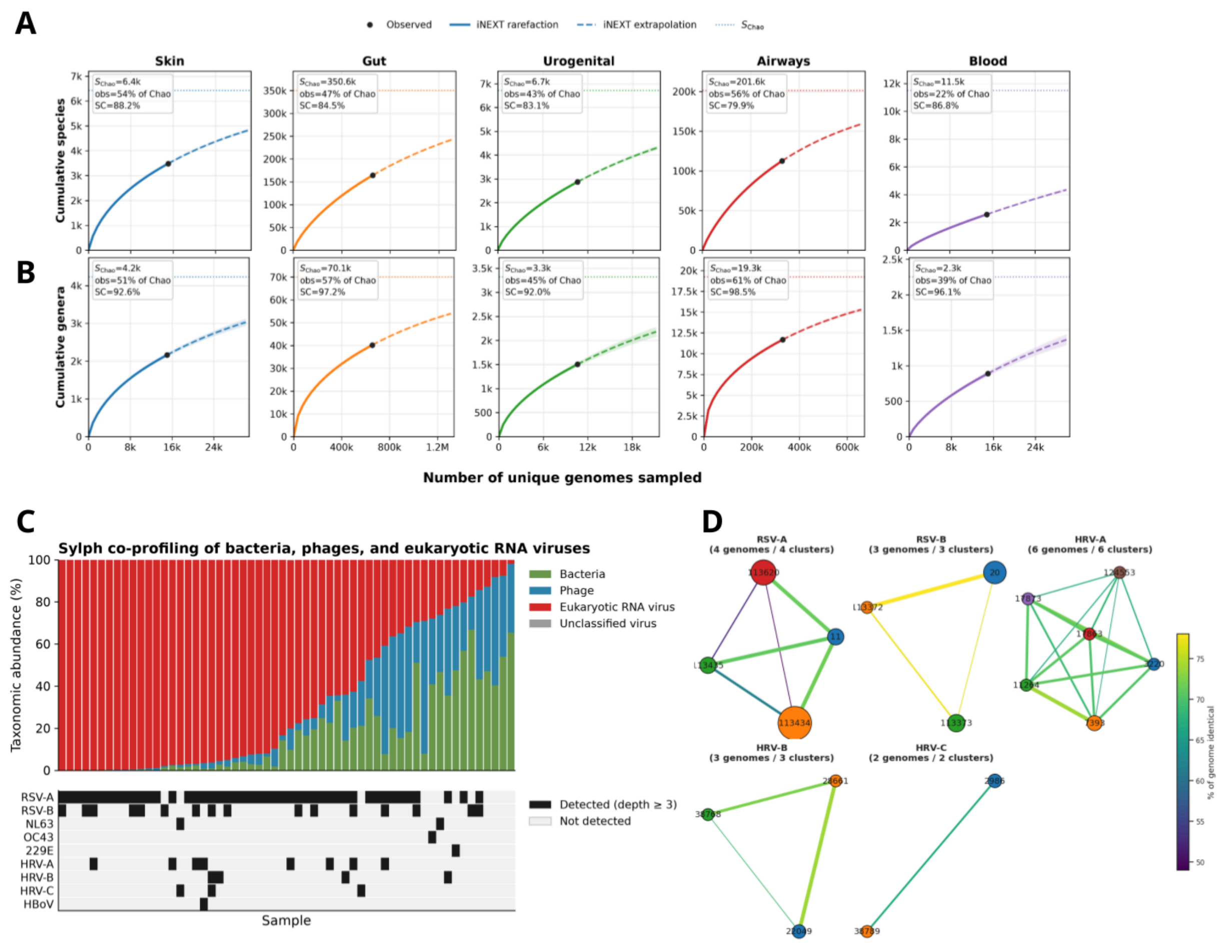


**Supplementary Figure S21.** Characteristics of UHVDB r6. Extrapolations of the (**a**) species and (**b**) genus accumulation curves for UHVDB r6 split by body site. $S_{Chao}$ is the total number of taxa predicted to be in each body site (plateau), obs=the percentage of total predicted taxa that are currently in UHVDB, SC=sample coverage, or the percentage of newly added unique viruses that are expected to represent a new taxa. (**c**) Taxonomic abundance of bacteria, phage, and eukaryotic viruses in nasopharyngeal metatranscriptomes (top). A mosaic plot indicating which human respiratory virus was detected in the sample depicted in the stacked bar chart (bottom). (**d**) Network plot displaying the genomic distance (percentage of the genomes that are identical) between UHVDB species representatives classified as RSV-A, RSV-B, HRV-A, HRV-B, and HRV-C. (RSV=respiratory syncytial virus, HRV=human rhinovirus).

### Supplementary Tables

**Table S1.** List and description of features used to train and evaluate the random forest classifier for identifying uninducible prophages.

| Feature | Description |
|---|---|
| **Genome truncation** | |
| breadth_ratio | Bulk CoverM observed genome breadth divided by expected breadth 1-exp(-0.833 * mean coverage). |
| annot_aai_af | CheckV AAI alignment fraction (aai_af) of the species-rep genome versus the closest complete reference. |
| Naive_ANI | sylph profile Naive_ANI of the genome in the bulk sample. (kmer containment) |
| **Direct terminal repeat presence** | |
| complete_count | 1 if the species-rep CheckV quality is Complete, else 0 |
| **Spurious detections** | |
| variance_ratio | Bulk CoverM coverage variance divided by trimmed_mean when trimmed_mean > 0. |
| cov_evenness | sylph Median_cov / Mean_cov_geq1 in the bulk sample (evenness of coverage among positions with coverage ≥ 1). |
| kmers_reassigned | sylph kmers_reassigned count for the genome in the bulk sample. |
| **Viral confidence** | |
| Score | viralVerify Score of the species-rep genome. |
| viral_gene_frac | CheckV viral_genes / (viral_genes + host_genes) on the species-rep genome. |
| virus_score | geNomad virus_score of the species-rep genome. |
| num_proteins_per_kb | Species-rep protein count (num_proteins) divided by contig length in kb. |
| **Integration/Temperate evidence** | |
| is_integrated | 1 if the species-rep integration_status is "integrated", else 0. |
| temperate | BACPHLIP temperate score of the species-rep genome. |
| empathi_integration_per_kb | Empathi integration-gene count on the species-rep divided by contig length in kb. |
| **Virus structural gene coverage** | |
| med_prop_capsid_packaging | Fraction of genes with CoverM breadth > 0.8 annotated as capsid/packaging (Pharokka, Phold, or Empathi). |
| med_prop_tail | Fraction of genes with CoverM breadth > 0.8 annotated as tail (Pharokka, Phold, or Empathi). |

| | |
|---|---|
| med_prop_lysis | Fraction of genes with CoverM breadth > 0.8 annotated as lysis (Pharokka, Phold, or Empathi). |
| med_prop_connector | Fraction of genes with CoverM breadth > 0.8 annotated as connector / head–tail joining (Pharokka or Phold). |
| med_prop_transcription | Fraction of genes with CoverM breadth > 0.8 annotated as transcription regulation (Pharokka or Empathi). |
| n_struct_genes | Count of major capsid (MCP), terminase large subunit (TerL), and portal genes on the genome in the bulk gene-coverage table (Pharokka, Phold, or Empathi). |
